# Unraveling viral identity: Avoiding the trap of endogenous sequences for viral surveillance of small ruminant oncogenic retroviruses

**DOI:** 10.64898/2026.03.05.709768

**Authors:** Riocreux-Verney Benjamin, Verneret Marie, Dolmazon Christine, Shirin Ashraf, Stella A. Atim, Navratil Vincent, Leroux Caroline, Turpin Jocelyn

## Abstract

Small ruminants (sheep and goats) are one of the few mammals in which an exogenous retrovirus (XRV) and closely related endogenous retroviral elements (ERV) coexist within the same host genome. The betaretroviruses Jaagsiekte sheep retrovirus (JSRV) and enzootic nasal tumor virus (ENTV) cause pulmonary and nasal adenocarcinomas, respectively, and share extensive sequence similarity with their endogenous counterparts. Consequently, molecular surveillance must rely on assays that can unequivocally distinguish true exogenous infection from ERV-derived templates; failure to do so compromises diagnosis, phylogenetic inference, and epidemiological conclusions.

We retrieved all complete JSRV, ENTV-1/2, and related ERV genomes deposited in public repositories and performed a comprehensive alignment. Only a limited number of genomic segments were capable of distinguishing exogenous from endogenous sequences. We refer to these as discriminating regions (DRs). Phylogenies built using DRs revealed that several entries annotated as XRV are, in fact, ERV-derived or chimeric artefacts generated by short-amplicon reconstruction.

A systematic literature review of over 100 articles identified 286 distinct primers and probes used for the XRV amplification. *In-silico* mapping of each oligonucleotide onto the full alignment showed that only 28 % reliably differentiate XRV from ERV. We experimentally validated the predictive power of this approach for 17 primer/probe sets, confirming that non-discriminating assays produce false-positive signals from endogenous templates.

The misannotation of ERV sequences as exogenous viruses has resulting in the population of databases with dubious entries, fostering erroneous hypotheses such as vector-borne transmission of JSRV and ENTV. To address this issue, we propose a concise set of criteria for assay design, validation, and database annotation emphasizing DR targeting, specificity testing against endogenous templates, and transparent reporting.

Although this framework was developed for small ruminants, it is readily applicable to any host–virus system in which exogenous viruses coexist with endogenous viral elements. This will strengthen viral surveillance, phylogenetics, and the One Health initiatives.

## Introduction

Cancers induced by the exogenous retroviruses (XRVs), Jaagsiekte Sheep RetroVirus (JSRV) and Enzootic Nasal Tumor Virus (ENTV), have been documented in all regions where sheep and goats are reared, except in Australia, New Zealand, the Falkland Islands and Iceland, where JSRV has been eradicated [1–3]. Multiple copies of endogenous retrovirus (ERV) family II.5, previously named enJSRV and enENTV [4,5], are present in goats (ERV II.5 Ch for *Capra hircus*) and sheep (ERV II.5 Oa for *Ovis aries*), with an average of 223 and 98 loci respectively [6]. The presence of whole copies with little divergence and with preserved open reading frames (*gag, pro, pol* and *env*), suggests the recent activity of ERV II.5 Ch in goats [6]. Despite their close genetic relationship with domestic sheep and goats XRVs, ERVs are not implicated in the development of respiratory cancers in domestic small ruminants. Therefore, the exogenous retroviruses JSRV and ENTV remain the only known aetiological agents of respiratory cancers in domestic small ruminants.

Developing robust virus detection tools is an essential for managing the infectious risk posed by these incurable, fatal and transmissible cancers that threaten small-ruminant farming. Serological detection of antibodies is a tool widely used in both veterinary and human medicine. Apart from the fact that it only allows indirect detection of infection through the demonstration of antibody production, it requires a detectable and stable humoral response. In a context of multiple ERV copies impairing the development of a XRV-specific humoral response, antibody detection in JSRV or ENTV infected animals is not a reliable tool to establish the infectious status [7–10].

Therefore infection can only be detected *ante or post mortem* using molecular approaches, *i.e.* PCR and/or RT-PCR [7,11]. The genetic diversity of JSRV and ENTV is only partially known, and before our recent study reporting on 29 JSRV and 24 ENTV partial and full proviral sequences [12], only a few dozen complete sequences were available in the databases, i.e. nine complete sequences of JSRV isolated in Africa, North America, Europe and Asia [13–17], and 50 complete or near-full length ENTV genomes isolated in Canada and Europe for ENTV-1 [18,19] and in Europe and Asia for ENTV-2 [20–26], including sequences not associated with accessible publications or unpublished.

The widespread use of high-throughput sequencing tools has made pathogen sequences widely available. In theory, detecting viral or proviral genomes using targeted molecular approaches is straightforward, given the multiple bioinformatic tools available for designing primers and probes. However, when dealing with the coexistence of genetically closely related ERV family II.5 and XRVs, it is crucial to ensure the strict XRV specificity of the primers or probes. To optimize the molecular XRV detection tools in regards to published studies, we reviewed multiple reports, focusing on the molecular detection and sequencing tools used and their relevance in the context of the coexistence of XRVs with ERV II.5 *loci* [6]. A recent study of all available sequences in databases allowed us to identified sequences wrongly identified as exogenous JSRV or ENTV despite the absence of the YXXM motif in the region of the *env* gene encoding the intracytoplasmic tail (CT), suggesting their endogenous origin [12]. The YXXM motif is an essential oncogenic determinant, and a genetic marker of exogenous sequences, that is strictly absent from all the ERV II.5 Oa and Ch sequences identified up to date [6,12,18,27]. On the other hand, some studies have relied on RNAs to specifically detect XRVs [20,23,28–31]. This approach assumes that ERVs are not expressed in tissues, which is risky as we will show in this study.

In a large retrospective study based on published and unpublished sequences included in the public archives, we questioned the truly exogenous nature of the detected and sequenced viral genomes, after highlighting errors in the annotation of sequences unduly identified as XRV. To this end, we defined the regions of the JSRV and ENTV genomes specific to XRVs. We then reanalysed the available sequences and published amplification tools to determine their specificity, verified the XRV specificity of primers reported in the literature, and proposed a systematic approach with rigorous controls to ensure the exogenous nature of the sequences. We apply our methodology to case studies reporting on the detection of JSRV or ENTV in ticks and mosquitos [32–37] using metagenomics or virome approaches and their potential transmission by those vectors.

Our study raises questions about the robustness of the virus discovery in the context of XRVs/ERVs coexistence in small ruminants and the overall challenge in metagenomic and transcriptomic studies in the presence of ERVs and more globally endogenous viral elements (EVEs) [38].

## Materials and Methods

### Determination of discriminating regions between XRVs et ERVs

Complete nucleic sequences from 30 JSRV, 14 ENTV-1 and 35 ENTV-2 sequences available in GenBank (S1 Table, sequences identified with an asterisk *) were aligned with the ERV II.5 consensus for *Ovis aries* for JSRV and ENTV-1, and for *Capra hircus* for ENTV-2 [6]. Nucleotide alignments were performed with the ‘MAFFT v7.490’ software [39] and processed with the ‘R v4.4.1’ suite to picture the variability of the identity scores obtained from the comparison between XRV sequences and the ERV consensus and to calculate the percentage of identity for each position in the alignment, averaged over a 100 bp window with a 10 bp step. The analysis was performed individually for each sequence of the alignment, and the mean percentage of identity was plotted. A 95% confidence interval was estimated using a bootstrap approach with the ‘boot v1.3-31’ suite [40], debiased and corrected for asymmetric data. The limited number of samples used and the potential lack of independence between sequences originated the same origins (herd, geographical location, etc.) are potential biases that we have identified. Four discriminant regions (referred further as DR1, DR2, DR3, DR4), were identified with a percentage of identity lower than 80%. DR2 sequence variability was evaluated by aligning (MAFFT Alignment v7.490 [39]) copies containing the *gag* region, *i.e.* 28 ERV II.5 copies annotated from the *Ovis aries* assemby (ARS-UI_Ramb_v2.0 -GCA_016772045.1) and 155 ERV II.5 copies from the *Capra hircus* assembly (ARS1.2 -GCA_001704415.2), and 27 ovine and 10 caprine ERV sequences recovered from GenBank [6] (S1 Table). The percentage of each nucleotide among all copies was calculated at each position.

### *in silico* and experimental evaluation of XRV primers specificity

The localization prediction of 286 published PCR primers and probes (S2 Table) reported for the detection of JSRV et ENTVs, was determined by mapping them onto the respective alignments for each virus. Publications not written in English were automatically translated with DeepL. A maximum of 4 mismatches from the XRV sequences was allowed, and the position of each primer was extracted from the alignment retaining the multi-mapped positions. Primers including additional sequences, *e.g.* restriction sites, were not included. Primers only mapping on XRV were defined as XRV specific whereas primers mapping on a least one copy of ERV, were defined as non-specific to exogenous sequences. A selection of seventeen primer pairs has been tested on various DNA templates, covering different parts of the viral genomes. Fourteen of them were tested by PCR on mammalian DNAs; they targeted ENTV-2 for primer pairs # 01-10 or JSRV for primers pairs 12-15 (S3 Table). Their ability to detect only XRV was tested using genomic DNAs from JSRV-induced lung tumours of clades I (FR2054) and II (FR989), and ENTV-2-induced nasal tumours (FR3824) [12] (Table 1). To evaluate their potential for cross hybridization with ERV sequences, PCRs were performed using DNAs from ovine IDO5 cells (Institut Mérieux, Lyon, France), caprine TiGEF cells [41] and human HeLa cells [42] (Table 1). Primer pairs 10 (ENTV-2 U3/gag) and 15 (JSRV U3/gag) have been recently published by our team and shown to be highly specific for JSRV and ENTV-2, with no cross hybridization with ERV sequences [12]. The amplicons obtained with pairs 01 and 13 for each condition (JSRV, ENTV and cell references), with pairs 03 to 08 and 14 for the IDO5 cell reference and with pair 04 for HeLa cells were subjected to Sanger sequencing (TubeSeq Supreme Eurofins) and aligned against XRV and ERV reference proviral sequences. A BLAT search [43] on the human genome (hg38) was performed to identify the nature of the amplicon sequence obtained with pair 04 from human HeLa cells.

**Table 1.** Biological samples: origin and sequences.

| Name | Biological description | Species origin | Genbank accession # | Reference |
| --- | --- | --- | --- | --- |
| FR4403 | French ENTV-1 | *Ovis aries* | PP669280 | (Riocreux-Verney et al., 2024) |
| FR3824 | French ENTV-2 | *Capra hircus* | PP669281 | (Riocreux-Verney et al., 2024) |
| FR2054 | French JSRV Clade I | *Ovis aries* | PP707050 | (Riocreux-Verney et al., 2024) |
| FR989 | French JSRV Clade II | *Ovis aries* | PP707047 | (Riocreux-Verney et al., 2024) |
| IDO5 | Ovine dermal fibroblasts | *Ovis aries* | NA | NA |
| TiGEF | T-immortalized Goat Embryonic Fibroblasts | *Capra hircus* | NA | (Da Silva Teixeira et al., 1997) |
| HeLa | Immortalized human cell line | *Homo Sapiens* | NA | (Scherer et al., 1953) |
| Aa | Mosquito | *Aedes aegypti* | NA | NA |
| Cq/a | Mosquito | *Culex quinque/asciatus* | NA | NA |

A similar approach was taken for the primer pairs published to target ENTV (primer pair #11) and JSRV (primer pair 16 and 17) in arthropods [35,36] by adding DNA extracted from an ENTV-1-induced nasal tumour (FR4403) [12] and mosquitos (*Aedes aegypti and Culex Quique/asciatus*) that have fed on human blood. All the amplicons were subjected to Sanger sequencing (TubeSeq Supreme Eurofins).

### ERV II.5 expression in sheep tissues

Data of mRNA-seq from 60 tissues (Table 2, Ovine FAANG project [44,45]) of the Benz2616 ewe, used to assemble the ARS-UI_Ramb_v2.0 ovine genome (GCA_016772045. 1, [46]), and of lung RNA-seq of JSRVJS21-experimentally infected lambs and the associated uninfected controls [47] were collected from the ENA (European Nucleotide Archive) database. ERV II.5 family - specific k-mers of 21 base pairs were defined in the DR1 and DR3 regions in order to cover the largest number of copies of ERVs annotated in the ARS-UI_Ramb_v2.0 genome (Table 3 and S1 Fig) [6]. These k-mers were used to identify and count the reads corresponding to ERV sequences in the selected RNAseq data using the BBduk analysis tool (Bbtools v38.95, https://quay.io/repository/biocontainers/bbmap) by allowing two mismatches. As a control, k-mers specific to JSRV were defined in DR3 and DR4 and searched in the same RNAseq data (Table 3 and S1 Fig), as previously described [12]. The reads containing the kmers were extracted from the raw data (FastQC file), low-quality sequences and adapters were removed with Trimmomatic [48]. The sequences were then mapped (Bwa-mem, [49]) on the *Ovis aries* family II.5 consensus sequence [6]. The alignment was visualised with IGV (Integrative Genomics Viewer, [50]).

**Table 2.** Tissues used for the ERV expression from RNA-seq data of the sheep Benz2616. Tissues were grouped by physiological systems.

| System | Tissue | ENA accession # | System | Tissues | ENA accession # |
| --- | --- | --- | --- | --- | --- |
| Cardiac | Right ventricle | ERR3728426 | Neurological | Retina | ERR3728428 |
|  | Atrioventricular valve | ERR3728429 |  | Spinal cord | ERR3728434 |
|  | Right atrium | ERR3665709 |  | Pons | ERR3728433 |
|  | Left ventricle | SRR7612070 |  | Thalamus | ERR3728435 |
|  | Vena cava | ERR3728430 |  | Hippocampus | ERR3665708 |
| Immunitary | Spleen | ERR3728431 |  | Pituitary gland | ERR3650379 |
|  | Mesenteric lymph node | ERR3665711 |  | Cerebellum | SRR6651979 |
|  | Prescapular lymph node | ERR3665712 |  | Frontal cortex | SRR7612068 |
|  | Mandibular lymph node | SRR7543059 | Renal | Adrenal medulla | SRR7543054 |
|  | Alveolar macrophage | SRR6651987 |  | Adrenal cortex | SRR6651976 |
| Gastro-intestinal | Ileum | ERR3728427 |  | Renal medulla | SRR7543058 |
|  | Ileum Peyer Patches | ERR3665710 |  | Renal cortex | SRR7612071 |
|  | Cecum | ERR3665706 | Respiratory and buccal | Tongue | SRR7612076 |
|  | Jejunum | SRR7543057 |  | Soft palate | ERR3650378 |
|  | Duodenum | ERR3650374 |  | Epiglottis | SRR6651978 |
|  | Rectum | ERR3728432 |  | Palatine tonsil | SRR7612075 |
|  | Rectum | ERR3665713 |  | Lung | SRR6651986 |
|  | Descending colon | SRR7543055 | Urinary | Urinary bladder | ERR3728424 |
|  | Spiral colon | ERR3665714 |  | Urethra | ERR3665716 |
|  | Esophagus | ERR3665707 |  | Ureter | SRR7543063 |
|  | Abomasum | ERR3665705 | Endocrine | Mammary gland | ERR3728425 |
|  | Abomasum pylorus | ERR3650373 | Genital | Uterus | ERR3665717 |
|  | Reticulum | ERR3650375 |  | Oviduct | SRR7543061 |
|  | Rumen atrium | ERR3650376 |  | Ovary | SRR6651991 |
|  | Rumen ventral | ERR3650377 |  | Vagina | SRR7612077 |
|  | Omasum | SRR7543060 | Muscular | Biceps femoris | SRR6651988 |
|  | Gall bladder | SRR7543056 |  | Semimembranosus | SRR6651990 |
|  | Liver | SRR7612072 |  | Longissimus dorsi | SRR7612073 |
| Skin | Skin hairless | ERR3665715 |  | Supraspinatus | SRR7612074 |
|  | Skin of back | SRR7543062 |  | Diaphragm | SRR7612069 |

**Table 3.**
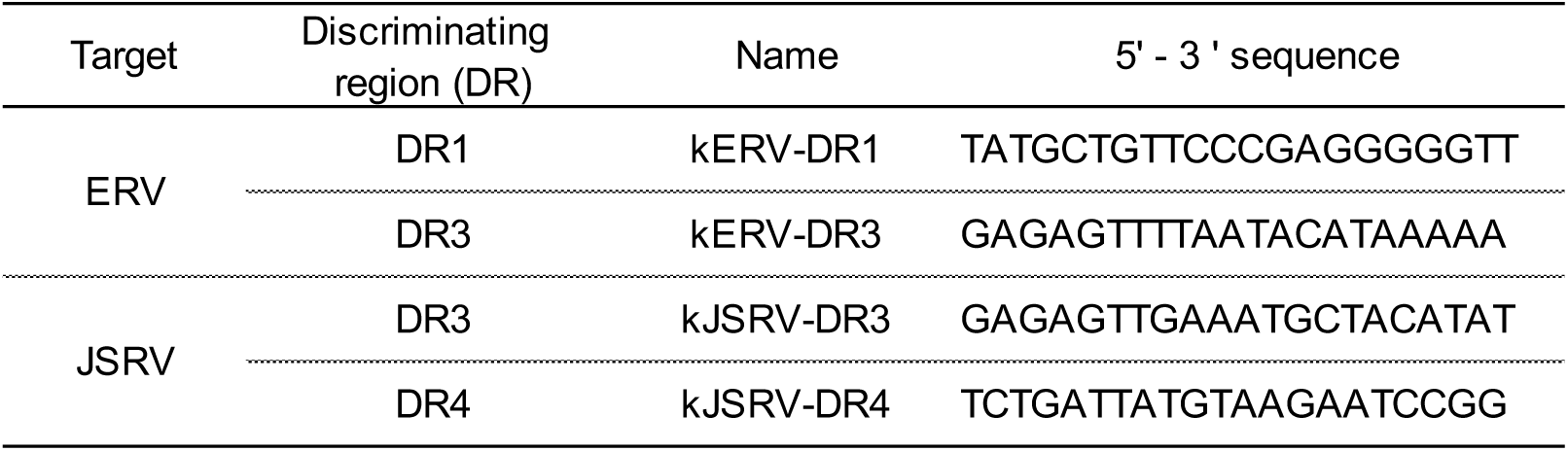
k-mer sequences and localization for the identification 195 of tissue expression of ERVs of family II.5 or the exogenous virus.

### Alignment and phylogenetic analyses

The alignments of *pol*, DR1-DR2, DR3-DR4, complete genomes and sequenced amplicons were carried out using MAFFT Alignment v7.490 [39] and Geneious Prime software (v 2025.0.2). Maximum Likelihood (ML) phylogenetic trees in the *pol*, DR1-DR2, DR3-DR4 regions were generated using IQ-TREE v1.5.3 [51] with 10,000 ultrafast bootstrap and rooting on the BT ERV (Bos taurus ERV) family II.4 consensus sequence [6]. ML-type phylogenetic trees (PhyML 3.3.20180621, Jukes Cantor substitution model) of the *gag* (pair #01, 826nt) and *env* (pair #13, 864 nt) sequences were created using the ENTV1-FR4403 sequence (GenBank #PP669280, [12]) for rooting, with 1000 iterations (bootstraps) and edited through the iTOL software.

Publicly available sequences used in the study are reported in S1 Table. ERV copies from the family II.5, annotated in the *Capra hircus* assembly ARS1.2 (GCA_001704415.2) and the *Ovis aries* assembly ARS-UI_Ramb_v2.0 (GCA_016772045.1), were retrieved from [6]. The partial proviral sequence OR991120 described in [35] and the partial *env* sequence KT266728 [36] were also analyzed.

## Results

### The sections of the genome that differ between ERVs and XRVs are limited

To determine the diversity of XRV sequences, we compiled a dataset comprising 79 complete sequences of JSRV, ENTV-1, ENTV-2 and ERV II.5 consensus sequences. Analysis of the variation, in the percentage of identity of the XRV genomes against the ERV II.5 Oa consensus for the JSRV and ENTV-1, which infect sheep, or the ERV II.5 Ch consensus for the ENTV-2, which infects goats, clearly showed that regions with less than 80% identity between the XRVs and the ERVs are restricted to four discriminating regions, DR1 to DR4. The other regions of the genome do not allow for discrimination between XRVs and ERVs (Fig1). The confidence interval for the ENTV-2 percentage identity is higher than those calculated for JSRV or ENTV-1. In the non-discriminating *pro* and *pol* regions of ENTV-2, the diversity is greater than that observed in the equivalent regions of JSRV and to a lower extent to ENTV-1, notably due to sequences that are genetically close to the goat ERV consensus (S2 Fig, *e.g.* sequences KU258870.1, MK210250.1, ON843769.1, MK164396.1). The DR2 region is less marked in ENTV-1 compared to JSRV and ENTV-2, with a percentage of identity higher than 80%. In addition to the nucleotide polymorphism, the DR1, 3 and 4 regions, displayed specific insertion/deletion (Indels) patterns that can discriminate between exogenous and endogenous sequences (S3 Fig). The insertion in DR2 is only present in ERV II.5 Ch and its length varies between ERV copies (S3B Fig).

Of the 286 PCR primers/probes listed in the 109 published studies, only 28% (79 out 286) were qualified as discriminating between XRVs and ERVs by our *in silico* analysis (S2 Table). Notably, among the 286 primers, 74 (26%) are in the *pro* or *pol* genes, which are regions that are highly similar between XRV and ERV sequences. Similarly, less than one third of the primers in R / U5 region are specific. Therefore, in order to minimize the risk of amplifying ERVs instead of XRV, they must be combined with XRV specific primers.

### Phylogenetic reconstructions reveal the challenge of correctly classifying XRV

Phylogenetic trees, were reconstructed in the *pol* region (Fig 2), which weakly discriminates between XRVs and ERVs, as well as in the DR1-DR2 (Fig 2B) and the DR3-DR4 (Fig 2C) segments from 32 JSRV, 14 ENTV-1, 45 ENTV-2, 27 ERV II.5 Oa and 10 ERV II.5 Ch sequences were extracted from GenBank and supplemented by sequences identified on the assemblies of the *Ovis aries* (ARS-UI_Ramb_v2.0) and *Capra hircus* (ARS1.2) reference genomes [6].

**Fig 1.**
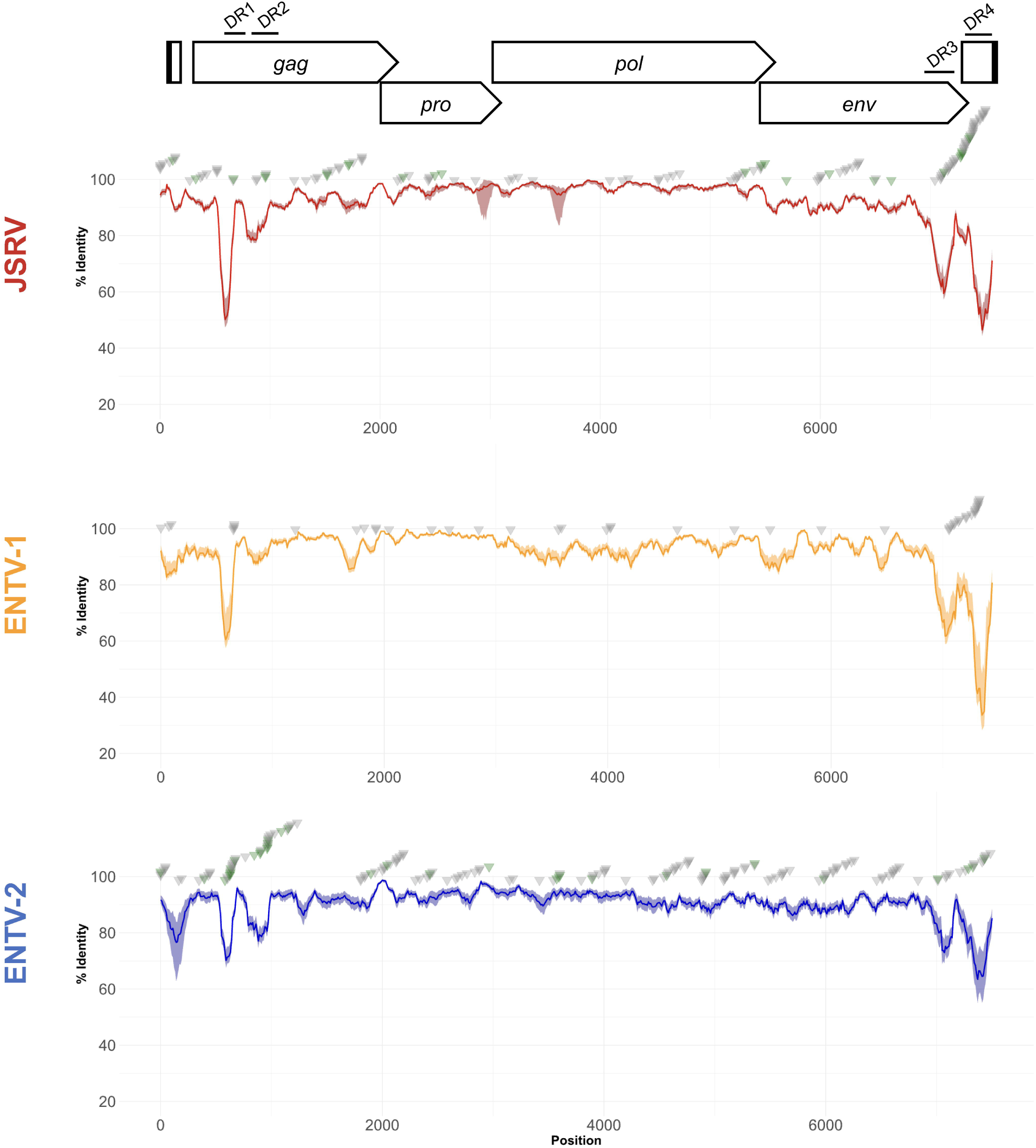
XRV are genetically close to ERVs. The identity score was calculated from a multiple alignment (MAFFT) of XRV sequences against a consensus sequence of ERV II.5 Oa for JSRV and ENTV-1 and ERV II.5 Ch for ENTV-2. The average identity percentage was calculated using a 100-base pair window and plotted in steps of 10. A 95% confidence interval was represented for each virus, using a bootstrap-type repetition approach. The 286 primers (grey triangles ▾), reported in publications (S2 Table) and used for the detection of XRV, were positioned with a tolerance of 4 mismatches. Primers tested in this study are identified by green triangles (▾). DR1-DR4: discriminating regions 1-4.

**Fig 2.**
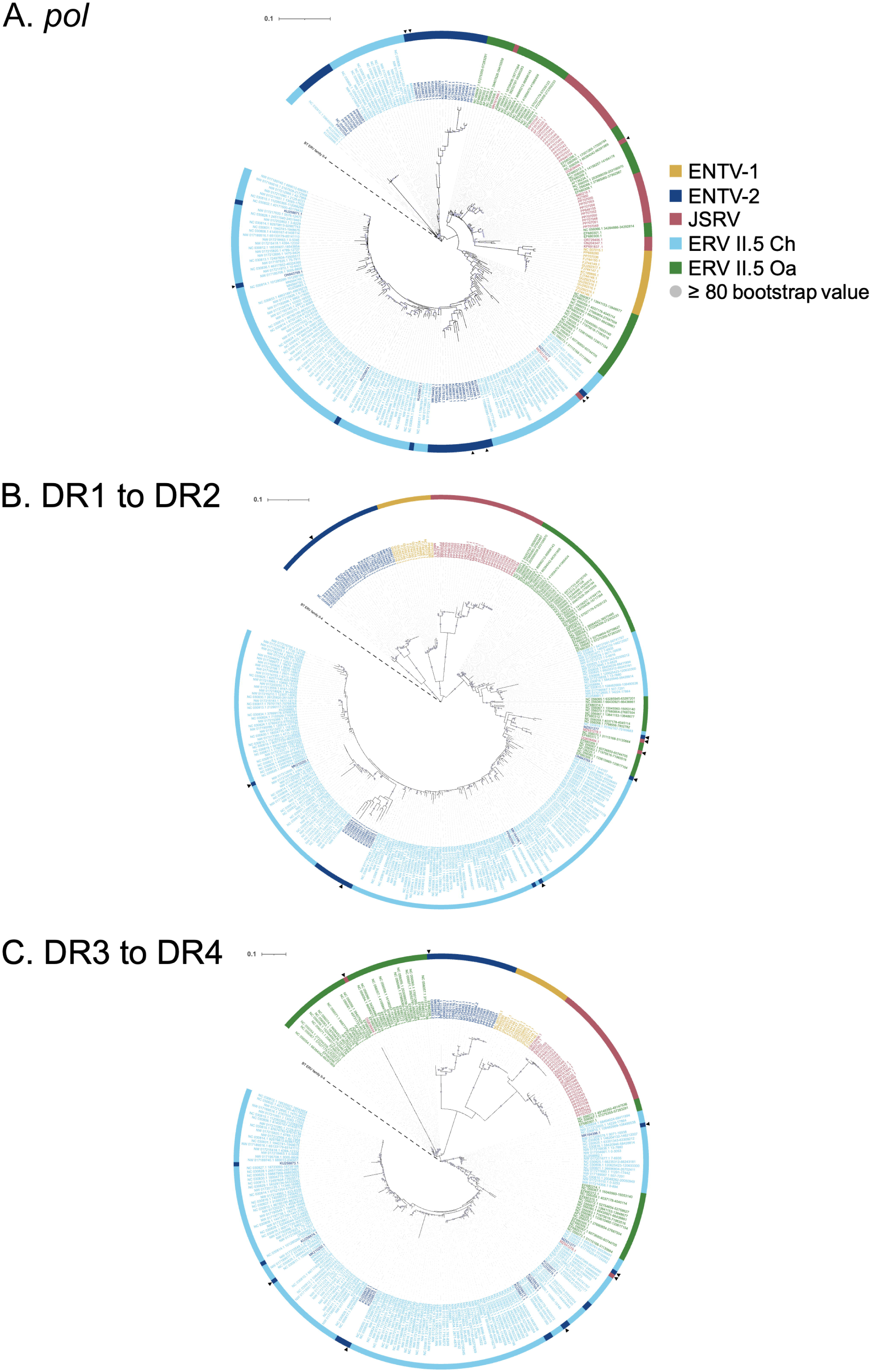
DR1-DR2 et DR3-DR4 are sufficient to classify JSRV, ENTVs and ERV sequences. Full-length sequences XRVs and ERVs available on GenBank by July 2025 and ERV copies annotated in *Capra hircus* and *Ovis aries* were used for phylogenetic reconstructions. The maximum likelihood trees generated using 10,000 ultrafast bootstraps were rooted on the Bt ERV (Bos taurus ERV) family II.4 consensus sequence [6] for **A.** the *pol* gene, **B.** the 360-nt DR1 - DR2 region and **C.** the 449-nt DR3 to DR4 region. The scale bar indicates the number of substitutions per site. Black triangles indicate sequences that have been referred to exogenous JSRV or ENTV despite the absence of the YXXM oncogenic motif in the TM region.

These analyses revealed that the segregation of viral species varies considerably depending on the region considered. The *pol* region did not permit a clear distinction between XRV and ERV II.5 (Fig 2A). Conversely, the DR1-DR2 and DR3-DR4 regions (Fig 2B-C) enabled the different viral forms to be separated into XRVs and ERVs. However, 17 of the XRV sequences were grouped with the ERV sequences, regardless of whether the trees were reconstructed using the DR1-DR2 or DR3-DR4 regions, this calls into question their exogenous origin. Some of the potentially misidentified sequences presented a clear ERV pattern, including specific insertions/deletions (indels) in the DR1-DR4 segment and the lack of the YXXM motif in Env (e.g. MK164396 and MK210250 for ENTV-2 or DQ838494 and MZ931278 for JSRV, Fig 3). The MK210250 sequence displayed a very strong sequence identity (>96%) with the ERV II.5 *Capra hircus* consensus sequence in DR4, whereas the MK164396 sequence did not show a clear pattern, being equally distant from the exogenous and endogenous reference sequences (with an identity percentage ranging from 80 to 83%). Two JSRV sequences, DQ838494 & MZ931278, which cluster with ERVs had DR4/U3 sequences that are very different from the ERV II.5 consensus sequence with 35% and 71% identity respectively, as well as from the JSRV-FR2054 (PP707050.1) reference sequence with 33% and 52% identity, respectively. The BLAT of the DR4 sequence of DQ838494 against the sheep genome (Oar_v4.0/oviAri4) resulted in multiple hits. The first hits (with over 95% identity) corresponded to LINE sequences of the RTE-BovB family, suggested the unmonitored amplification of another transposable element.

**Fig 3.**
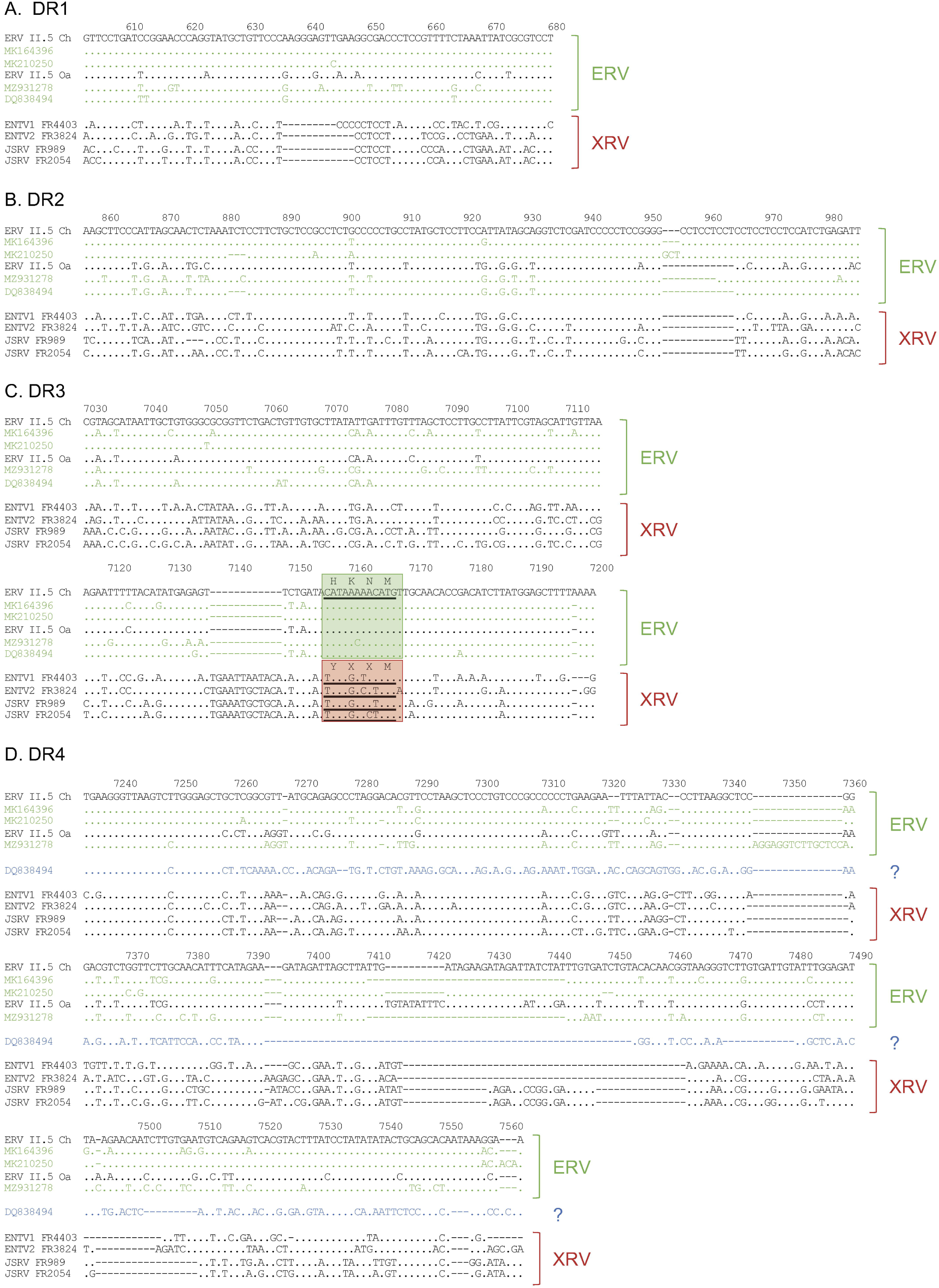
Example of wrongly annotated XRV retroviruses. Viral genome sequences of JSRV (DQ838494 ; MZ931278) or ENTV (MK164396 ; MK210250) strains were aligned (MAFFT) with viral genomic reference sequence of *Capra hircus* and *Ovis aries* II.5 ERV consensus sequences [6], JSRV clade I (FR2054; PP707050.1), clade II (FR989; PP707047.1), ENTV-1 (FR4403, PP669280.1) and ENTV-2 (FR3824, PP669281.1) in **A.** DR1, **B.** DR2, **C.** DR3 and **D.** DR4. Only nucleotides distinct from the ERV II.5 Ch (top sequence) are indicated, identical nucleotides are marked by a dot. The position on the alignment is indicated on top.

Different combinations of mixed XRV/ERV patterns can be observed, such as DR1 and DR3 insertions (“ERV like”) for ENTV-2 ON843769 and MZ931277, and DR3 deletion (“XRV like”) for PP682590. A group of sequences reported as ENTV-2 (KU258870 to KU258880) and integrated into the caprine ERV II.5 phylum does not display the described typical ERV indels profile. Although only KU258870 lacks the YXXM motif, they were still more similar to ERVs in DR1-DR2 and DR3-DR4 regions. No accessible articles describing the conditions under which the sequences were generated were available for any of these sequences except DQ838494 [16] and MK164396 [25], which cluster with ERV II.5 sequences. In the cases of DQ838494 and MK164396, the associated primer sets were predicted to have very low to no exogenous specificity for most primer pairs (S2 Table). This was likely the original cause of the misidentification of these sequences. These genomes, which are most probably ERVs rather than XRV or ERVs/XRV artificial chimeras, have been used as reference sequences in other studies to develop molecular tools [20,21,25].

These analyses highlight the importance of selecting specific regions for detecting XRVs without interfering with ERVs, which are present in multiple copies in ovine and caprine genomes.

### Reliability of XRV detection in an ERV context

Numerous published studies have reported the detection of JSRV and/or ENTV using primers located in regions that are conserved between XRV and ERVs. This makes it impossible to exclude the amplification of ERVs (S2 Table). To illustrate this challenge, fourteen primer sets, defined as specific to JSRV or ENTVs were selected (S3 Table). Specific primers for XRVs were included as specificity control [12].

The primers were defined by the following 3 criteria. The first one was the ability of primers to detect either exogenous ENTV-2 (pairs 1-10) or JSRV (12-15) according to the authors (S2-S3 Tables). The second criterion was the specificity of the primer predicted through XRV and ERV sequences alignment. The third criterion relied on the ability of primers to produce an amplicon of the expected size only from genomic DNA extracted from JSRV- or ENTV-2-induced tumours and not from uninfected ovine, caprine and human cells.

The theoretical specificity of the primers was predicted by aligning them with the XRV and ERV sequences published from the public databases (S1-S2 Tables). Of the ten pairs selected for ENTV-2, primer alignment identified seven pairs (#1, 3, 4, 5, 6, 7, 8) that were non-specific for XRV, due to high homology with ERV sequences. Only three pairs (2, 9, and 10) were specific for ENTV-2. Of the four pairs selected for JSRV, only pair #14 was predicted to be specific for XRV. The primer pairs were then evaluated for their ability to specifically amplify XRV, with amplicon generated only from gDNA extracted from ENTV-2 or JSRV-induced tumours, and conversely to fail to produce signal from gDNA extracted from IDO5, TiGEF or HeLa cells. Amplicons of the expected size from all the gDNAs tested, whether extracted from tumours induced by ENTV-2 as well as by JSRV or from uninfected ovine, caprine were generated for the seven primer pairs # 1, 3, 4, 5, 6, 7, 8 predicted to be non-specific for ENTV-2. More surprisingly, DNA was amplified with the primer pair #4 for human cells. Sequencing confirmed this to be HERV-K (S4 Fig).

Similarly, of the four primer pairs tested for JSRV (12-15), only one pair (#15) was found to be strictly specific, allowing amplification only from lung tumour gDNA; no signal was observed from ENTV-2-induced nasal tumours or from uninfected human, goat and sheep cells, confirming the perfect discrimination between ERVs and exogenous retroviruses (Fig 4). In contrast, the other three primer pairs (#12, 13, 14) produced an amplicon regardless of the gDNA or infection status. This confirms that the published XRV primers are non-specific, enabling, indiscriminate amplification of XRV and ERV sequences.

**Fig 4.**
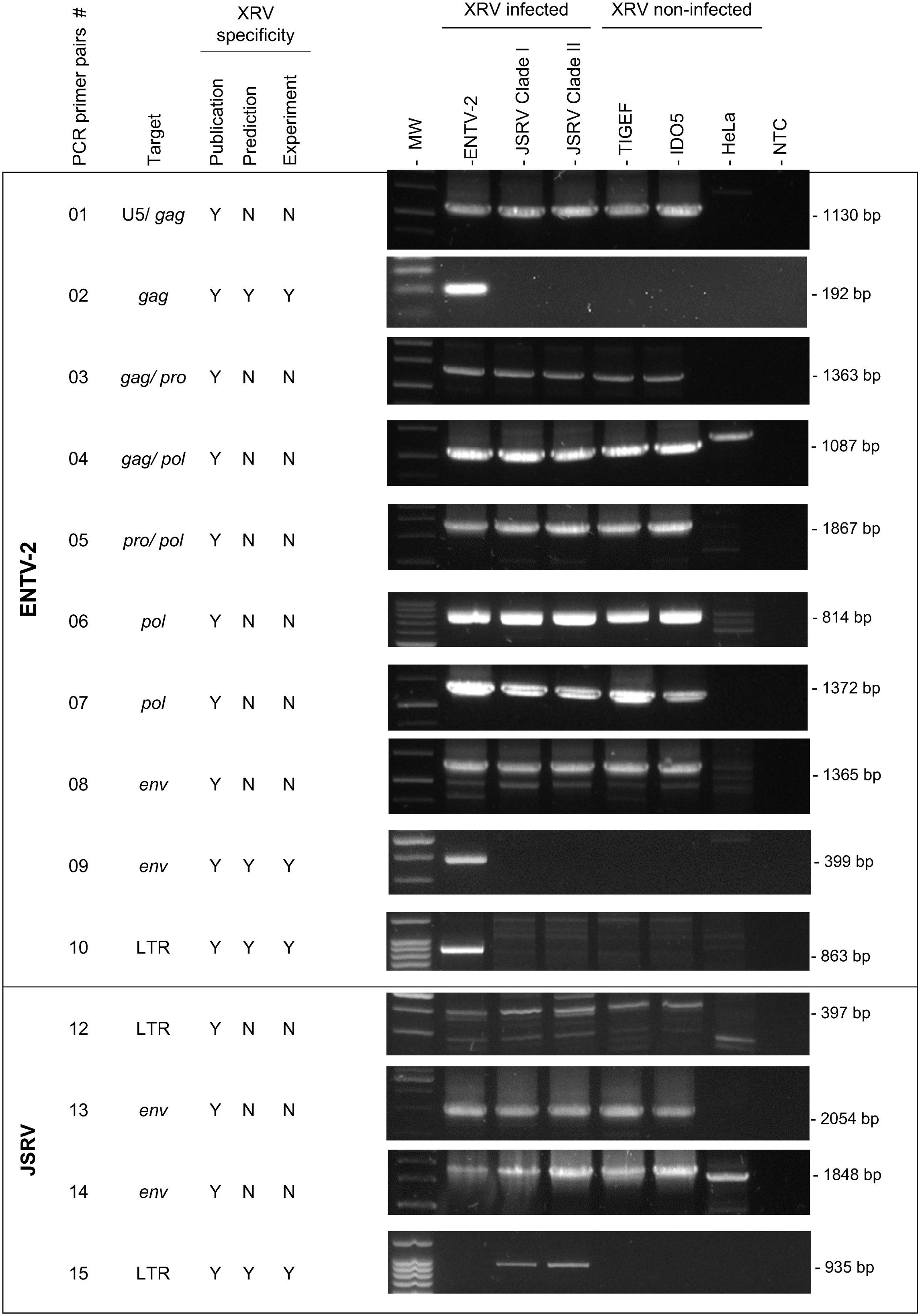
Lack of specificity of selected XRV primers. We analysed the ‘XRV specificity’ of fourteen published primer pairs (01 to 10 and 12 to 15) according to the authors’ statement (‘Publication’), our *in silico* analysis (‘Prediction’) and their profil of detection in PCR (‘Experiment’). The “XRV infected” reference panel consisted of genomic DNAs extracted from a nasal tumour induced by ENTV-2 (FR3824, GenBank accession PP669281.1) and lung cancers induced by clade I (FR2054; GenBank accession PP707050.1) or clade II (FR989; GenBank accession PP707047.1) JSRV. The “XRV non-infected” reference panel consisted of genomic DNAs extracted from goat (TiGEF), sheep (IDO5) and human (HeLa) cell lines. Y: Yes; N: No. NTC: No Template Control.

Amplicons obtained from DNA extracted from IDO5 sheep cells were sequenced. Aligning the obtained sequences with the JSRV and ENTV-2 references sequences (JSRV: FR2054 - PP707050; FR989 - PP707047; ENTV-2: FR3824 - PP669281) and ERV II.5 confirmed their endogenous nature as evidenced by their high homology with ERV II.5 family sequences (S5 Fig).

To unequivocally established their XRV or ERV nature, amplicons obtained with primers containing the discriminating regions DR1 in *gag* (pair #1) and DR3 in *env* (pair #13), were sequenced, and aligned against XRVs and ERV II.5 sequences (Fig 5). The primer pair #1 encompassed DR1 and amplified a region a 1130 bp U5/gag region. The absence of the 12-nt deletion, present only in exogenous JSRV and ENTV sequences, confirms that these primers failed to amplify XRVs (Fig 5A). The absence of the YXXM motif, which is only present in oncogenic XRVs, confirms the amplification of ERV sequences using primers pair #13 (Fig 5B). Phylogenetic reconstruction based on amplicons generated with primer sets #1 (Fig 5C) and #13 (Fig 5D) from DNA of non-infected ovine IDO5 cells or JSRV-induced cancers showed that the sequences were closely related to the ERV II.5 *Ovis aries* consensus sequence (ERV II.5 *Oa*), confirming the amplification of ERVs as part of the sheep genome. Symmetrically, the sequences of amplicons obtained from DNA of non-infected caprine TiGEF cells or ENTV-induced nasal tumours were highly similar to ERV II.5 *Capra hircus* (ERV II.5 *Ch*) consensus sequence (Fig 5C-D).

**Fig 5.**
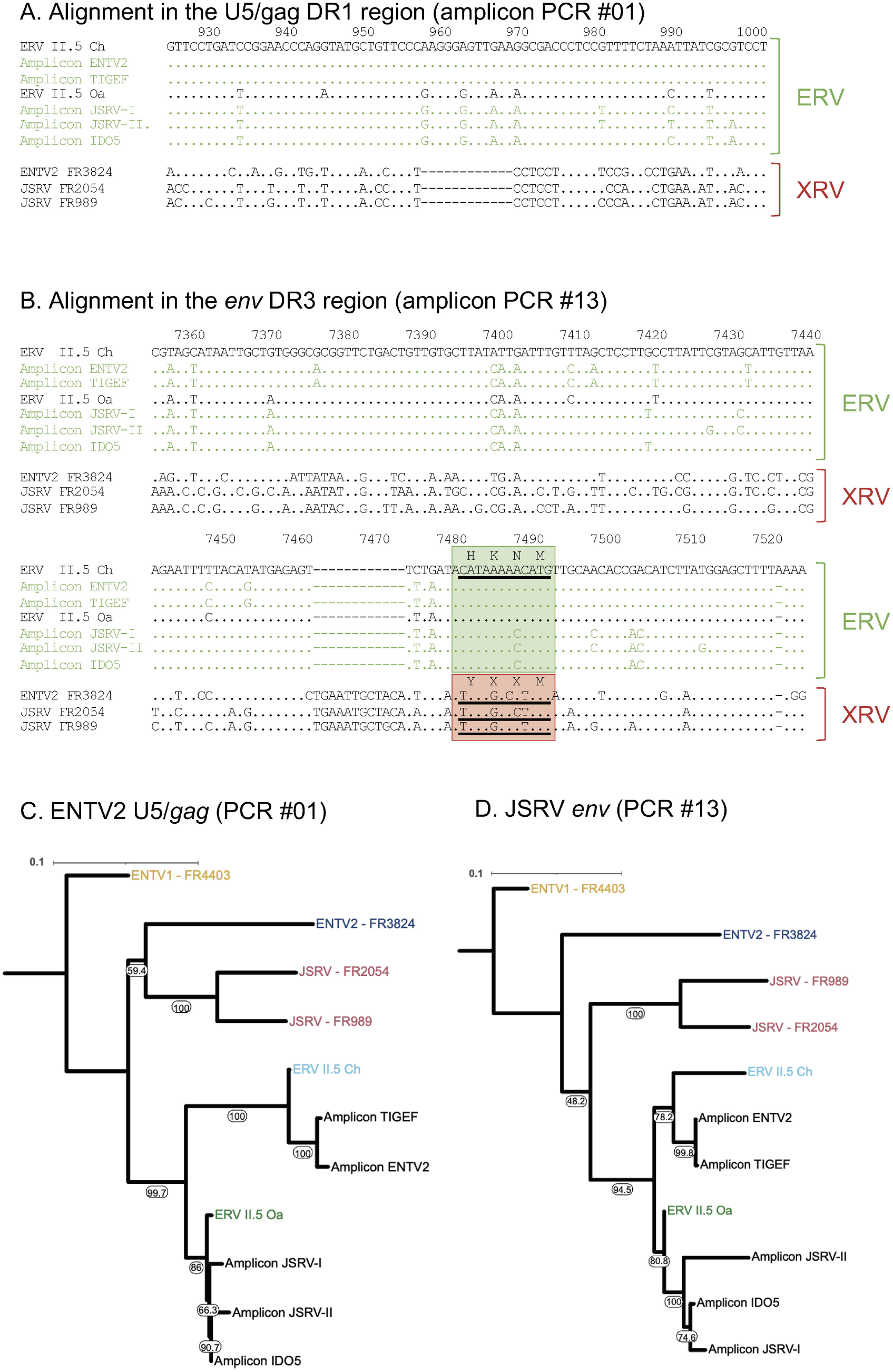
Absence of exogenous DR1 and DR3 discriminatory regions in amplicons obtained with primer pairs #01 (ENTV-2) and #13 (JSRV). Amplicons obtained with primer pairs #1 (ENTV-2) (**A** and **C**) and #13 (JSRV) (**B** and **D**) were sequenced and aligned (MAFFT) with a reference sequence of ENTV-2 (FR3824, GenBank accession PP669281.1), JSRV clade I (FR2054; GenBank accession PP707050.1) & II (FR989; GenBank accession PP707047.1) and the consensus sequences of ERV II.5 Oa and ERV II.5 Ch [4]. Only nucleotides distinct from the ERV II.5 *Ch* consensus sequence (on top) have been indicated, identical nucleotides are marked by a dot. Maximum likelihood (ML) trees were reconstructed using PhyML from **C.** 826-nt of the U5/gag or **D.** 864-nt of the *env* amplicons with ENTV1-FR4403 sequence (GenBank accession PP707036) as an outgroup for rooting. Bootstrap values (1000 replicates) are expressed as a percentage and indicated at the node level. The scale bar indicates the number of substitutions per site.

### Multiple tissues express endogenous sheep and goat retroviruses

Several studies have been based on amplifying JSRV and ENTV from RNAs under the assumption that only exogenous retroviruses are expressed. As RNA is an appropriate target for retroviruses detection, the absence of ERV expression must be confirmed to rule out dual expression of XRVs and ERVs at the same sites. We analysed the expression of ERV II.5 using transcriptomic data from 60 tissues of the Benz2616 ewe, the source of ARS-UI_Ramb_v2.0 reference ovine genome. Short sequences or k-mers were defined in DR1 in *gag* and DR3 in *env* (Fig 6, Table 3 and S1 Fig) from annotated ERV II.5 copies of the ARS-UI_Ramb_v2.0 genome [6]. JSRV k-mers located in DR3 and DR4 were used to detect JSRV expression from RNA-Seq data from JSRV-experimentally lambs and confirm the JSRV-negative status of the Benz2616 ewe (Fig 6A, Table 3 and S1 Fig) [12,47]. For each of the RNA-Seq datasets, sequences with specific ERV or JSRV k-mers were counted using the BBduk tool. Conversely, ERV k-mers identified small quantities of ERV transcripts in most of the tested tissues, with notably higher expression in the lung and kidney (particularly the internal medullary part) and oviduct (Fig 6B). Sequences identified by the ERV k-mers were mapped to the consensus sequence of ERV II.5 Oa, confirming their endogenous origin proven the specificity of this approach to identify transcripts (Fig 6C). This demonstration in sheep shows that amplifying RNAs does not guarantee the sole detection of JSRV or ENTV. If the primers are not highly XRV-specific, ERVs will be amplified. Additionally, the absence of cellular DNA, even as traces, must be carefully controlled to prevent the amplification of ERV copies, present in multiple copies.

**Fig 6.**
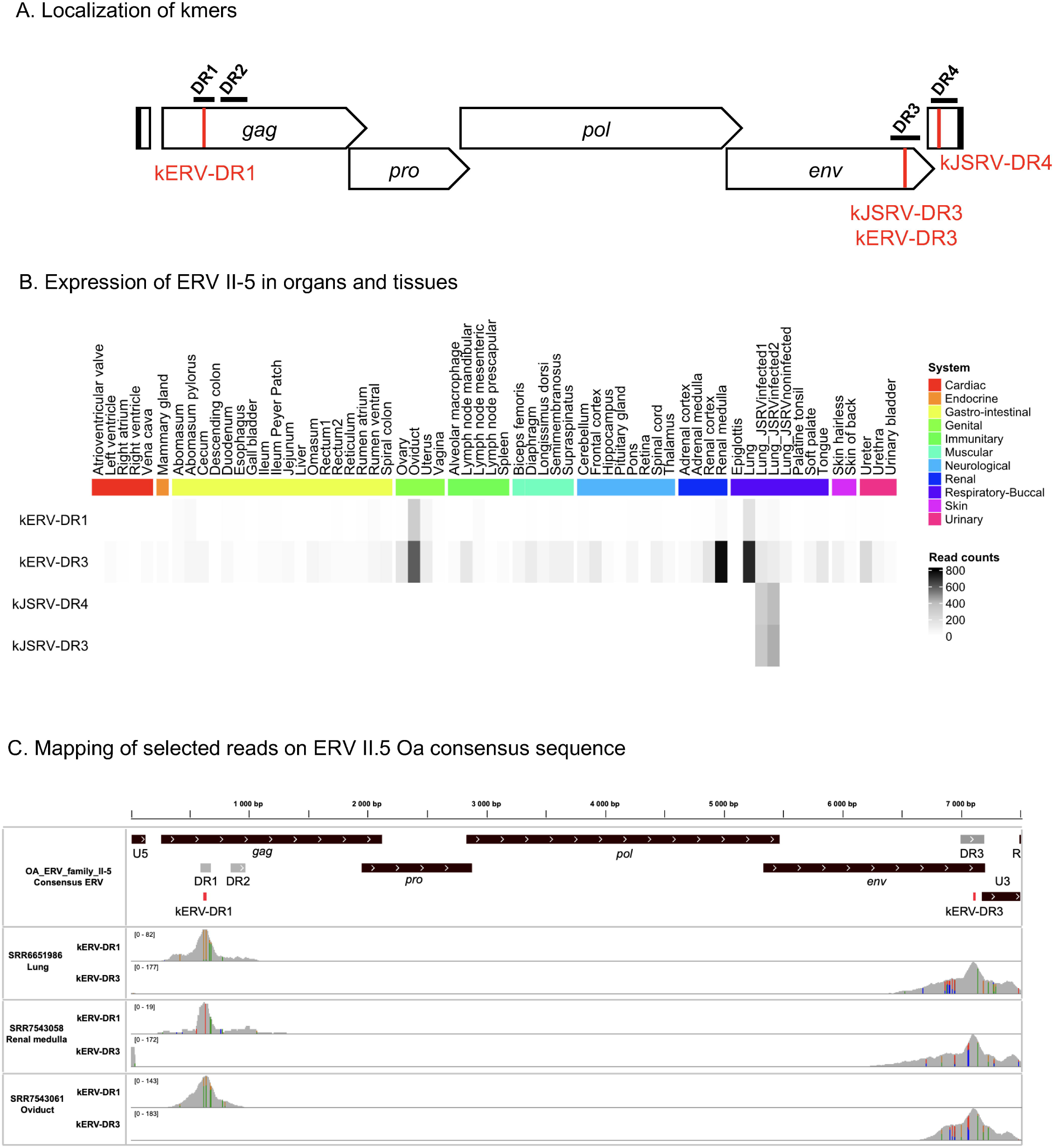
Expression of ERV family II.5 in tissues. **A**. K-mers localization on the JSRV provirus. kERV-DR1 and kERV-D3 are specific to ERV II.5 Oa and located in the DR1 (*gag*) and DR3 (*env*) discriminating regions, respectively. kJSRV-DR3 and JSRV-DR4 are specific to JSRV and located in DR3 (*env*) and DR4 (U3). **B**. The number of reads containing one of the four selected k-mers was calculated for each tissue or organ. The samples were grouped by physiological system and visualised using a colour code. JSRV-infected and uninfected control sample [47] were added and labelled as “Lung_JSRV_infected” for lung RNA-Seq data of JSRV-JS21-infected lamb or “Lung_JSRV_uninfected” for uninfected control lambs. **C.** Reads containing kERV-DR1 and kERV-DR3 k-mers were mapped on the ERV II.5 Oa consensus sequence and visualised on IGV.

### The presence of JSRV and ENTV in arthropods cannot be not confirmed

Six recent studies identified small ruminant XRV sequences [32–37] in insect vectors using global approaches, such as metagenomics, transcriptomics and whole RNA-Seq enriched for viral RNA. We used these datasets to challenge both our strategy for specifically detecting XRVs in a context of coexistence of both XRV and ERVs in the same individuals and the hypothesis of a potential vectorisation. Those approaches rely on the BLASTn of the *de novo* assembled genomes to identify reference sequences. Two studies made the reference matches available [33,36]. While analysing the different datasets, we observed that identification of sequences deposited in GenBank can be misleading. For example, MN564750.1 used as a reference, is associated in GenBank with the organism “Enzootic nasal tumor virus 2” (Taxonomy ID: 2913605), ID which also includes “Endogenous enzootic nasal tumor virus”. MN564750.1 is identified as “Enzootic nasal tumor virus 2 isolate enENTV-FJ2 genomic sequence”. A scientist familiar with these viruses would recognise that the name of the isolate ‘enENTV-FJ12’ corresponds to an endogenous (en) virus. However, this is confusing, particularly in the absence of an associated article or any further information about the origin in the deposit entry. We clearly identified ERV sequences wrongly tagged as XRV in public databases such as MK210250 (Figs 2 and 3) or KT266728 (S6 Fig) used as a reference respectively for the detection of ENTV-2 [33] and JSRV [36] which both lack the sequence coding for the YXXM motif. Overall, this detailed initial analysis highlights the importance of carefully selecting the reference sequence to exclude ERV sequences.

In their study [36], the authors identify both sequences of JSRV and ENTV-2 within mosquitoes but only validated the presence of JSRV. While the amplification and sequencing led to the identification of the sequence as “enJSRV”, the validation was initially biased as primers (S2-S3 Tables) not specific for exogenous viruses were designed based on the sequence initially matching with their reference. The non specificity of primers #16 and #17 was confirmed using the same experimental design with reference samples of DNA extracted from XRV-induced tumours, negative sheep and goat controls, including DNA extracted from different mosquito species. The amplification pattern was non-specific, with amplification in both uninfected and infected samples (Fig 7A). The #16 and #17 primer sets amplified regions confirmed to be of ERV origin by sequencing. The DR3 amplicon obtained with PCR #16 (Fig 7B) does not contain the insertion specific of XRVs nor the sequence coding for the YXXM motif; the DR4 amplicon obtained with PCR #17 (Fig 7 A-C) has both the size and sequence compatible with the ERV II.5 consensus. Neither the sequence nor the accession number were made available for the “nearly complete sequence” of ENTV-2 identified in mosquitoes. However the authors reported that the sequence was closely related to the GDQY2017 Chinese strain (accession number MK164396), a sequence that we identified as ERVs, questioning the XRV nature of the ENTV-2 sequence identified in this study (Figs 2 and 3) [36].

**Fig 7.**
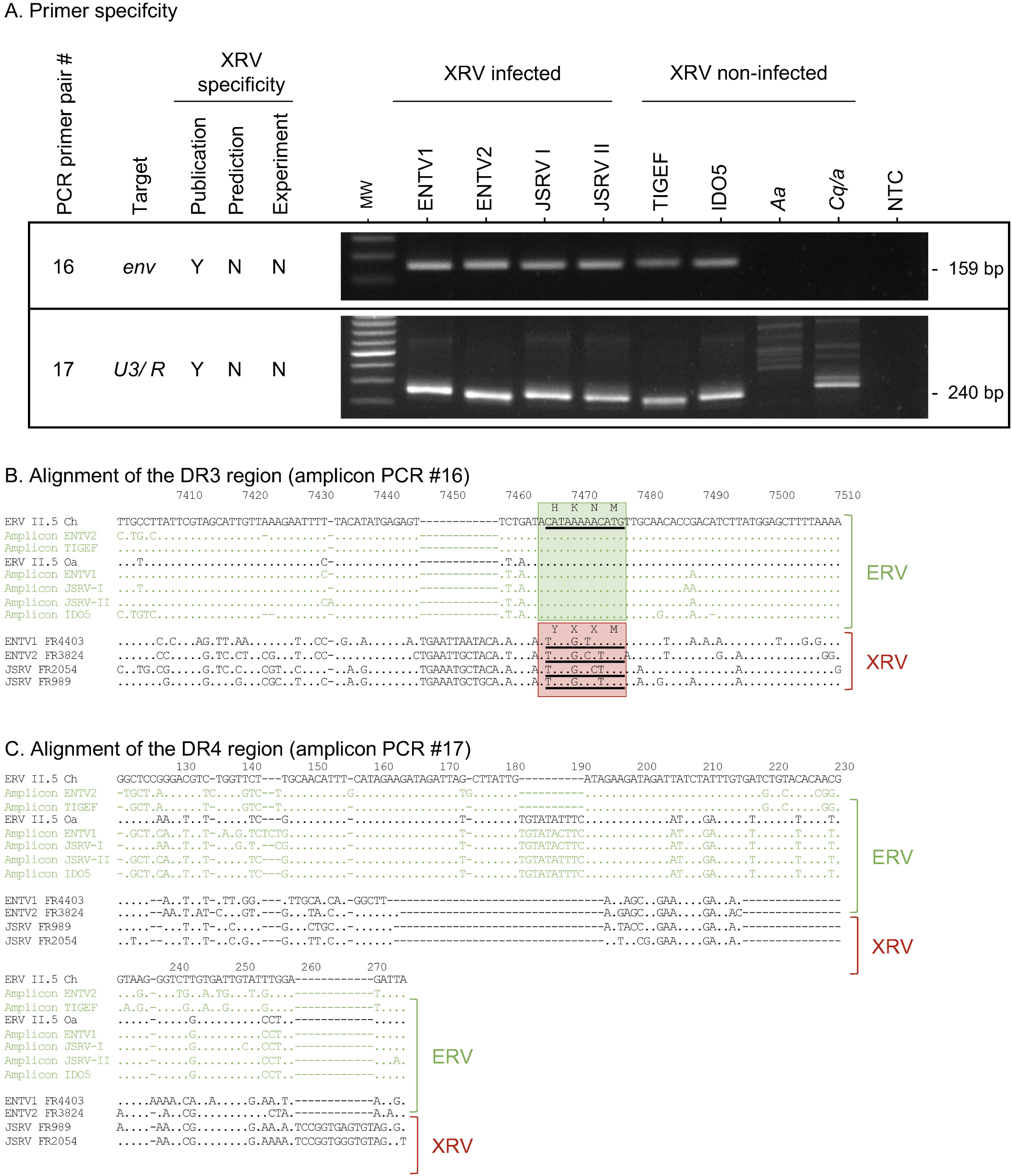
Molecular detection of JSRV and ENTVs in mosquitoes. **A.** Evaluation of the specificity of primer pairs # 16 and #17 on genomic DNA extracted from JSRV or ENTV tumours, from uninfected *sheep (IDO5) and goat (TIGEF)* cell lines, and from *Aedes aegypti (Ae)* and *Culex Quique/asciatus (Cq/a*). The amplicons obtained with primer pairs #16 (**B**) and #17 (**C**) were sequenced and aligned against ERV II.5, ENTV and JSRV reference sequences. Only nucleotides distinct from the ERV II.5 *Ch* consensus sequence (on top) have been indicated, identical nucleotides are marked by a dot.

In a recent study [35], ENTV was amplified and sequenced from ticks collected in Pakistan. A nearly full-length proviral genome (OR991120) was amplified (Fig 8A). The sequence displayed 98% and 92 % overall nucleotide identity respectively with the caprine and sheep ERV II.5 consensus, but less than 89% with JSRVs and ENTVs. The DR3 and DR4 regions were unavailable. However, the DR1 and DR2 regions displayed indels characteristics of ERVs (Fig 8C-D), as well as an ERV-specific partial insertion in DR2 region. These findings suggest that the “provirus” was amplified from a goat ERVs. The single set of primer targeting the *gag/ pol* was not predicted as specific of XRVs, allowing an amplicon from both XRV infected and uninfected sources (Fig 8B). The DR1 and DR2 sequencing of the amplicon confirmed the ERV nature of the sequence (Fig 8C-D).

**Fig 8.**
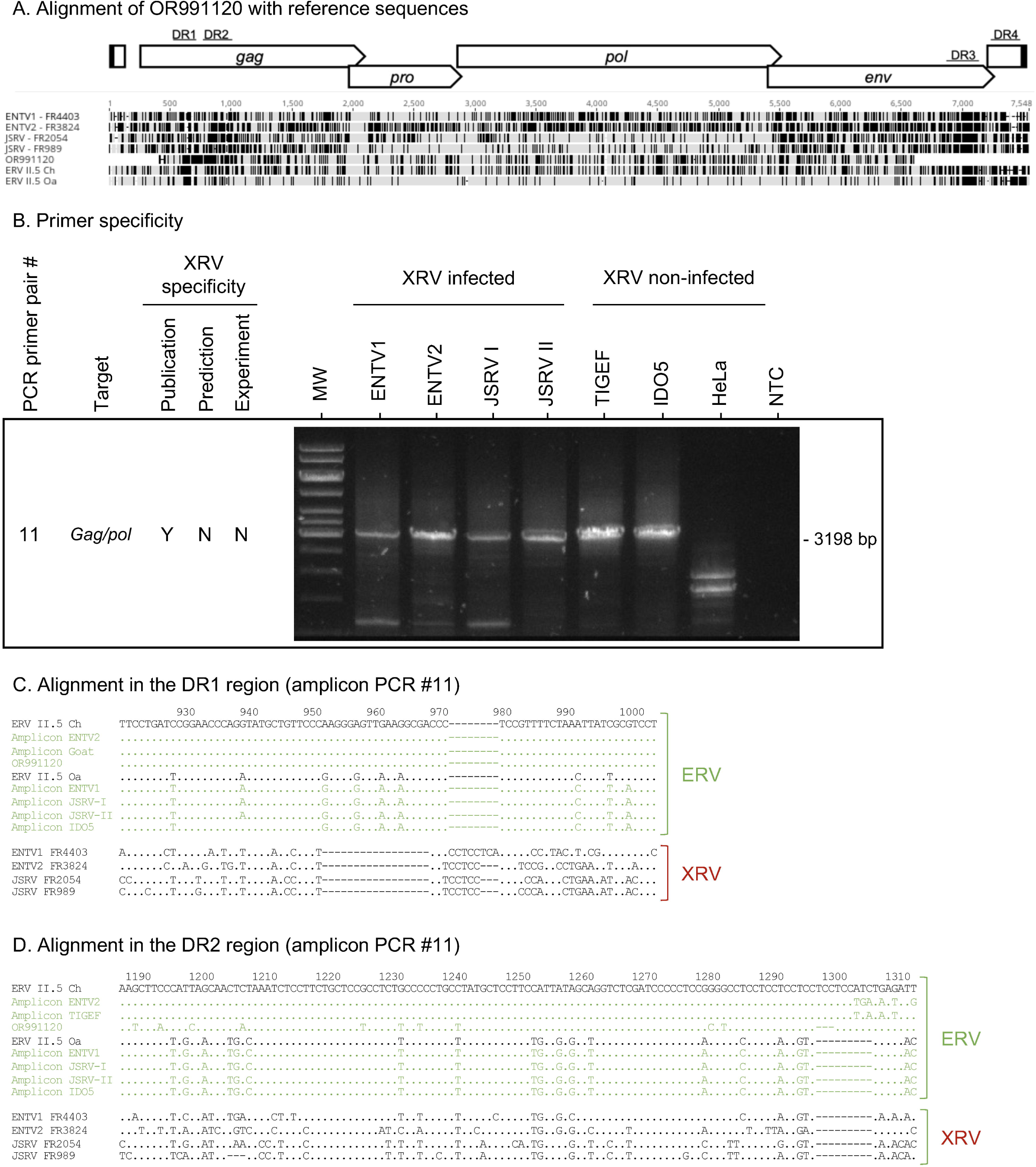
Molecular detection of ENTV in ticks. **A**. The partial OR991120 proviral sequence, missing the LTRs and the DR3 region, reported in ticks [35] was aligned with II.5 ERV Oa and Ch consensus sequences [6], JSRV clade I (FR2054; GenBank accession PP707050.1), clade II (FR989; GenBank accession PP707047.1), ENTV-1(FR4403, GenBank accession PP669280.1) and ENTV-2 (FR3824, GenBank accession PP669281.1). The black and grey bars correspond respectively to the differential or identical nucleotides. **B**. Evaluation of the primer specificity. The amplicon #11 was sequenced and aligned against proviral sequences of JSRV, ENTV and ERV II.5. **C.** Alignment of the DR1 region. **D.** Alignment of the DR2 region.

Three studies identified ENTV-2 in ticks isolated from cattle [32], JSRV from sheep [37], ENTV-2 and JSRV from ticks isolated from yaks and ENTV-1 from Tibetan sheep [34]. Unfortunately, neither data on the overall or DR regions coverage were available, nor molecular validation of the presence of JSRV and ENTVs with XRV specific tools were reported, only information regarding the abundance of the reads were available for [32] and [34]. Therefore, the conclusions that can be drawn about the exogenous nature of these sequences are limited, without further investigations. *A contrario* a thorough checking based on the current recommendation can dispel doubts about the nature of the strains detected by these high-throughput approaches. We therefore collaborated with the authors to rework the preprint [33], which initially described ENTV-2 within ticks. One of the tick metagenome samples displayed coverage of the ENTV-2 provirus of over 80%, thus enabling to distinguish between ERV/XRV and clearly identifying the DR1-3 regions associated with ERV. DR4 coverage was lower but also suggested the detection of ERV. Phylogenetic analysis confirmed these conclusions, as the consensus sequence clustered with the ERV sequences (S7 Fig).

## Discussion

Accurate detection of β-retroviruses responsible for respiratory cancers in sheep and goats is a critical prerequisite for controlling these diseases in livestock. This task is complicated by the coexistence of exogenous retroviruses (JSRV and ENTV) with numerous copies of endogenous retroviruses belonging to the ERV II.5 family in the genome of sheep and goats. Molecular diagnostic approaches must therefore allow unequivocal detection of XRV and therefore target regions that clearly discriminate between ERVs and XRVs. Insufficient consideration of this constraint has directly impacted on the specificity of molecular assays, the interpretation of phylogenetic relationships, and the integrity of sequence annotations.

Early studies, based on a limited number of available sequences, suggested that regions 1 to 4 could discriminate between ERVs and XRVs due to their low sequence homology [5,19]. Building on these observations, we conducted an extensive analysis integrating available XRV and ERV sequences with *in silico* annotation of ERV II.5 loci in sheep and goat assemblies, that allowed us to define four discriminating segments (DR1-DR4) that robustly discriminate exogenous from endogenous retroviruses. DR1 and DR2 localize in the 5‘gag region, DR3 in the 3’ terminus of *env*, and DR4 to U3. Thus, they are prime targets for virus detection, reconstruction of phylogenetic relationships and sequencing of near-full length of exogenous β-retroviruses. As we established in this study from *in silico* analysis of primers reported in the literature, targeting other regions with limited variation between ERVs and XRV forms poses a risk of ERV detection.

As frequently reported in the literature, primers targeting conserved regions shared between ERVs and XRVs are inherently non-specific and prone to amplifying endogenous sequences. As we established in this study, most primers used to detect JSRV or ENTV are unable to distinguish between exogenous and endogenous sequences present in animal genomes and if used must be paired with XRV specific primers. In a similar way, the growing use of high-throughput sequencing without adequate discrimination criteria has amplified this problem. In recent years, this methodological flaw has led to widespread mis-annotation of ERVs as infectious JSRV or ENTV, distorting evolutionary analyses and misleading the development of molecular tools.

Misinterpretation extends beyond host samples. Recent reports suggesting JSRV or ENTV presence in ticks or mosquitoes may most likely reflect the detection of goat or sheep DNA acquired from blood meal rather than true viral infection. As we have shown there is no evidence that arthropods could serve as reservoirs or vectors for JSRV and ENTV. Xenosurveillance may help to identify infection [36] but the clear frontier with ERVs has to be drawn.

The distinction between XRV and ERV is often blurred in sequence databases, such as GenBank, due to the presence of redundant and inconsistent taxonomic identifiers that merge XRV and ERV. While a single taxonomic ID (69576) is defined for ENTV-1, JSRV is classified within the taxonomic ID 47898 (recognised by the ICTV), which also includes the “Endogenous Jaagsiekte retrovirus**”** (also classified with the *Ovis aries*, taxonomic ID: 9940). A subclass “Exogenous Jaagsiekte sheep retrovirus” (Taxonomic ID: 47898) is also reported, but this only contains 6 sequences. For ENTV-2 the problem is even more complex. Recently, four taxonomic IDs coexisted for the same virological entity, but taxonomic ID 2762664 (Endogenous enzootic nasal tumor) and taxonomic ID 239365 (Enzootic nasal tumour virus of goats) were recently merged with taxonomic ID 2913605 (Enzootic nasal tumor virus 2). The fusion of taxonomic ID 2762664 with 2913605 artificially grouped XRV and ERV sequences, whereas the latter could be defined as part of the goat genome with identifier 9925 (*Capra hircus*). Finally, the only singularity of taxonomic ID 2584748 is that it is defined by its common name in US English, with “tumour” instead of “tumor”.

The classification of ERVs based on whether they are considered part of the virus or the host is still a challenge, not limited to retroviruses in small ruminants. As an example, porcine endogenous retroviruses are associated in GenBank either to their host genome, *Sus scrofa* (Taxonomic ID 61673) or to the viral entity PERV (Porcine endogenous retrovirus) (Taxonomic ID: 61673) subdivided in 11 subclasses with their own taxonomic IDs. Brait and colleagues [38] suggest a rational solution to bypass this challenge by adding a second taxonomic identifier. In this context, referring to ERV II.5 with the taxonomy ID of the host in which they have been identified and a second taxomic ID related to the closest XRV offers a pragmatic solution.

Finally, genome assembly approaches using short amplicon sequencing are ill-suited to contexts where ERVs and XRVs coexist, as they cannot exclude chimeric reconstructions. Long read sequencing technologies are therefore preferable an less prone to generate *in silico* artefacts [12]. Regardless the approach used, the genetic nature of the DR1 to DR4 discriminating regions is a minimal requirement for classifying sequences as exogenous. Hallmarks features such as deletions in DR1/ DR2 regions as compared to ERV II.5, and the presence of the exogenous oncogenic YXXM motif in DR3 (strictly absent in ERV) provide unambiguous evidence of exogenous origin.

Although most of the complete sequences of JSRV and ENTV present in GenBank possess these motifs indicating a probable exogenous origin, doubts remain about the non-discriminatory regions such as the end of *gag*, *pro*, *pol* and the beginning of *env*. For example, a recent study suggests possible recombination between different strains of ENTV-2, precisely in the *pro* and *pol* regions, in which the absence of the XRV/ ERV discriminatory region complicates amplification and the specific identification of sequences of exogenous origin [21]. In this case, the reconstruction of chimeric XRV/ ERV sequences cannot be ruled out.

The DR1-DR4 regions exhibit sufficient polymorphism to resolve viral clades and geographic lineages. Updated diagnostic tools must therefore balance specificity for exogenous viruses with coverage of known diversity.

In conclusion, we urge the stringent validation of the molecular tools used to detect JSRV, ENTV-1, and ENTV-2. Primer pairs should include at least one primer located in discriminating regions DR1–DR4, and the absence of amplification from DNA of uninfected ovine and caprine cells specificity must be confirmed to establish specificity. RNA-based detection alone is insufficient to exclude ERV contamination, particularly in complex clinical samples and if we consider that ERV expression is present in various tissues. Finally, assembled viral genomes lacking exogenous hallmarks in DR1–DR4 using separated amplicons should be considered as potential chimeric genomes, and not the reflect of the circulating strains. Only a rigorous, discrimination-based framework will restore confidence in published sequences, improve database reliability, and prevent further propagation of ERV/XRV confusion.

## Acknowledgements

We thank Loïc Schwaller and Emmanuelle Lerat for the scientific advice; Vincent Raquin and Barbara Viginier for providing us the mosquitoes. This work was performed using the computing facilities of the CC LBBE/PRABI.

## Funding

This research was funded by ANR, grant ANR-22-CE35-0002-01 and by INRAE “AAP-Action incitative sur priorités SSD 2021-2025”. BRV was the recipient of a VetAgro Sup PhD grant. MV PhD fellowship was funded by ANR, grant ANR-22-CE35-0002-01.

## Data availability

This article used publicly available data. All the accession numbers from public databases and sequences extracted from the articles were made available in the supplementary tables 1 and 2.

## Conflict of Interest

The authors declare that the research was conducted in the absence of any commercial or financial relationships that could be construed as a potential conflict of interest.

## Author Contributions

BRV and CD realized the experiments. BRV, MV, JT, CL performed the analyses. BRV, VN, MV, CL, and JT designed the analysis. BRV, JT and CL wrote the manuscript. CL and JT conceived the project and secured the fundings. CL, JT, and VN supervised the work.

All participants contributed to the design and implementation of the research, participated to the interpretation of the data, and revised the manuscript.

## Legends

**S1 Table. Accession number list of all full-length and near full sequences of JSRV, ENTV-1, ENTV-2 and ERVs used in the study.**

**S2 Table. Description of the 286 JSRV and ENTV published primers used in the study**.

The ‘Publication specificity’ column relies on the authors’ statement of whether the primer pair was specific (“S”) or not (“NS”) for JSRV or ENTVs in the corresponding article(s). The ‘Predicted XRV’ column is based on *in silico* control of the specificity for XRV (Yes) or non-XRV (No). “ND” indicates that the XRV or ERV specificity could not be determined.

**S3 Table. Sets of primers tested for XRV detection and PCR conditions.**

**S1 Fig. Characterisation of the k-mers used to detect ERV II.5 and JSRV reads.** The k-mers were identified from multiple alignments (MAFFT) of JSRV strains present in GenBank, whose exogenous nature was validated, and the copies of ERV II.5 Oa annotated in the ARS-UI_Ramb_v2.0 genome which contains the DR1 region (A), the DR3 region (B) or the DR4 region (C). Mismatches against the chosen k-mers are highlighted.

**S2 Fig. Individual identity scores per XRV sequence**. The individual identity scores for all the XRV sequences were calculated using a multiple alignment (MAFFT) of the XRV sequences against an ERV II.5 Oa consensus for JSRV (A) and ENTV-1 (B), and against an ERV II.5 Ch consensus for ENTV-2 (C). The average identity percentage was calculated using a 100-base pair window and plotted in steps of 10.

**S3 Fig. DR1-DR4 discriminating regions between endogenous retroviruses and JSRV & ENTVs.** Viral genomic sequence of ERV II.5 Oa and Ch consensus sequences [6] and JSRV clade I (FR2054; PP707050.1), clade II (FR989; PP707047.1), ENTV-1 (FR4403, PP669280.1) and ENTV-2 (FR3824, PP669281.1) were aligned using MAFFT. **A.** DR1 and **B**. DR2 in the *gag* gene. For DR2 the percentage of each nucleotide (A,T,G,C) per position for all the II.5 copies in the Goat (ARS1.2) and Sheep (ARS-UI_Ramb_v2.0) references was plotted in the deletion area. **C.** DR3 in *env* and **D.** DR4 in U3. Only distinct nucleotides with the first sequence of the alignment (ERV II.5 Oa) have been indicated, identical nucleotides are marked by a dot. Numbering corresponds to the position on the alignment.

**S4 Fig. HERV-k sequence obtained with primer pair #4, supposedly specific of ENTV-2 from genomic DNA of human HeLa cells.** Alignment (MAFFT) of the sequence of PCR fragment #4 obtained by the Sanger method with the closest β-HERV (HERVK-13) identified by BLAT.

**S5 Fig. Sequencing amplicons obtained from ovine IDO5 cells.** MAFFT alignment of the amplicon sequence with ERV II.5 Oa consensus sequence. **A.** Primers set #3. **B.** Primers set #4. **C.** Primers set #5. **D.** Primers set #6. **E.** Primers set #7. **F.** Primers set #8. **G.** Primers set #14.

**S6 Fig. KT266728.1 Env sequence does not contain the oncogenic « YXXM » motif.** The JSRV viral sequences KT266728.1, reported as the best match in [36], was aligned (MAFFT) with the ERV II.5 Oa consensus sequence [6] as well as with JSRV clade I (FR2054; PP707050.1) and clade II (FR989; PP707047.1) sequences. The nucleotides encoding YXXM are not present in KT266728.1; this strain has a HKXM predicted sequence that is common to ERV II.5.

**S7 Fig. Verification of metagenomic sequences identified as exogenous.** The consensus sequence (TickP118) obtained from one tick metagenome from [33] and the viral genomic sequence of ERV II.5 Oa and Ch consensus sequences [6] and JSRV clade I (FR2054; PP707050.1), clade II (FR989; PP707047.1), ENTV-1 (FR4403, PP669280.1) and ENTV-2 (FR3824, PP669281.1) were aligned using MAFFT in A. DR1, B. DR2, C. DR3 and D. DR4. Only distinct nucleotides with the first sequence of the alignment (ERV II.5 Ch) have been indicated, identical nucleotides are marked by a dot. Numbering corresponds to the position on the alignment. Full-length sequences of ENTV-2 and enENTV available on GenBank and the consensus sequence obtained from one tick metagenome from [33] were used for phylogenetic reconstructions. Maximum likelihood (ML) trees were reconstructed using PhyML **F.** the 449-nt DR3 to DR4 region. Bootstrap values (1000 replicates) are expressed as a percentage and indicated at the node level. To help the visualisation the tree was rooted on the ERV branch. The scale bar indicates the number of substitutions per site.

## Notes

### Competing Interest Statement

The authors have declared no competing interest.

### Summary of Updates

Addition of the supplementary figures and tables

