## Supplemental Table 1 for "Unraveling viral identity: Avoiding the trap of endogenous sequences for viral surveillance of small ruminant oncogenic retroviruses"

| | II.5 Ch | II.5 Oa | JSRV | ENTV-1 | ENTV-2 |
|---|---|---|---|---|---|
| Taxonomic IDs #: | 2762664 & 2913605 (with entries including “endogenous” or “enENTV” in their “definition”) | 9940 | 11746 and 47898 | 69576 | 2913605, 2913605, and 2584748 |
| | KU258881.1 | AF136224 | A27950.1* | FJ744146.1* | HM104174.1 |
| | KU258882.1 | AF136225.1 | AF105220.1* | FJ744147.1* | KU179192.1 |
| | KU258883.1 | AF153615.1 | AF357971.1* | FJ744148.1* | KU258870.1* |
| | KU258884.1 | EF680296.1 | CQ964469.1* | FJ744149.1* | KU258871.1* |
| | KU258885.1 | EF680297.1 | DQ838494.1 | FJ744150.1* | KU258872.1* |
| | KU258886.1 | EF680298.1 | KP691837.1* | GU292314.1* | KU258873.1* |
| | MN564749.1 | EF680299.1 | M80216.1* | GU292315.1* | KU258874.1* |
| | MN564750.1 | EF680300.1 | MN161849.1 | GU292316.1* | KU258875.1* |
| | MN564751.1 | EF680301.1 | MZ931278.1* | GU292317.1* | KU258876.1* |
| | OR669623.1 | EF680302.1 | ON204347.1* | GU292318.1* | KU258877.1* |
| | | EF680303.1 | OR729406.1* | KC189895.1* | KU258878.1* |
| | | EF680304.1 | PP646154.1* | NC007015.1* | KU258879.1* |
| | | EF680305.1 | PP646155.1* | PP669280.1* | KU258880.1* |
| | | EF680306.1 | PP707037.1* | PP707036.1* | KU980910.1 |
| | | EF680307.1 | PP707038.1* | | KU980911.1 |
| | | EF680308.1 | PP707039.1* | | KU980912.1 |
| | | EF680309.1 | PP707040.1* | | LC762616.1* |
| | | EF680310.1 | PP707041.1* | | LC762617.1* |
| | | EF680311.1 | PP707042.1* | | MK164396.1* |
| | | EF680312.1 | PP707043.1* | | MK164400.1 |
| | | EF680313.1 | PP707044.1* | | MK210250.1* |
| | | EF680314.1 | PP707045.1* | | MK559457.1* |
| | | EF680315.1 | PP707046.1* | | MT254061.1* |
| | | EF680316.1 | PP707047.1* | | MT254062.1* |
| | | EF680317.1 | PP707048.1* | | MT254063.1* |
| | | EF680318.1 | PP707049.1* | | MT254064.1* |
| | | JQ995521.1 | PP707050.1* | | MT598195.1 |
| | | | PP707051.1* | | MZ931277.1 |
| | | | PP707052.1* | | NC_004994.2* |
| | | | PP707053.1* | | ON843769.1* |
| | | | PP707054.1* | | OQ989633.1* |
| | | | PP707055.1* | | OR024676.1 |
| | | | | | OR682176.1* |
| | | | | | OR965522.1* |
| | | | | | PP130116.1* |
| | | | | | PP130117.1* |
| | | | | | PP130118.1* |
| | | | | | PP682590.1 |
| | | | | | PP669281.1* |
| | | | | | PP707056.1* |
| | | | | | PP707057.1* |
| | | | | | PP707058.1* |
| | | | | | PP707059.1* |
| | | | | | PP707060.1* |
| | | | | | PP707061.1* |

* FL used for Identity plot
