## Supplemental Table 2 for "Unraveling viral identity: Avoiding the trap of endogenous sequences for viral surveillance of small ruminant oncogenic retroviruses"

| # | Virus | *Position* | F/R | Publication specificity # | Sequence (5' -> 3') | Predicted XRV? | Associated references |
|---|---|---|---|---|---|---|---|
| 1 | ENTV-1 | U3 | F | S | TCAGGAAGTCTTAAGAGCTTTTAG | yes | (Cousens et al., 1999) |
| 2 | ENTV-1 | U3 | F | S | GATGTCTTTTGGTTTTGCAACATG | yes | (Cousens et al., 1999) |
| 3 | ENTV-1 | U3 | F | S | ATGATCTCAAGTTACTTAACTTGC | yes | (Cousens et al., 1999) |
| 4 | ENTV-1 | U3 | F | S | TCAGGAAGTCTTAAGAGCTTTTGG | yes | (Ortín et al., 2003, 2004) |
| 5 | ENTV-1 | U3 | F | S | AATGTGTTTTGGTTTTGCAACATG | yes | (Ortín et al., 2003, 2004) |
| 6 | ENTV-1 | U3 | F | S | GGAAGTCTTAAGAGCTTTTGGA | yes | (Walsh et al., 2010) |
| 7 | ENTV-1 | U3 | F | S | GGGCTCAGATGTTTTTTGGTTATGCA | yes | (Walsh et al., 2010, 2013) |
| 8 | ENTV-1 | U3 | F | S | ATGATCTTAAGTCACCTAAGTTGCC | yes | (Walsh et al., 2010, 2013) |
| 9 | ENTV-1 | U3 | R | S | CAAAAAACATCTGAGCCCTATAA | yes | (Walsh et al., 2010) |
| 10 | ENTV-1 | U3 | R | S | CCATGTTGCAAAACCAAAACAC | yes | (Kane et al., 2004) |
| 11 | ENTV-1 | U3 | R | S | TTTTAAGCCATGTTGCAWAAC | no * | (Riocreux-Verney et al., 2024) |
| 12 | ENTV-1 | U3 | F | S | GAGGGTTAAGTCCTGGGAG | no | (Riocreux-Verney et al., 2024) |
| 13 | ENTV-1 | U3 | R | S | AAGCAAGTTAAGTAACTTGAGATC | yes | (Cousens et al., 1999; Ortín et al., 2003, 2004; Incili et al., 2025) |
| 14 | ENTV-1 | U3 | R | S | GACAGGGAGCTTAGGTACTTGC | no | (Jahns and Cousens, 2020) |
| 15 | ENTV-1 | U3 | R | S | TCCCAGGACTTAACCCTCCG | no | (Jahns and Cousens, 2020) |
| 16 | ENTV-1 | R | F | S | AGCAAGGATCAGCCATTCT | no * | (Walsh et al., 2016) |
| 17 | ENTV-1 | R | R | S | GGCTGATACCTTGCTTTATTG | no | (Walsh et al., 2016) |
| 18 | ENTV-1 | U5 | F | S | GATGCTCCGTTCTCTCCTTATA | no | (Walsh et al., 2014; de Cecco et al., 2019) |
| 19 | ENTV-1 | U5 | R | S | CCAGCACGGAACAAAAGTTACA | yes | (Eckstrand et al., 2013) |
| 20 | ENTV-1 | U5 | R | S | ACGGACAAAGTTACACCTG | yes | (Walsh et al., 2010) |
| 21 | ENTV-1 | *gag* | R | S | GGGACGCGACGAATGTAGG | yes | (Walsh et al., 2014; de Cecco et al., 2019) |
| 22 | ENTV-1 | *gag* | R | S | CCTTGAACATCTGTTTGGACC | no | (Walsh et al., 2010) |
| 23 | ENTV-1 | *gag* | F | S | AACAAGCCCGAAACAAGAGG | no | (Walsh et al., 2010) |
| 24 | ENTV-1 | *gag* | R | S | AGGGACGCGACGAATGTAGG | yes | (Walsh et al., 2010, 2013) |
| 25 | ENTV-1 | *gag* | R | S | CGCGACGAATGTAGGGAC | yes | (Riocreux-Verney et al., 2024) |
| 26 | ENTV-1 | *gag* | F | S | ATCCGTCCCTACATTCGTC | yes | (Fox et al., 2011) |
| 27 | ENTV-1 | *gag* | R | S | CCTTGAACATCTGTTTTGGACC | no | (Fox et al., 2011) |
| 28 | ENTV-1 | *pro* | R | S | CTGTTCTGGATTGGTAGTTTGG | no | (Cousens et al., 1999) |
| 29 | ENTV-1 | *pol* | F | S | ATGGGAGCCCTACAACCTGG | no | (Cousens et al., 1999) |
| 30 | ENTV-1 | *pol* | R | S | CACGAGGATACAAGGAGAAACC | no | (Cousens et al., 1999) |
| 31 | ENTV-1 | *pol* | R | S | AAGGAACACAGATAAGGGGA | no | (Walsh et al., 2010) |
| 32 | ENTV-1 | *pol* | F | S | TAACACCCAACTAGTAAAGCTG | no | (Walsh et al., 2010, 2016) |
| 33 | ENTV-1 | *pol* | R | S | GAACACAGATAAAGGGAGGC | no | (Walsh et al., 2016) |
| 34 | ENTV-1 | *env* | F | S | TCTTTTGGTTCCTTGCCTCA | no * | (Eckstrand et al., 2013) |
| 35 | ENTV-1 | *env* | F | S | CTTTTAGTTCCCTGCCTCAC | yes | (Walsh et al., 2010) |
| 36 | ENTV-1 | *env* | F | S | GAGAATTGAATTAATACATATG | yes | (Kane et al., 2004) |
| 37 | ENTV-1 | *env* | F | S | GCTTAGCCGTCCTAAAAGAG | no | (Ortín et al., 2003; Incili et al., 2025) |
| 38 | ENTV-1 | *env* | F | S | CTCACTCGAGRTTTAATTAAAGATTTCC | yes | (Jahns and Cousens, 2020) |
| 39 | ENTV-1 | *env* | F | S | CCTACAAATGAGARTTGAATTAATACATA | yes | (Jahns and Cousens, 2020) |
| 40 | ENTV-2 | U3 | R | S | TGTTTTATTGTGTCATAGTATA | yes | (Li et al., 2024b) |
| 41 | ENTV-2 | U3 | R | S | CTTGTTGTTTTATTGTGTCATAGTATATAT | yes | (He et al., 2017; Zhai et al., 2019) |
| 42 | ENTV-2 | U3 | R | S | TAAGACTTCCTGAGGGTGGCGGGACA | no | (Ye et al., 2019) |

| | | | | | | | |
|---|---|---|---|---|---|---|---|
| 43 | ENTV-2 | U3 | R | S | CACCGGATCCTTAYGTAATCRGATTTCCTG | yes | (Apostolidi et al., 2019; Li et al., 2025b) |
| 44 | ENTV-2 | U3 | R | S | TTTATTGTGCAGCAGTATATATA | no | (Ortín et al., 2003) |
| 45 | ENTV-2 | U3 | F | S | CCCTCAGGAAGTCTTAAAAGCTC | yes | (Ortín et al., 2003) |
| 46 | ENTV-2 | U3 | R | S | GATCTTATCTGCTTATTTTCAG | yes | (Ortín et al., 2003, 2007; Fox et al., 2011; Riocreux-Verney et al., 2024; Incili et al., 2025) |
| 47 | ENTV-2 | U3 | F | S | GCAAAATGCCAGGACCTTGG | yes | (Ortín et al., 2003, 2007; Fox et al., 2011; Riocreux-Verney et al., 2024) |
| 48 | ENTV-2 | U3 | F | S | CCCTCAGGAAGTCTTAAAAG | yes | (Ortín et al., 2003, 2007) |
| 49 | ENTV-2 | U3 | R | S | GATCCTTATGTAATCAGATTTCC | yes | (Riocreux-Verney et al., 2024) |
| 50 | ENTV-2 | U3 | R | S | TCCCAGGACTTAACCATTC | no | (张靖鹏 et al., 2021) |
| 51 | ENTV-2 | U3 | F | S | CAATCACCGGATCCTTACTTAATCG | yes | (潘启东 et al., 2021) |
| 52 | ENTV-2 | R | F | S | ACAAGGCATCAGCCATTTTGGTCTGATCCTCTCAACCCCA | no | (He et al., 2017; Ye et al., 2019; Zhai et al., 2019) |
| 53 | ENTV-2 | R | F | S | ACAAGGCATCAGCCATTTTGGTC | no | (Li et al., 2025b) |
| 54 | ENTV-2 | R | R | S | GCTGATGCCTTGTTGTTTTATTGT | yes | (Li et al., 2025b) |
| 55 | ENTV-2 | R | F | S | ACAAGGCATCAGCCATTTTGGT | no | (Li et al., 2024b) |
| 56 | ENTV-2 | U5 | R | S | TTTTGTCTGTCTCTTATTTTCTTAGCGGGGG | no | (潘启东 et al., 2021) |
| 57 | ENTV-2 | *PBS* | F | S | GTTTTCCTCGCCACTACTCTTG | no | (He et al., 2017; Zhai et al., 2019) |
| 58 | ENTV-2 | *gag* | R | S | AACCTCACCAAGTCGCTGC | no | (Li et al., 2024b) |
| 59 | ENTV-2 | *gag* | F | S | GGGGAACAAATTCGAACTCATTATACT | no | (Li et al., 2024b) |
| 60 | ENTV-2 | *gag* | F | S | CCTTGGTTCCCCAGAGAAGG | no | (Jamil et al., 2024) |
| 61 | ENTV-2 | *gag* | F | S | GTCCCTAAAAATGCGACCTT | yes | (He et al., 2021) |
| 62 | ENTV-2 | *gag* | R | S | GCGACTCCTGAGTTCTGTAAAACCAC | yes | (He et al., 2021) |
| 63 | ENTV-2 | *gag* | F | S | AAATGCGACCTTCCGATAATGATGA | no | (He et al., 2021) |
| 64 | ENTV-2 | *gag* | R | S | CTTCTGTAGCGGGGACATATTCTCA | no | (He et al., 2021) |
| 65 | ENTV-2 | *gag* | R | S | AGGAGGAGGAGCATCATAACCAGGCTCTGGGTCAGGAATA | yes | (He et al., 2017; Zhai et al., 2019) |
| 66 | ENTV-2 | *gag* | R | S | TACCCAATAAGCGTCGGATGAT | no | (He et al., 2017; Zhai et al., 2019) |
| 67 | ENTV-2 | *gag* | F | S | CACTCCTAATTTGTGCCCACG | no | (He et al., 2017; Zhai et al., 2019) |
| 68 | ENTV-2 | *gag* | R | S | GGGCGGAGGAGCAGCATAACCGGATTCTGGATCAGGAACA | yes | (Ye et al., 2019) |
| 69 | ENTV-2 | *gag* | F | S | GGGGAACAAATTCGAACTCATTATACTTTA | no | (Ye et al., 2019) |
| 70 | ENTV-2 | *gag* | F | S | TGGTCAACCTGGCCATCGAGCTGCAGTATG | no | (Ye et al., 2019) |
| 71 | ENTV-2 | *gag* | F | S | CACAAAGATCAGGTAATTCGGG | no | (Ortín et al., 2003) |
| 72 | ENTV-2 | *gag* | F | S | CACAAAAGATCAGGTAATTCGGG | no | (Ortín et al., 2003) |
| 73 | ENTV-2 | *gag* | R | S | GTAAATGGAGCGGTAGGACCATATTGTG | no | (Li et al., 2025b) |
| 74 | ENTV-2 | *gag* | F | S | ACAACCAGCGTTATTATGAATCACTGCC | no | (Li et al., 2025b) |
| 75 | ENTV-2 | *gag* | R | S | AGGAGGAGGAGGAGCATC | no | (Riocreux-Verney et al., 2024) |
| 76 | ENTV-2 | *gag* | F | S | TTTCTTGTGCTAAAAAACCCCAATCTT | no | (潘启东 et al., 2021) |
| 77 | ENTV-2 | *gag* | R | S | CTAAACCTAAATTAATCAATTTTCTT | no | (潘启东 et al., 2021) |
| 78 | ENTV-2 | *pro* | R | S | CCCATCTCAGGTGCTAGTATTGTAT | no | (Li et al., 2024b) |
| 79 | ENTV-2 | *pro* | F | S | GCAACCCTGCAGGTTCCAA | no | (Li et al., 2024b) |
| 80 | ENTV-2 | *pro* | R | S | ACTCCCATCTCAGGTGCTAGTACTGTATAG | no | (Ye et al., 2019) |
| 81 | ENTV-2 | *pro* | R | S | CGAAACCTTTATCTTGACGGTC | no | (Ortín et al., 2003) |
| 82 | ENTV-2 | *pro* | R | S | TTGCACCCAATAAGCGTCAGAT | no | (Li et al., 2025b) |
| 83 | ENTV-2 | *pro* | F | S | GCGGGACAGCGTATAGCTCAACTCC | no | (Li et al., 2025b) |
| 84 | ENTV-2 | *pro* | F | S | GCCTCCTATACAGTACTAGCACCTG | no | (Li et al., 2025c) |
| 85 | ENTV-2 | *pro* | R | S | GATATTAGCTTCCGCTCCTAATAGA | no | (Li et al., 2025c) |

| | | | | | | | |
|---|---|---|---|---|---|---|---|
| 86 | ENTV-2 | *pro* | PROBE | S | TGCCTCCAGGGACAGCTGGATTGCTC | no | (Li et al., 2025c) |
| 87 | ENTV-2 | *pro* | F | S | AAGATGGGGCAAAATATAAGGTTTAA | no | (潘启东 et al., 2021) |
| 88 | ENTV-2 | *pro* | R | S | AGTTCAAACCCTTGCTACCGGAGT | no | (潘启东 et al., 2021) |
| 89 | ENTV-2 | *pro* | R | S | CAATTTCCACTCTTCAAGGTATTGGC | no | (潘启东 et al., 2021) |
| 90 | ENTV-2 | *pol* | F | S | TCCACCCTTCCTGGTGCC | no | (Yu et al., 2014; 江锦秀 et al., 2017; Li et al., 2022, 2024a, 2025a) |
| 91 | ENTV-2 | *pol* | R | S | CACAAACATGCCCTCGTCCCC | no | (Yu et al., 2014; 江锦秀 et al., 2017; Li et al., 2022, 2024a, 2025a) |
| 92 | ENTV-2 | *pol* | F | S | CCCTATAATTATTTGGGTTTCTCCTTAT | no | (Ye et al., 2019; Li et al., 2024b) |
| 93 | ENTV-2 | *pol* | R | S | AGTGACGAGTTGATTCTCCAGTATAG | no | (Li et al., 2024b) |
| 94 | ENTV-2 | *pol* | R | S | TGTGGGTGCTCGAGGGGTA | no | (Li et al., 2024b) |
| 95 | ENTV-2 | *pol* | F | S | ATTGCTGATGAAAAAATT | no | (Li et al., 2024b) |
| 96 | ENTV-2 | *pol* | R | S | TATTGAGGGAGAACAAA | no | (Li et al., 2024b) |
| 97 | ENTV-2 | *pol* | F | S | GCAAATGATTGAAACTGT | no | (Li et al., 2024b) |
| 98 | ENTV-2 | *pol* | R | S | TGGGTATTATARRCACGAGGA | no | (Jamil et al., 2024) |
| 99 | ENTV-2 | *pol* | R | S | GGCCACTGATCGACCCATAC | no | (He et al., 2017; Zhai et al., 2019) |
| 100 | ENTV-2 | *pol* | F | S | GAAGAGGTTTGGGGTGTTTTCCCTAGGGACCTCTGATTCTCCTGTGAC | no | (He et al., 2017; Zhai et al., 2019) |
| 101 | ENTV-2 | *pol* | R | S | GTTTAAGACGTTGATGAGCTCGTTCTACAATCCCTTGTCCCTGTGGGT | no | (He et al., 2017; Zhai et al., 2019) |
| 102 | ENTV-2 | *pol* | F | S | AGAACGAGCTCATCAACGTCTTAAACATCAACT | no | (He et al., 2017; Zhai et al., 2019) |
| 103 | ENTV-2 | *pol* | R | S | TCTATTGATTGTAAGGCTGATCGTCCTTCT | no | (Ye et al., 2019) |
| 104 | ENTV-2 | *pol* | R | S | AGTGACGAGTTGATTCTCCAGTATGAAG | no | (Ye et al., 2019) |
| 105 | ENTV-2 | *pol* | F | S | CCTCAATTCTTTGTTCTCCCTCAATATG | no | (Ye et al., 2019) |
| 106 | ENTV-2 | *pol* | F | S | CCATGATGCATATGGGAGC | no | (Ortín et al., 2003) |
| 107 | ENTV-2 | *pol* | R | S | CTGGGTAGTATTAAGCACGAG | ND | (Ortín et al., 2003) |
| 108 | ENTV-2 | *pol* | F | S | GCGCGAATATGTCCAAAGAAGC | no | (Ortín et al., 2003) |
| 109 | ENTV-2 | *pol* | R | S | CTATCTGCTACTCCAGCTAAAC | no | (Ortín et al., 2003) |
| 110 | ENTV-2 | *pol* | R | S | GCCTCATGACGTCCCTTCGCCACAA | no | (Li et al., 2025b) |
| 111 | ENTV-2 | *pol* | F | S | GCACCCACAGGGGTTCTCTATCAAG | no | (Li et al., 2025b) |
| 112 | ENTV-2 | *pol* | R | S | TGGTTTTTTGCAGGAGGATCGCT | no | (Li et al., 2025b) |
| 113 | ENTV-2 | *pol* | F | S | TAGCCCTTCACCACATAATGCCTTG | no | (Li et al., 2025b) |
| 114 | ENTV-2 | *pol* | F | S | TTCACCACATAATCCTTG | no | (Riocreux-Verney et al., 2024) |
| 115 | ENTV-1/ENTV-2 | *pol* | F | S | CTCATGGCTTCCCTTCATACTG | no | (Cousens et al., 1999;Ortín et al., 2003) |
| 116 | ENTV-2 | *pol* | F | S | TATAGCGGAAGGAGTGGGCAAC | no | (潘启东 et al., 2021) |
| 117 | ENTV-2 | *pol* | F | S | TGAAGGGAGGCCATGAGAAAATG | no | (潘启东 et al., 2021) |
| 118 | ENTV-2 | *pol* | R | S | CATTCTTTACGGTTACAATTTAA | no | (潘启东 et al., 2021) |
| 119 | ENTV-2 | *pol* | R | S | TTCCATATAACCCGCAGGGGCAGGG | no | (潘启东 et al., 2021) |
| 120 | ENTV-2 | *env* | R | S | ACTATTGCCATGACCAAA | no | (Li et al., 2024b) |
| 121 | ENTV-2 | *env* | F | S | CTCCTTGGACTTTATGTCGAGC | no | (Li et al., 2024b) |
| 122 | ENTV-2 | *env* | F | S | ATGTACAACCTCCGGTGACG | no | (Sobhy et al., 2023) |
| 123 | ENTV-2 | *env* | R | S | AATGGGTGGATGCTGTCCTC | no | (Sobhy et al., 2023) |
| 124 | ENTV-2 | *env* | R | S | ATACACTGATGCAACTCGTGCTCGACAT | no | (Ye et al., 2019) |
| 125 | ENTV-2 | *env* | F | S | TAGACACTCCTTGGACTTTATG | no | (Ye et al., 2019) |
| 126 | ENTV-2 | *env* | F | S | GAGGCAAATTGAGGCGTTGAT | no | (Huang et al., 2019) |
| 127 | ENTV-2 | *env* | R | S | CCCGTTCTGCATTCGCTGTAG | no | (Huang et al., 2019) |
| 128 | ENTV-2 | *env* | F | S | GCAATCCATTCAAGCTGCTCATAC | no | (Apostolidi et al., 2019) |
| 129 | ENTV-2 | *env* | F | S | CCTAACCTTCATTCRTTATGGCARAGT | no | (Apostolidi et al., 2019; Li et al., 2025b) |

| | | | | | | | |
|---|---|---|---|---|---|---|---|
| 130 | ENTV-2 | *env* | Probe | S | TGTTTAGTTCCTTGCCTCCTTCGTGG | no | (Apostolidi et al., 2019; Li et al., 2025b) |
| 131 | ENTV-2 | *env* | F | S | AAAGAGAGGGGAGCTGCC | no | (Ortín et al., 2003) |
| 132 | ENTV-2 | *env* | R | S | CGCAGCCTCCCCACTTTGCATTC | no | (Ortín et al., 2003) |
| 133 | ENTV-2 | *env* | F | S | CCTTAGGATGGGATAAGGAAG | no | (Ortín et al., 2003) |
| 134 | ENTV-2 | *env* | R | S | GTATAGGCTATTGCCATGACC | no | (Ortín et al., 2003) |
| 135 | ENTV-2 | *env* | F | S | CTTATATCCACGAGCATGTGTC | no | (Ortín et al., 2003) |
| 136 | ENTV-2 | *env* | R | S | CCTTATCCCACGCAAAATCAG | no | (Ortín et al., 2003) |
| 137 | ENTV-2 | *env* | R | S | CAGAAGCAGTGACAGCAGTTGCTAT | no | (Li et al., 2025b) |
| 138 | ENTV-2 | *env* | F | S | TAGCCGCAAGGAAAGAAGTGTGAGC | no | (Li et al., 2025b) |
| 139 | ENTV-2 | *env* | F | S | ATGGGGACGAGGG | no | (Riocreux-Verney et al., 2024) |
| 140 | ENTV-2 | *env* | F | S | AGCTGCTCATACTGTGGATC | no | (Ortín et al., 2003; Incili et al., 2025) |
| 141 | ENTV-2 | *env* | F | S | GAGATTTCTTACACATGAGAGC | no * | (张靖鹏 et al., 2021) |
| 142 | ENTV-2 | *env* | F | S | GTCGAGCACGAGTTGCATCA | no | (黎家明 et al., 2023) |
| 143 | ENTV-2 | *env* | R | S | GGTGGATGCTGTCCTCGGTA | no | (黎家明 et al., 2023) |
| 144 | ENTV-2 | *env* | PROBE | S | TGGGCACCCGGAGGAACACCTGATTTC | no | (黎家明 et al., 2023) |
| 145 | ENTV-2 | *env* | F | S | AGGCAGCTCACTCGTAGGCTCG | no | (潘启东 et al., 2021) |
| 146 | ENTV-2 | *env* | R | S | CAGTCCATTCAAGCTGCTCATACTG | no | (潘启东 et al., 2021) |
| 147 | ENTV-2 | *env* | F | S | TGGTACGATGAGACTGCCTTAGAG | no | (Li et al., 2024c) |
| 148 | ENTV-2 | *env* | R | S | CACTTTCGTGACATTATATGACAGG | no | (Li et al., 2024c) |
| 149 | ENTV-2 | *env* | PROBE | S | CCGCAAGGAAAGAAGTGTGAGCCTGATT | no | (Li et al., 2024c) |
| 150 | ENTV-2 | *ND* | F | S | AAAAAACCAACTTGTTGGGAACCCCCCTGTGTTTTATTGTACTGCA | ND | (潘启东 et al., 2021) |
| 151 | JSRV | U3 | F | S | AAGAATTTTTAAAAGCTCTTAAGG | no * | (DeMartini et al., 2001) |
| 152 | JSRV | U3 | R | S | ACAATGCTATATTTATAAAGTACA | yes | (DeMartini et al., 2001) |
| 153 | JSRV | U3 | Probe | S | AGCTCCCTGTCCCGCCACCCTC | no | (Bahari et al., 2016a, 2016b; Rosato et al., 2023) |
| 154 | JSRV | U3 | R | S | AGGGAGCTTAGGTACTTGTCC | no | (Zhang et al., 2014) |
| 155 | JSRV | U3 | R | S | CACCGGATTTTTACACAATCACCGG | yes | (Palmarini et al., 1996b, 1999a, 1999b; Holland et al., 1999; García-Goti et al., 2000; Gonzalez et al., 2001; Sanna et al., 2001; Ortín et al., 2004; Salvatori et al., 2004; Caporale et al., 2005; De Las Heras et al., 2005; Ortín et al., 2006, 2007; Cousens et al., 2007; Voigt et al., 2007a, 2007b; Grego et al., 2008; Lewis et al., 2011; Maeda et al., 2011; Humann-Ziehank et al., 2013; Minguijón et al., 2013; Azizi et al., 2014; Devi et al., 2014, 2016; Londoñe et al., 2014; Can-Sahna et al., 2015; Borobia et al., 2016, 2021; Sonawane et al., 2016; Singh et al., 2020; Mishra et al., 2021; Yadav et al., 2021, 2024; Consalter et al., 2022; Rosato et al., 2023; Valecha et al., 2023; Duan et al., 2024; Kumar et al., 2024; Patel et al., 2025; Rather et al.,2025) |
| 156 | JSRV | U3 | R | S | CATAACCTGTTTATTCATCTCAA | yes | (Riocreux-Verney et al., 2024) |
| 157 | JSRV | U3 | R | S | CCCACCGGATTTTTACACAATCACCGGATTC | yes | (Yadav et al., 2024) |
| 158 | JSRV | U3 | F | S | CCCTCTAGGATTCTTGAAAG | no | (Palmarini et al., 1996b; Riocreux-Verney et al., 2024) |
| 159 | JSRV | U3 | R | S | CCGGATTCTTACATAATCAG | yes | (Riocreux-Verney et al., 2024) |
| 160 | JSRV | U3 | F | S | CTGCGGGGGACGAC | no | (Riocreux-Verney et al., 2024) |
| 161 | JSRV | U3 | R | S | CTGCGGGGGACGACC | no | (Eckstrand et al., 2013) |
| 162 | JSRV | U3 | R | S | CTTAAGAGCCTTTAAAAATTCTTG | no * | (Riocreux-Verney et al., 2024) |
| 163 | JSRV | U3 | F | S | GAATCCGGTGATTGTGTAAAAATCCGGTGGG | yes | (Yadav et al., 2024) |
| 164 | JSRV | U3 | R | S | GGATTTTTACACAATCACC | no | (Riocreux-Verney et al., 2024) |
| 165 | JSRV | U3 | PROBE | S | GGCACTGCTTCACAGAAATATCAGGAAATCTGATT | yes | (Palmarini et al., 1996b; Sanna et al., 2001; Can-Sahna et al., 2015) |

| | | | | | | | |
|---|---|---|---|---|---|---|---|
| 166 | JSRV | U3 | R | S | TGATATTTCTGTGAAGCAGTGCC | yes | (Palmarini et al., 1996b, 1999a; Holland et al., 1999; García-Goti et al., 2000; Gonzalez et al., 2001; Ortín et al., 2004; Caporale et al., 2005; De Las Heras et al., 2005; Ortín et al., 2006, 2007; Voigt et al., 2007a, 2007b; Cousens et al., 2009, 2015; Lewis et al., 2011; Maeda et al., 2011; Martineau et al., 2011; Paethaisong et al., 2012; Humann-Ziehank et al., 2013; Minguijón et al., 2013; Azizi et al., 2014; Devi et al., 2014, 2016; Londoñe et al., 2014; Borobia et al., 2016, 2021; Suknao et al., 2016; Lee et al., 2017, 2019; Toma et al., 2020; Yadav et al., 2021, 2024; Consalter et al., 2022; Rosato et al., 2023; Hodor et al., 2024; Kumar et al., 2024; Rather et al., 2025) |
| 167 | JSRV | U3 | F | S | TGGGAGCTCTTTGGCAAAAGCC | yes | (Palmarini et al., 1996a, 1996b, 1999a, 1999b; Holland et al., 1999; García-Goti et al., 2000; Rosati et al., 2000; Gonzalez et al., 2001; Sanna et al., 2001; Ortín et al., 2004; Salvatori et al., 2004; Caporale et al., 2005; De Las Heras et al., 2005; Ortín et al., 2006, 2007; Cousens et al., 2007, 2009, 2015; Voigt et al., 2007a, 2007b; Grego et al., 2008; Lewis et al., 2011; Maeda et al., 2011; Martineau et al., 2011; Paethaisong et al., 2012; Humann-Ziehank et al., 2013; Minguijón et al., 2013; Devi et al., 2014, 2016; Londoñe et al., 2014; Can-Sahna et al., 2015; Borobia et al., 2016, 2021; Sonawane et al., 2016; Suknao et al., 2016; Lee et al., 2017, 2019; Singh et al., 2020; Toma et al., 2020; Mishra et al., 2021; Yadav et al., 2021, 2024; Consalter et al., 2022; Ortega et al., 2023; Rosato et al., 2023; Valecha et al., 2023; Duan et al., 2024; Hodor et al., 2024; Kumar et al., 2024; Patel et al., 2025; Rather et al., 2025) |
| 168 | JSRV | U3 | PROBE | S | TTTTTAAAAGCTCTTAAGGCTCGGATGTTTGCTT | yes | (Palmarini et al., 1996b; Sanna et al., 2001) |
| 169 | JSRV | U3 | F | NS | CGTGAAGGGTTAAGTCCTGGGAGCTCTTTGGCA | no | (Bai et al., 1996) |
| 170 | JSRV | U3 | F | S | CTGGGAGCTCTTTGGCAAAAGCCAAAG | yes | (Bai et al., 1996) |
| 171 | JSRV | U3 | F | S | GCTCTTAAGGCTCGGATGTTTGCTGC | yes | (Bai et al., 1996) |
| 172 | JSRV | U3 | F | S | ATCCGGTGGGTGTAGTTTGAGATGAAT | yes | (Bai et al., 1996) |
| 173 | JSRV | U3 | F | S | GTTATGTAATGTACTGTATAAATATAGCAAAGT | yes | (Bai et al., 1996) |
| 174 | JSRV | U3 | F | S | AATGCCGGTGATTGTGTAAGAATCCGGTGGGT | yes | (Bai et al., 1996) |
| 175 | JSRV | U3 | F | S | CCTAAGCTCCCTGTCCCG | no | (Ndione et al., 2025) |
| 176 | JSRV | U3 | PROBE | S | AACATGTTGCAACACCGACA | no | (Ndione et al., 2025) |
| 177 | JSRV | U3 | PROBE | S | TGTGAATGTCAGAAGTCACGT | no | (Ndione et al., 2025) |
| 178 | JSRV | U3 | R | S | AGCTCCCAGGACTTAACCCTTCACG | no | (Liu et al., 2006) |
| 179 | JSRV | U3 | R | S | TGGGAGCTCTTTGGCAGAAGCC | yes | (Bahari et al., 2016a, 2016b; Miller et al., 2017) |
| 180 | JSRV | U3 | F | S | CACCGGATTCTTACACAATCACCGG | yes | (Bahari et al., 2016a, 2016b) |
| 181 | JSRV | U3 | PROBE | S | AGCAAACATCCGARCCTTAAGAGCTTTCAAAA | yes | (Cousens et al., 2007, 2009, 2015; Martineau et al., 2011; Paethaisong et al., 2012; Suknao et al., 2016) |
| 182 | JSRV | U3 | F | S | GCCTAGGACAAGTACCTAAGCTC | no | (Miller et al., 2017) |
| 183 | JSRV | U3 | PROBE | S | AGCAAACATCCGAGCCTTAAGAGCTTTC | yes | (Lewis et al., 2011) |
| 184 | JSRV | U3 | PROBE | S | AGCAAACATGGCARCCTTAAGAGCTTTCAAAA | yes | (Lee et al., 2017, 2019) |
| 185 | JSRV | U3 | R | S | CACCGGCTTTTTACACAATCACCGG | yes | (Rosati et al., 2000) |
| 186 | JSRV | U3 | PROBE & R | S | ACCCACCGGATTCTTACACAATCACCGGCATT | yes | (Bai et al., 1996, 1999) |
| 187 | JSRV | U3 | PROBE & F | S | ATTCATCTCAAACTACACC | yes | (Bai et al., 1996, 1999) |
| 188 | JSRV | U3 | F | S | ATCCTCTCAACCCCATCTTTT | no | (Kumar et al., 2024) |
| 189 | JSRV | U3 | R | S | TTCATTATTCACTACACCCACCGG | yes | (Kumar et al., 2024) |
| 190 | JSRV | U3 | F | S | GGGTTAAGTCCTGGGAGCTC | no | (Kumar et al., 2024) |

| | | | | | | | |
|---|---|---|---|---|---|---|---|
| 191 | JSRV | U3 | R | S | GGATTCTTACACAATCACC | yes | (Gomes et al., 2017) |
| 192 | JSRV | R | F | S | GCAAGGTATCAGCCGTTCTGGTCTGATC | no | (Yadav et al., 2024) |
| 193 | JSRV | R | R | S | CACATGCTGATACCTTGCTT | no | (Liu et al., 2006) |
| 194 | JSRV | R | F | S | GCAGAGTATCAGCCATTTTG | no | (Kumar et al., 2024) |
| 195 | JSRV | U3-R | R | S | GCCTTCCTTTATTGTGCTGC | no | (Ndione et al., 2025) |
| 196 | JSRV | U3-R | R | S | GCTGATACTCTGCTTTATTACAATG | yes | (Kumar et al., 2024) |
| 197 | JSRV | R-U5 | F | S | GCATTGTAATAAAGCAGAGTATCAGCC | yes | (Palmarini et al., 1999a) |
| 198 | JSRV | R-U5 | F | S | GCAAGGTATCAGCCGTTCTG | no | (Liu et al., 2005, 2006) |
| 199 | JSRV | U5 | F | S | CTGCCGCGGCCACG | no | (Eckstrand et al., 2013) |
| 200 | JSRV | U5 | R | S | GCTGATACTCTGCTTTATTACAATGCTATA | yes | (Yadav et al., 2024) |
| 201 | JSRV | U5 | R | NS | GCACAAACAAGAGTCGCACCTGCACAGGGAG | no | (Bai et al., 1996) |
| 202 | JSRV | *PBS* | F | NS | CAACGTGGGGCTCGAGCTCGACAGTTTTCTTC | no | (Bai et al., 1996) |
| 203 | JSRV | *PBS* | R | S | CGAGCTCGAGCCCCACGTTG | no | (Ortín et al., 2004) |
| 204 | JSRV | *gag* | R | S | GAAGGGTGCATTTTCAGAGATGG | yes | (Riocreux-Verney et al., 2024) |
| 205 | JSRV | *gag* | R | S | GGAACCAAGGGCAAACTTCCTCAATAAATGAA | no | (Palmarini et al., 1999a) |
| 206 | JSRV | *gag* | F | NS | ACAGGCATGGAAAAAACTTCCTAGCTCCAGTAC | no | (Bai et al., 1996, 1999) |
| 207 | JSRV | *gag* | R | NS | TCTTGTTCCGGGCTTGCTGCTGGAAAAGTACT | no | (Bai et al., 1996) |
| 208 | JSRV | *gag* | R | NS | TTGAGGGCATACTGCAGCTCGATGGCCAGG | no | (Bai et al., 1996) |
| 209 | JSRV | *gag* | F | NS | AACTTTAGACACAGAAGGCAATTCAGCAGCCCA | no | (Bai et al., 1996, 1999) |
| 210 | JSRV | *gag* | R | S | CTCCCACCTTCTTCCATGTCTC | no | (Ortín et al., 2004) |
| 211 | JSRV | *gag* | F | S | CCCCATCTCTGAAAATGCAC | yes | (Fox et al., 2011; Linnerth-Petrik et al., 2014) |
| 212 | JSRV | *gag* | R | S | TGTTTAGACGGTGGAGGAAA | yes | (Fox et al., 2011; Linnerth-Petrik et al., 2014) |
| 213 | JSRV | *gag* | R | S | TCCAGGCAGTCACGAATCAA | no | (Liu et al., 2005, 2006) |
| 214 | JSRV | *gag* | F | S | TGATTCGTGACTGCCTGGT | no | (Liu et al., 2006) |
| 215 | JSRV | *gag* | R | S | GAGCCGCATTCGATATTTGT | no | (Liu et al., 2004, 2005, 2006) |
| 216 | JSRV | *gag* | F | S | CAGACCTCCTATGAAATGTT | no | (Liu et al., 2005, 2006) |
| 217 | JSRV | *gag* | R | S | CTGACGACTATGCGTTTGTCCC | no | (Miller et al., 2017) |
| 218 | JSRV | *gag* | F | NS | GCTGCTTTGAGACCTTATCGAAA | no | (Rosati et al., 2000; Singh et al., 2020; Patel et al., 2025) |
| 219 | JSRV | *gag* | R | NS | ATACTGCAGCTCGATGGCCAG | no | (Rosati et al., 2000; Singh et al., 2020; Patel et al., 2025) |
| 220 | JSRV | *gag* | R | S | CGGAGCAGAGGGAAAGGTATTAT | yes | (Kumar et al., 2024) |
| 221 | JSRV | *gag* | F | S | CCTCATAATAATACCTTTCCCTCTG | yes | (Kumar et al., 2024) |
| 222 | JSRV | *gag* | R | S | TACCATTCCGGCTTTCTCAT | no | (Kumar et al., 2024) |
| 223 | JSRV | *gag* | F | S | TTGTGGCACGACTTTTGGATAC | no | (Kumar et al., 2024) |
| 224 | JSRV | *gag* | R | S | CTAGGGAACCAAGGGCAAAC | no | (Kumar et al., 2024) |
| 225 | JSRV | *gag* | F | S | TGATTCGTGACTGCCTGGAT | no | (Liu et al., 2004, 2005) |
| 226 | JSRV | *pro* | R | S | CCTGTATAATCGGAGTCAAT | no | (Liu et al., 2005, 2006) |
| 227 | JSRV | *pro* | F | S | CCAGGGACAGTTGGATTACT | no | (Liu et al., 2006) |
| 228 | JSRV | *pro* | R | S | GCCTCGGTAACATTTTGCAC | no | (Kumar et al., 2024) |
| 229 | JSRV | *pro* | F | S | TTACCGAGGCACGACCAGAA | no | (Kumar et al., 2024) |
| 230 | JSRV | *pro* | F | S | TCTTGCTACGGGAGTGTTTG | no | (Kumar et al., 2024) |
| 231 | JSRV | *pro* | R | S | AGATGGGGCAGAATATAGGGTTTA | no | (Kumar et al., 2024) |
| 232 | JSRV | *pol* | F | S | TTCAGCAGCCCAGCGATTT | no | (Zhang et al., 2014) |
| 233 | JSRV | *pol* | R | S | ACAACCTCCATTTACCAGAC | no | (Liu et al., 2006) |
| 234 | JSRV | *pol* | F | S | GTAAGGTAAATGAGACGATGAT | no | (Liu et al., 2006) |
| 235 | JSRV | *pol* | R | S | CACGAATATGTCCAAAGAAA | no | (Liu et al., 2006) |

| | | | | | | | |
|---|---|---|---|---|---|---|---|
| 236 | JSRV | *pol* | F | S | AATTATCGGCACCACCTCTCCTGA | no | (Liu et al., 2006) |
| 237 | JSRV | *pol* | R | S | GCGGCGCTTCGGCATCC | no | (Liu et al., 2006) |
| 238 | JSRV | *pol* | F | S | ACAGGATGCCGAAGCGCCGCG | no | (Liu et al., 2006) |
| 239 | JSRV | *pol* | F | S | CAAGGCAATCACACTGCGGACGTT | no | (Bai et al., 1999) |
| 240 | JSRV | *pol* | F | S | CCAATCTGTGGTATGGGCCAGA | no | (Bai et al., 1999) |
| 241 | JSRV | *pol* | R | S | CTCCATTGATAGCGTTGCATAGG | no | (Kumar et al., 2024) |
| 242 | JSRV | *pol* | F | S | CCCACTCCTTCTGCTATACCTGAT | no | (Kumar et al., 2024) |
| 243 | JSRV | *pol* | R | S | AGGCATGAGAGCTAGCAAATTG | no | (Kumar et al., 2024) |
| 244 | JSRV | *pol* | F | S | TTGCAAATTACCCGGGACAGAT | no | (Kumar et al., 2024) |
| 245 | JSRV | *pol* | R | S | AGAGCATGGTTCAAGGCGTTAT | no | (Kumar et al., 2024) |
| 246 | JSRV | *pol* | F | S | CCCATCAACGCCTTAAACATCAA | no | (Kumar et al., 2024) |
| 247 | JSRV | *env* | F | S | ATGCCGAAGCGCCGCGCT | no | (Duan et al., 2024) |
| 248 | JSRV | *env* | R | S | CAGCTATTTCAACGGGCAGC | no | (Abass and Khudhair, 2022) |
| 249 | JSRV | *env* | F | S | GACCCCTCGACATTCCGTTT | no | (Abass and Khudhair, 2022) |
| 250 | JSRV | *env* | F | S | ATACGGGAACGGATCTGGACC | no * | (Walsh et al., 2010; Linnerth-Petrik et al., 2014) |
| 251 | JSRV | *env* | R | S | CAACATGAATGGATACGGCACGC | no | (Walsh et al., 2010; Linnerth-Petrik et al., 2014) |
| 252 | JSRV | *env* | R | S | TAGTTCTATATTTCATATGTAGCA | yes | (DeMartini et al., 2001) |
| 253 | JSRV | *env* | R | S | TCACGGGTCGTCCCCCGCATCTCCCCT | no | (Duan et al., 2024) |
| 254 | JSRV | *env* | F | S | TGGAAAACCCTGATCGGTCTAGGA | yes | (DeMartini et al., 2001) |
| 255 | JSRV | *env* | F | S | TTGCCTTGTTCGTGGCATG | no | (Riocreux-Verney et al., 2024) |
| 256 | JSRV | *env* | F | S | CGGTTCTGACTGTTGTGCTT | no | (Ndione et al., 2025) |
| 257 | JSRV | *env* | R | S | CGCAGCTCCCCTCTCTTTAT | no | (Ndione et al., 2025) |
| 258 | JSRV | *env* | F | S | CCGGAAAGAGATCGTACCGT | no | (Mansour et al., 2019; Al-Husseiny et al., 2020; Coskun et al., 2024) |
| 259 | JSRV | *env* | R | S | TAAGGAACACAAGCTCGGGG | no | (Mansour et al., 2019; Al-Husseiny et al., 2020; Coskun et al., 2024) |
| 260 | JSRV | *env* | F | S | TACCGTCGATAATTTCTTGC | no | (Liu et al., 2006) |
| 261 | JSRV | *env* | F | S | TTGGTGTAGGAATACTTGTGTT | yes | (Hudachek et al., 2010) |
| 262 | JSRV | *env* | R | S | TATTTCTATATTTCATATGCAGCA | yes | (Hudachek et al., 2010) |
| 263 | JSRV | *env* | PROBE | S | CTCGTTCGTGGCATGGTTCG | yes | (Hudachek et al., 2010) |
| 264 | JSRV | *env* | R | S | GCAGCCAAAAGTTTCCATAGTTCC | no | (Kumar et al., 2024) |
| 265 | JSRV | *env* | F | S | ACAGCATCCGCCTATTTTCTCT | no | (Kumar et al., 2024) |
| 266 | JSRV | *env* | R | S | CCACGAACAAGGCAAGGAAATATA | yes | (Kumar et al., 2024) |
| 267 | JSRV | *env* | F | S | TCGCTGTGGAAAACCCTGATTG | yes | (Kumar et al., 2024) |
| 268 | JSRV | *env* | F | S | GAGTTGAAATGCTGCATATG | yes | (Gomes et al., 2017) |
| 269 | JSRV | *env* | F | S | TTGGCTGCTTTTGGTCATGG | no | (Jassim et al., 2017) |
| 270 | JSRV | *env* | R | S | GGCCTTGTATCAACATGAATGGG | no | (Jassim et al., 2017) |
| 271 | JSRV | *env* | PROBE | S | TGGGAGCAAATATGGTGATGTGGGA | no | (Jassim et al., 2017) |
| 272 | JSRV | *env* | F | S | TGGAAAACCCTGATTGGTG | yes | (Grego et al., 2008) |
| 273 | JSRV | *env* | F | S | CATGGTTCGCGACTTTCTAAAG | yes | (Grego et al., 2008) |
| 274 | JSRV | *Orf-X* | F | S | ATCGGCACCACCTCTCCTGAA | no | (Rosati et al., 2000) |
| 275 | JSRV | *Orf-X* | R | S | GGCGTTATGTGGTGAGGGGCT | no | (Rosati et al., 2000) |
| 276 | JSRV/ERV/ENTV-2/ENTV-1 | *gag* | R | NS | ATACTGCAGCYCGATGGCCAG | no | (Cousens et al., 1996, 1999; Palmarini et al., 1996a, 1996b; Grossman et al., 2002; Ortín et al., 2003; Kane et al., 2004; Maeda et al., 2011; Shi et al., 2021; Ortega et al., 2023) |
| 277 | JSRV/ERV | *gag* | F | NS | GCTGCTTTRAGACCTTATCGAAA | no | (Cousens et al., 1996; Palmarini et al., 1996a; Grossman et al., 2002; Kane et al., 2004; Maeda et al., 2011; Shi et al., 202; Ortega et al., 2023) |

| | | | | | | | |
|---|---|---|---|---|---|---|---|
| 278 | JSRV/ENTV-1/ENTV-2 | *gag* | F | S | AAACAGACAGCTAGGGCGTG | no | (Hemida and Alnaeem, 2022) |
| 279 | ENTV-1/ENTV-2 | *gag* | R | S | GCTCGACAGAGGTCTGCAAT | no | (Hemida and Alnaeem, 2022) |
| 280 | JSRV/ENTV-1/ENTV-2 | *pro* | F | S | AATGTTACCGAGGCACGACC | no | (Hemida and Alnaeem, 2022) |
| 281 | JSRV/ENTV-1/ENTV-2 | *pro* | R | S | AGATCGAAAAGGCTTGGGGT | no | (Hemida and Alnaeem, 2022) |
| 282 | ENTV-1/JSRV | *pol* | F | S | GGAATGAACTGTATAGCCC | no | (Riocreux-Verney et al., 2024) |
| 283 | JSRV/ENTV-1/ENTV-2 | *pol* | F | S | GACGTCATGAGGCCATCCAA | no | (Hemida and Alnaeem, 2022) |
| 284 | JSRV/ENTV-1/ENTV-2 | *pol* | R | S | CAGCCCGAAAGTCCCATGAT | no | (Hemida and Alnaeem, 2022) |
| 285 | JSRV/ENTV-1/ENTV-2 | *env* | F | S | TGAGGCCACGAATGGACTAC | yes ** | (Hemida and Alnaeem, 2022) |
| 286 | JSRV/ENTV-1/ENTV-2 | *env* | R | S | CGACATTCCGTTTTGCGACA | no | (Hemida and Alnaeem, 2022) |

* Primer mapped on ERV sequences from discordant species (JSRV, ENTV-1 on goat ERV copeis and ENTV-2 on sheep ERV copies)
** Only specific for ENTV-1
### Specificity of the pair of primers described in the corresponding article(s)
