## Supplemental Table 3 for "Unraveling viral identity: Avoiding the trap of endogenous sequences for viral surveillance of small ruminant oncogenic retroviruses"

| Virus | Primer pair # | Target | 5'-3' sequence | Amplicon size (nt) | Reference | PCR enzyme | PCR conditions |
|---|---|---|---|---|---|---|---|
| ENTV | 01 | U5/ *gag* | ACAAGGCATCAGCCATTTTGGTC<br>GTAAATGGAGCGGTAGGACCATATTGTG | 1130 | (Li et al., 2025b)<br>(Li et al., 2025b) | Taq'Ozyme HS Mix | 35 x (95°C/ 15 sec; 55°C/ 15 sec; 72°C/ 100 sec) |
| | 02 | *gag* | GTCCCTAAAAATGCGACCTT<br>GCGACTCCTGAGTTCTGTAAAACCAC | 192 | (He et al., 2021)<br>(He et al., 2021) | Taq'Ozyme HS Mix | 35 x (95°C/ 15 sec; 56°C/ 15 sec; 72°C/ 30 sec) |
| | 03 | *gag/ pro* | ACAACCAGCGTTATTATGAATCACTGCC<br>TTGCACCCAATAAGCGTCAGAT | 1363 | (Li et al., 2025b)<br>(Li et al., 2025b) | Taq'Ozyme HS Mix | 35 x (95°C/ 15 sec; 53°C/ 15 sec; 72°C/ 100 sec) |
| | 04 | *gag/ pol* | CACTCCTAATTTGTGCCCACG<br>GGCCACTGATCGACCCATAC | 1087 | (He et al., 2017; Zhai et al., 2019)<br>(He et al., 2017; Zhai et al., 2019) | Taq'Ozyme HS Mix | 35 x (95°C/ 15 sec; 60°C/ 15 sec; 72°C/ 100 sec) |
| | 05 | *pro/ pol* | GCAACCCTGCAGGTTCCAA<br>TGTGGGTGCTCGAGGGGTA | 1867 | (Li et al., 2024b)<br>(Li et al., 2024b) | Taq'Ozyme HS Mix | 35 x (95°C/ 15 sec; 63°C/ 15 sec; 72°C/ 130 sec) |
| | 06 | *pol* | TCCACCCTTCCTGGTGCC<br>CACAAACATGCCCTCGTCCCC | 814 | (Yu et al., 2014; 江锦秀 et al., 2017; Li et al., 2022, 2024a, 2025a)<br>(Yu et al., 2014; 江锦秀 et al., 2017; Li et al., 2022, 2024a, 2025a) | Taq'Ozyme HS Mix | 35 x (95°C/ 15 sec; 63°C/ 15 sec; 72°C/ 60 sec) |
| | 07 | *pol* | CCCTATAATTATTTGGGTTTCTCCTTAT<br>AGTGACGAGTTGATTCTCCAGTATAG | 1372 | (Ye et al., 2019; Li et al., 2024b)<br>(Li et al., 2024b) | Taq'Ozyme HS Mix | 35 x (95°C/ 15 sec; 60°C/ 15 sec; 72°C/ 100 sec) |
| | 08 | *env*/ U3 | TAGACACTCCTTGGACTTTATG<br>TAAGACTTCCTGAGGGTGGCGGGACA | 1365 | (Ye et al., 2019)<br>(Ye et al., 2019) | Taq'Ozyme HS Mix | 35 x (95°C/ 15 sec; 58°C/ 15 sec; 72°C/ 100 sec) |
| | 09 | *env*/ U3 | CACCGGATCCTTAYGTAATCRGATTTCCTG<br>CCTAACCTTCATTCRTTATGGCARAGT | 399 | (Apostolidi et al., 2019; Li et al., 2025b)<br>(Apostolidi et al., 2019; Li et al., 2025b) | Taq'Ozyme HS Mix | 35 x (95°C/ 15 sec; 51°C/ 15 sec; 72°C/ 30 sec) |
| | 10 | U3/ *gag* | GCAAAATGCCAGGACCTTGG<br>AGGAGGAGGAGGAGCATC | 863 | (Ortín et al., 2003, 2007; Fox et al., 2011; Riocreux-Verney et al., 2024)<br>(Riocreux-Verney et al., 2024) | PrimeSTAR® GXL Premix | 35 x (98°C/ 10 sec; 63°C/ 15 sec; 68°C/ 70 sec) |
| | 11 | *gag/ pol* | CCTTGGTTCCCCAGAGAAGG<br>TGGGTATTATARRCACGAGGA | 3126 to 3206 | (Jamil et al., 2024)<br>(Jamil et al., 2024) | Taq'Ozyme HS Mix | 35 x (95°C/ 15 sec; 52°C/ 15 sec; 72°C/ 100 sec) |
| JSRV | 12 | U3RU5 | CTGCCGCGGCCACG<br>CTGCGGGGGACGACC | 397 | (Eckstrand et al., 2013)<br>(Eckstrand et al., 2013) | Taq'Ozyme HS Mix | 35 x (95°C/ 15 sec; 63°C/ 15 sec; 72°C/ 30 sec) |
| | 13 | *env* | TTCAGCAGCCCAGCGATTT<br>AGGGAGCTTAGGTACTTGTCC | 2054 | (Zhang et al., 2014)<br>(Zhang et al., 2014) | Taq'Ozyme HS Mix | 35 x (95°C/ 15 sec; 60°C/ 15 sec; 72°C/ 140 sec) |
| | 14 | *env* | ATGCCGAAGCGCCGCGCT<br>TCACGGGTCGTCCCCCGCATCTCCCCT | 1848 | (Duan et al., 2024)<br>(Duan et al., 2024) | Taq'Ozyme HS Mix | 35 x (95°C/ 15 sec; 65°C/ 15 sec; 72°C/ 130 sec) |
| | 15 | U3/ *gag* | CTGCGGGGGACGAC<br>GAAGGGTGCATTTTCAGAGATGG | 935 | (Riocreux-Verney et al., 2024)<br>(Riocreux-Verney et al., 2024) | Q5® Hot Start High-Fidelity DNA Polymerase | 35 x (98°C/ 10 sec; 65°C/ 15 sec; 72°C/ 40 sec) |
| | 16 | *env* | CGGTTCTGACTGTTGTGCTT<br>CGCAGCTCCCCTCTCTTTAT | 159 | (Ndione et al., 2025)<br>(Ndione et al., 2025) | Taq'Ozyme HS Mix | 40 x (95°C/ 15 sec; 52°C/ 15 sec; 72°C/ 30 sec) |
| | 17 | U3/ R | CCTAAGCTCCCTGTCCCG<br>GCCTTCCTTTATTGTGCTGC | 240 | (Ndione et al., 2025)<br>(Ndione et al., 2025) | Taq'Ozyme HS Mix | 40 x (95°C/ 15 sec; 52°C/ 15 sec; 72°C/ 30 sec) |
