## Supplementary figures and images for "Unraveling viral identity: Avoiding the trap of endogenous sequences for viral surveillance of small ruminant oncogenic retroviruses"

### Supplemental Figure 1

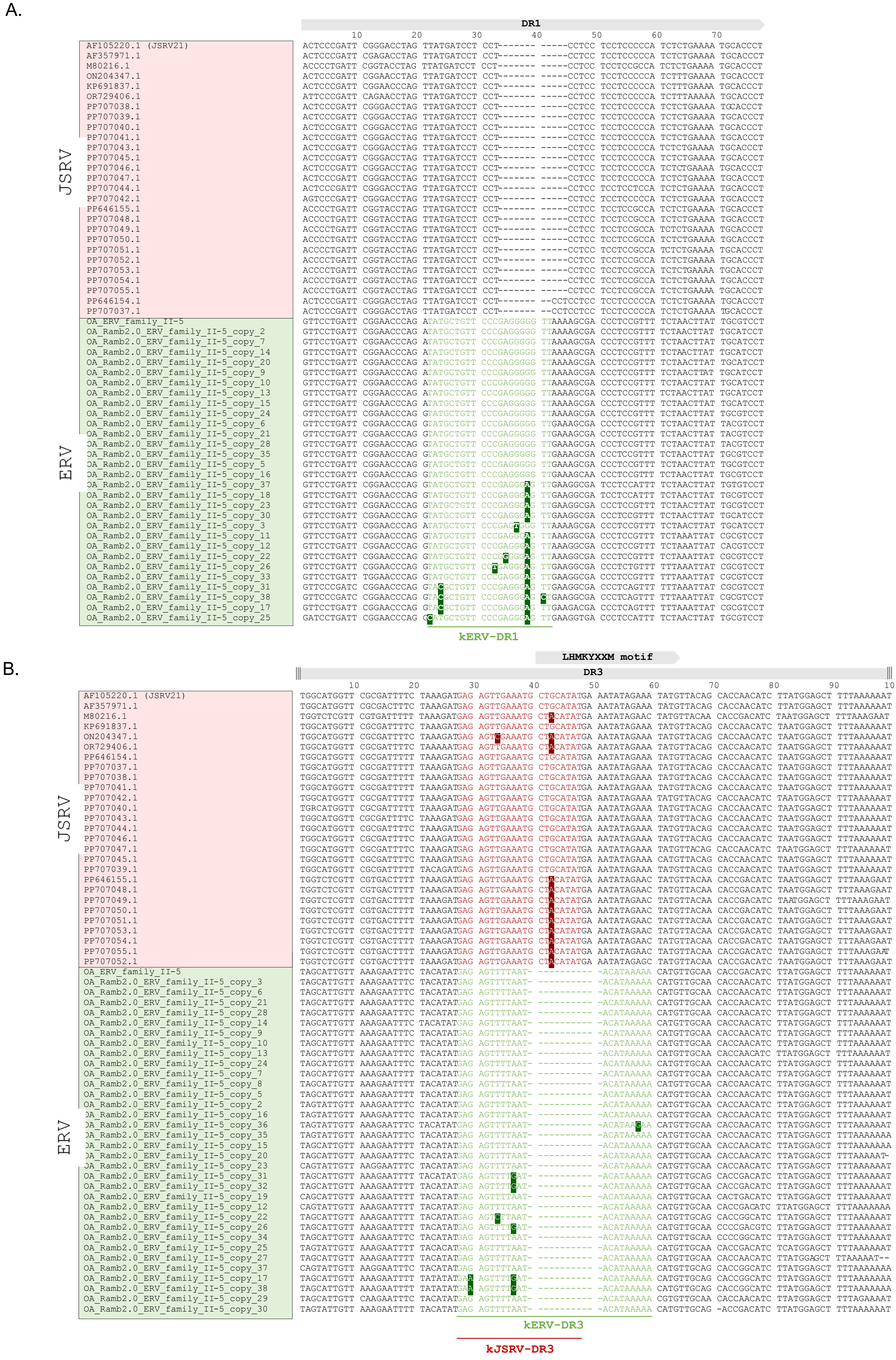


Supplementary Figure 1

C.

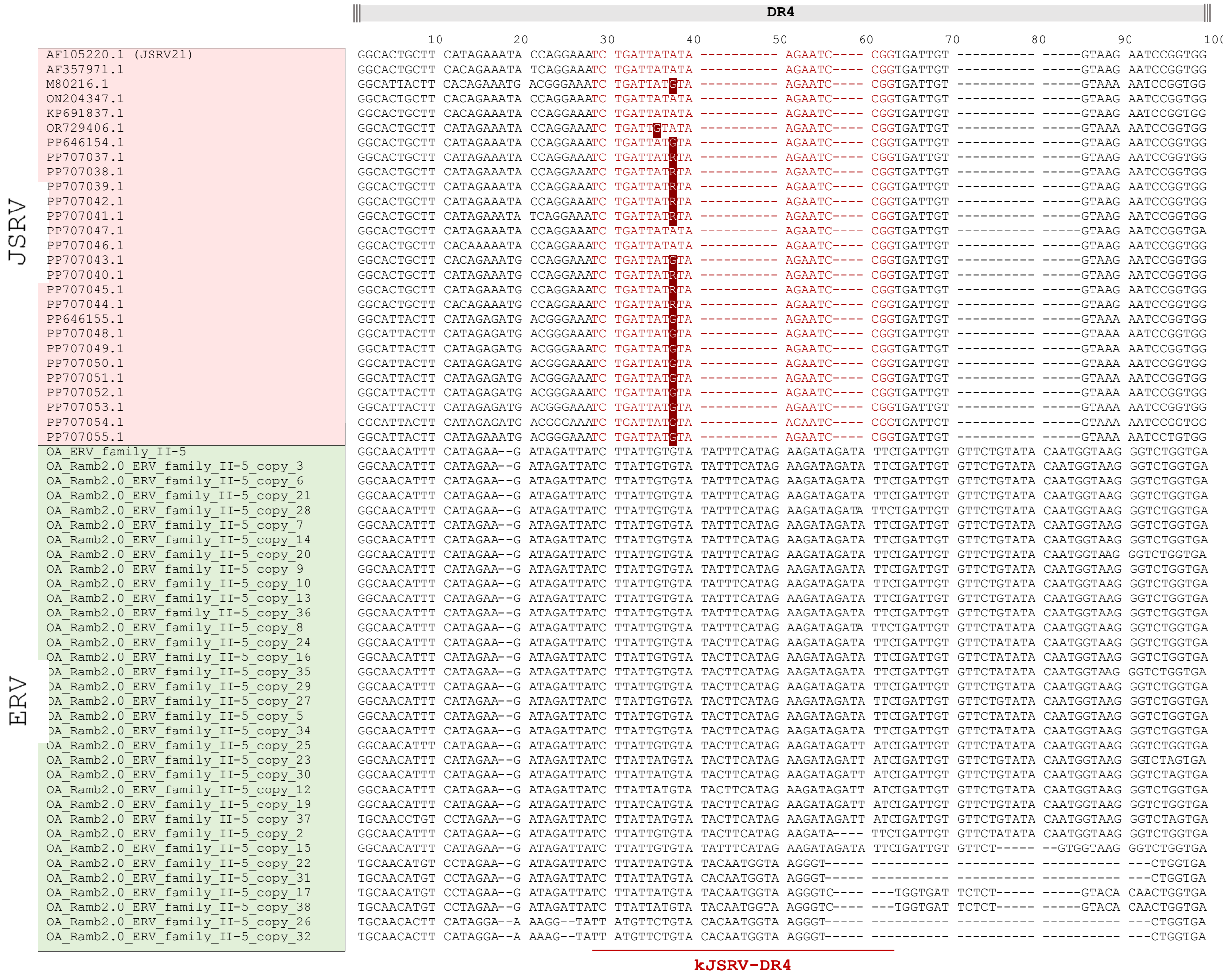


Supplementary Figure 1

### Supplemental Figure 2

A. JSRV

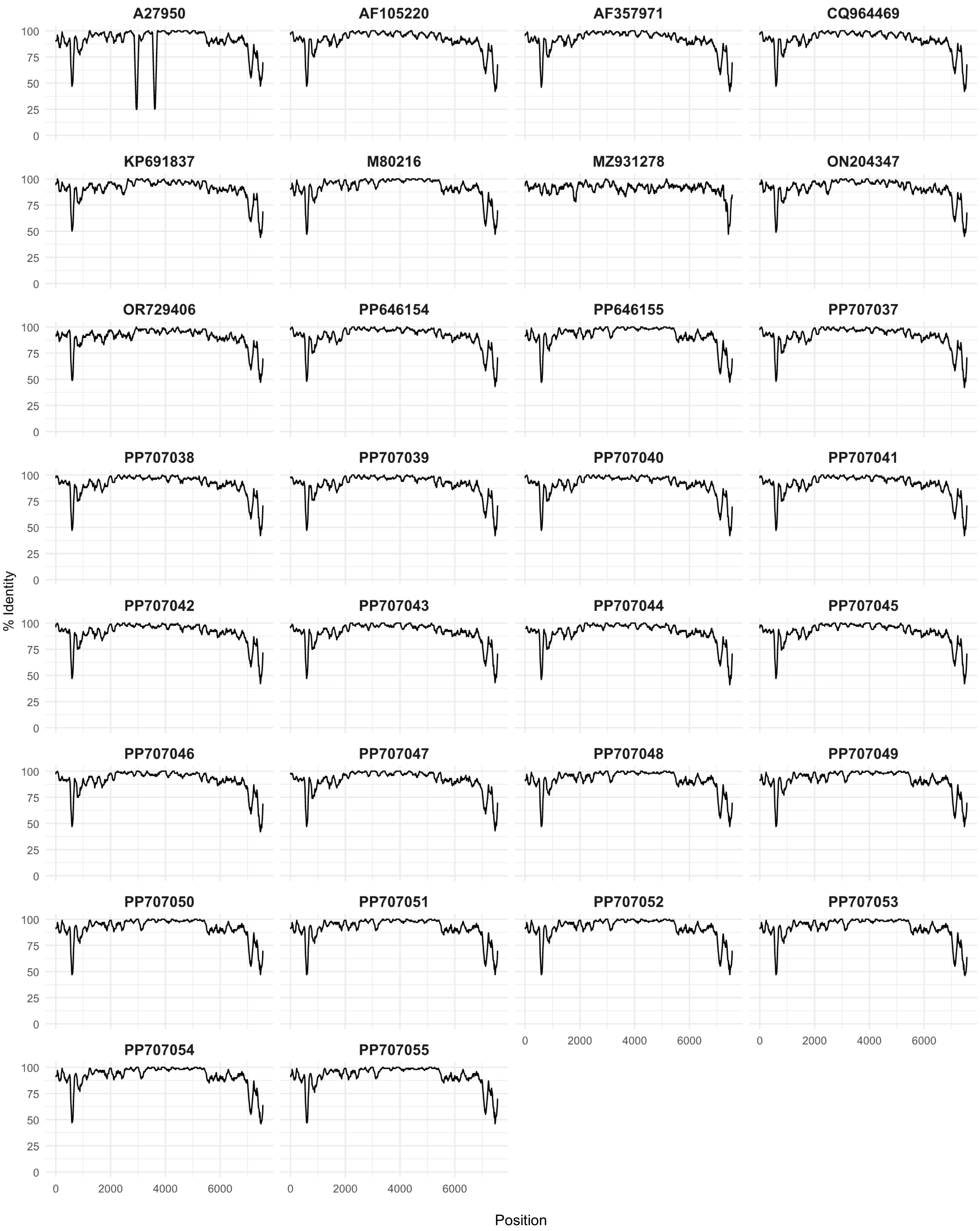


Supplementary Figure 2

B. ENTV-1

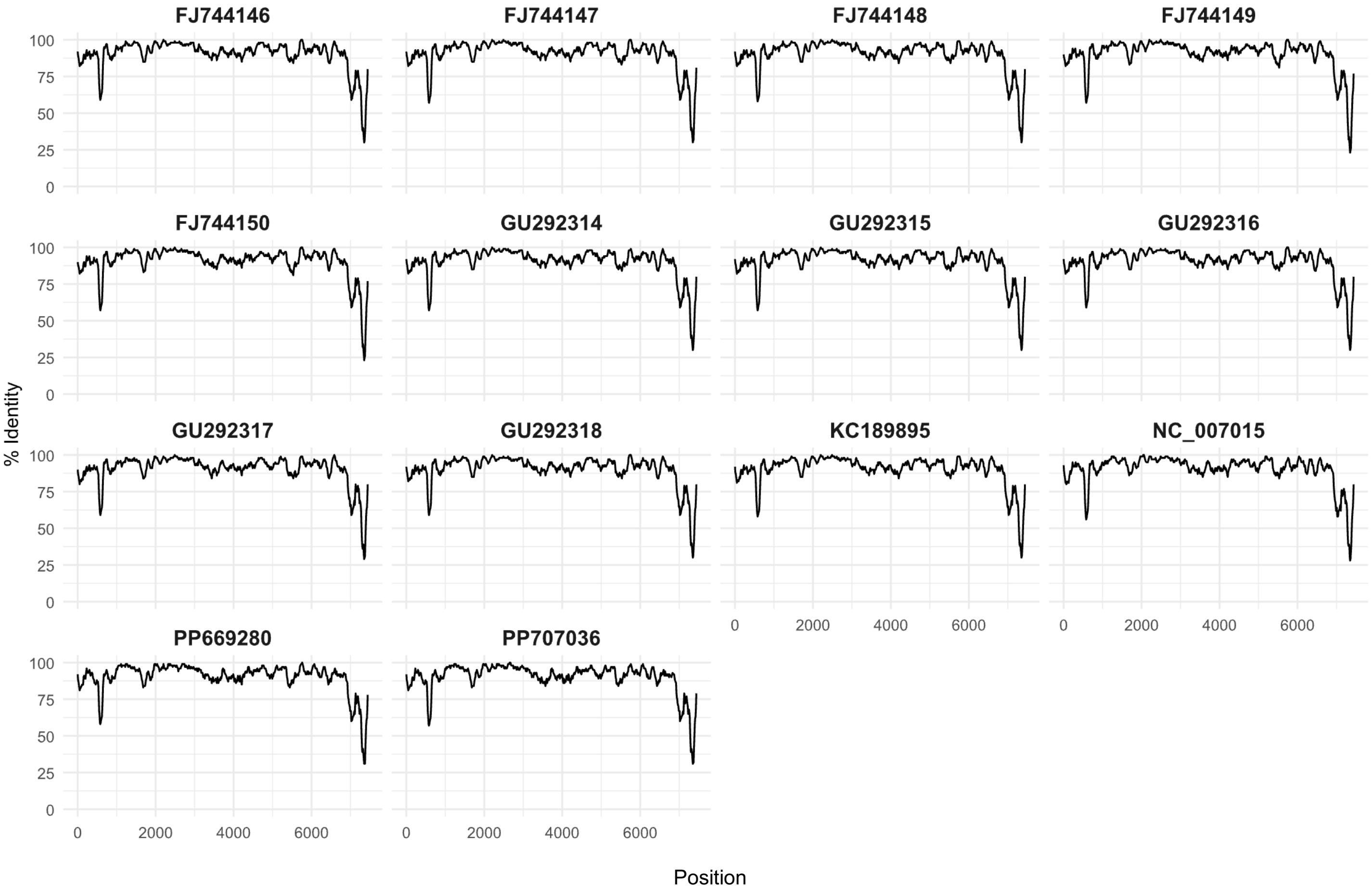


Supplementary Figure 2

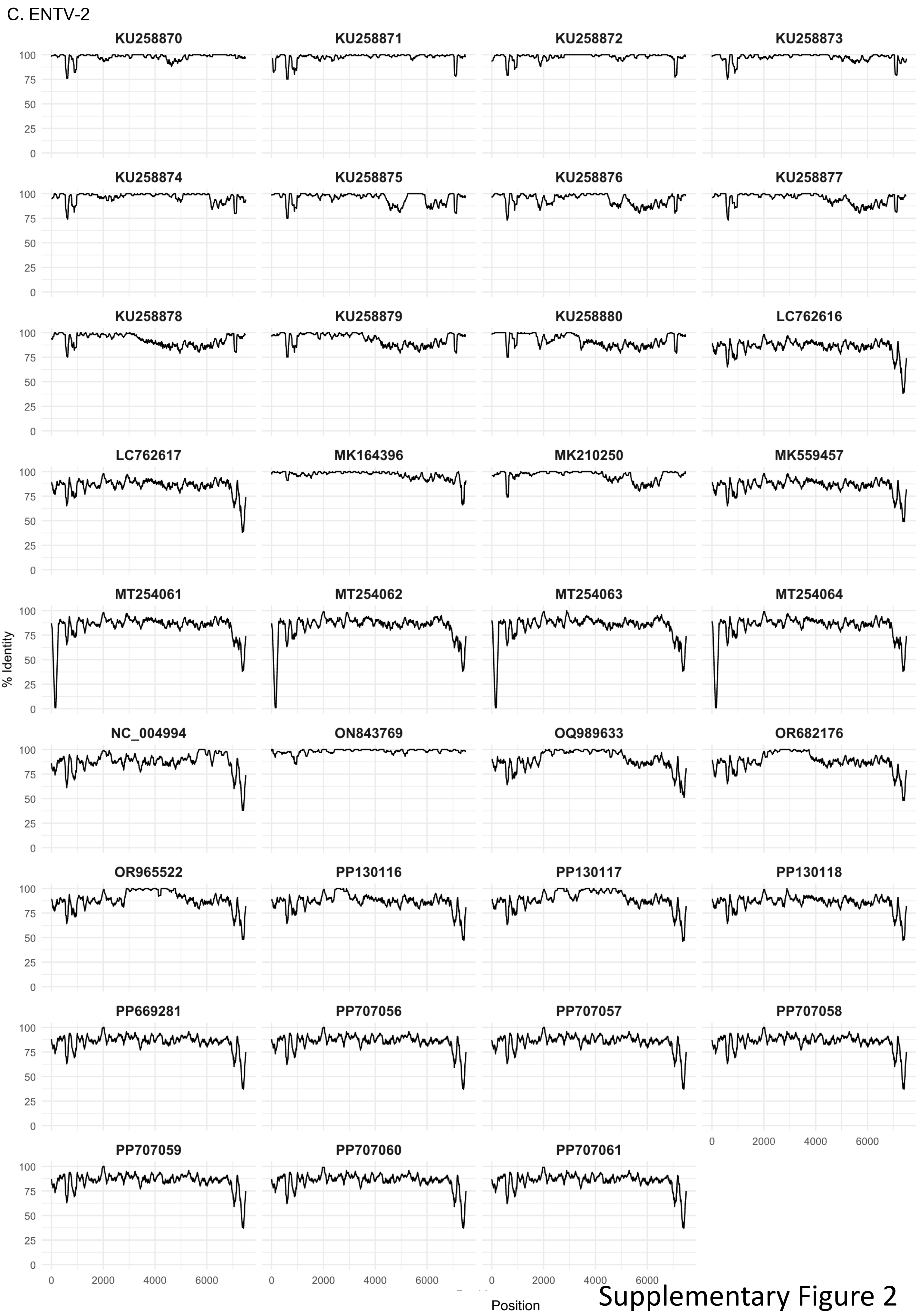


Supplementary Figure 2
