## Supplemental Figure 3 for "Unraveling viral identity: Avoiding the trap of endogenous sequences for viral surveillance of small ruminant oncogenic retroviruses"

## A. DR1

```
                    600       610       620       630       640       650       660       670
ERV II.5 Oa    GTTCCTGATTCGGAACCCAGATATGCTGTTCCCGAGGGGGTTAAAAGCGACCCTCCGTTTTCTAACTTATTGCGTCCT
ERV II.5 Ch    .........C.........G...........A...A...G..G.................A....C.......
ENTV1 FR4403   .A......C......A.T..T....A..C...T---------CCCCCTCCT.A.....CC.TAC.T.CG.C......C
ENTV2 FR3824   A.......CC.A..G..TG.T....A..C...T------------CCTCCT.....TCCG..CCTGAA......A...
JSRV FR2054    ACC..........T...T..T....A.CC...T------------CCTCCT......CCA...CTGAA.A...AC...
JSRV FR989     AC...C.......G...T..T....A.CC...T------------CCTCCT.....CCCA...CTGAA.A...AC...
```

## B. DR2

```
                         860       870       880       890       900       910       920       930       940       950       960       970       980
ERV II.5 Oa    AAGCTTCCTAGTAACAATGCCAAATCTCCTTCTGCTCCGCCTCTTCCCCCTGCTTATGCTCCTTCTGTTGTGGCTGGTCTCGATCCCCCTCCAGGG---------CCCCCTCCACCGTCTGAGAAC
ERV II.5 Ch    ........C.T..G...CT.T.....................G........C.........CA..A.A..A................G...CCTCCTCCT..T.....T..A.......TT
ENTV1 FR4403   ..A.......C...T....AT...CT.T.................T...........C..........C..A................G...---------...............A.A..T
ENTV2 FR3824   ...T..T...A...TC.C.T....C....C...........AT.C..A.......C.................C.....T..............---------..TT..TT....A.....T.
JSRV FR2054    C.............T..CAAT.CC.T...C..............T....T.........A....CA.....A.......C..T...........C...---------TTT.....G......A.AC..
JSRV FR989     TC......---.C.T...AAT.CC.T...C...........T....T..C......A...........A.......C..T........G..C...---------TTT............A.AC.T
```

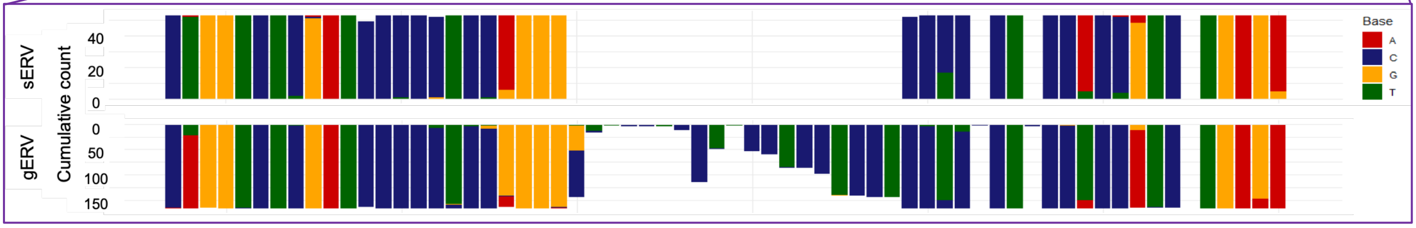


## C. DR3

```
                 7020      7030      7040      7050      7060      7070      7080      7090      7100      7110      7120      7130      7140
ERV II.5 Oa    CGAAGTATAATTGCTATGGGCGCGGTTCTGACTGTTGTGCTTATCATAATTTGTCTAGCTCCTTGTCTTATTCGTAGCATTGTTAAAGAATTTCTACATATGAGAGT------------TTTAATA
ERV II.5 Ch    ..T..C........G..........................AT.G......T..........C...........................T............------------.C.G...
ENTV1 FR4403   .A......T....T...A.CTATAA..G..TT.A.....A....TG.....CT.T...T......C..C.C...AG.TT.AA.......T..C..G..A......A.TGAATTAATACA.A.G.A.
ENTV2 FR3824   .AG......C.....G..ATTATAA..G...TC...A.AA....TG........T...T......C..CC.....G.TC.CT..CG...T..C.............CTGAATTGCTACA...G.A.
JSRV FR2054    AA..CCC.G..C.G.C.A..AATAT..G...TAA..A.TGC....G....C.T.G..TT...C...G.CG.....G.TC.C...CGT..C...T..A.G........TGAAATGCTACA.A.G.A.
JSRV FR989     AA..CCC.G....G.G.A..AATAC..G..TT.A..A.AA..G..G....CCT.A..TT......C...G.....G....G...CGC..T......A.G........TGAAATGCTGCA.A.G.A.
```

```
                     7150      7160      7170      7180
                H  K  N  M
ERV II.5 Oa    CATAAAAACATGTTGCAACACCGACATCTTATGGAGCTTTT-AAA
ERV II.5 Ch    ........................................A...
                Y  R  Y  M
ENTV1 FR4403   T...G.T..........T...A.A.A........T...G.---G
                Y  R  H  M
ENTV2 FR3824   T...G.C.T...A.....T.......G..A...........-.GG
                Y  R  T  M
JSRV FR2054    T...G..CT.....A..............A...........-...
                Y  R  N  M
JSRV FR989     T...G...T.....A..G....A......A...........-...
```

## D. DR4

```
                      7230      7240      7250      7260      7270      7280      7290      7300      7310      7320      7330      7340
ERV II.5 Oa    TGAAGGGTTAAGTCTTGGGAGCTCCCTGGCAGGTATGCCGGGCCCTAGGACACGTGCCTAAGCTCCCTGTCCCGCCACCCTCAAGAG-TTTTTATAACCCTTAAGGCTCCAAGATGTTTGGTTTCG
ERV II.5 Ch    .......................G.TC...-.T.....A.A..............T...................C....G....A---.....T..............GG..C..C.....CTT
ENTV1 FR4403   C.G...........C.........T....-AAA..A..A..G.......G..A..A...........................G..AG.C..A.G.G.TT..GG.....AG.TG.T...T.G..TT
ENTV2 FR3824   ..............C.....C....T....--AA.....A..A...T..GA.A..A.....A.......................G..AG.C..A.A.G.T....C.....--.A...A.C...G.T.
JSRV FR2054    ..............C.........TT....--AA.A...AAAG.T.......AA.A.........................T..GAT.C..G.A.G.T.......T..--G........C.GCT.
JSRV FR989     ..............C.........TT....--AR.A...AAAG.........A..A............................AT....A.AGG.T..........--G........C..CT.
```

```
                7350      7360      7370      7380      7390      7400      7410      7420      7430      7440      7450      7460      7470
ERV II.5 Oa    GCAACATTTCATAGAA---GATAGATTATCTTATTGTGTATATTTCATAGAAGATAGATATTCTGATTGTGTTCTGTATACAATGGTAAGGGTCTGGTGATTGTATCCTGAG-ATTAAAAAACAAC
ERV II.5 Ch    ................---.........G.......----------............TA...AT.....A......C....C...........T..........TTG...-......-G.....T
ENTV1 FR4403   .......GG.T..A..----GC..GAA....G...A...---------------------------------------------A.GAAAA.CA..A.....---------G.A.GT...TG.T
ENTV2 FR3824   ...TA.C.........AAGAGC..GAA....G...ACA.-----------------------------------------------....A..C...........T..A.AA..A.GC.G.T..G
JSRV FR2054    ...TT.C........GA-T..CG.GAA....G...A...--------------.AGA..CCGG.......---------------....AAA..C....GG....GTT....-..G..T....---
JSRV FR989     ...CTGC.........A-TACC..GAA....G...A.A.--------------.AGA..CCGG.......---------------.....AA..C.....G....GTGAATA-..G..T....---
```

```
                   7480      7490      7500      7510      7520
ERV II.5 Oa    CTTGTGAGTGCCTTAAGTCACGTACTTTACCCTATATATACCGCAGCAC
ERV II.5 Ch    .......A..T.AG...............T...........T.......
ENTV1 FR4403   ...-----------....T..C.GA...G.-..........T..TA...
ENTV2 FR3824   A.C-----------..........TAA...T..........TATG....
JSRV FR2054    --------------.G..T.T...A.G...TG....A...TA...A.GT
JSRV FR989     --------------....T.T..TG.A...TT....A...TA...TTGT
```

Supplementary Figure 3
