## Supplemental Figure 4 for "Unraveling viral identity: Avoiding the trap of endogenous sequences for viral surveillance of small ruminant oncogenic retroviruses"

```
                        10        20        30        40        50        60        70        80        90       100
HERVK13-int     AAATTGTCCTCGGCTTTCAGGCTGCCATTCTTTTCAATGCCAACAACAAAACCCTATACAGCAACAAATCTTGCCCTCCAGTCTGTGCCCACGATGCCGT
Amplicon HeLa   ----------------------------------------------------------------------------------------------------
                       110       120       130       140       150       160       170       180       190       200
HERVK13-int     AAGGGAAATCACTGGGCTGCTCAGTGCCGCGCAAAATTTGACATTGATGGTAACCCTTTAAAGCCATTCAGCAACTAGGGAAATGGAAAGAGGGGCCAGC
Amplicon HeLa   ----------------------------------------------------------------------------------------------------
                       210       220       230       240       250       260       270       280       290       300
HERVK13-int     CCCAGGCCCCTCCAAACAATGGGGAATTCCTCAACTCACAGCCTTTGGCAGCCAGCCAGATGAGCACTTTCCCAGCCCAATCCATTCAACCTGCCCCCCA
Amplicon HeLa   ----------------------------------------------------------------------------------------------------
                       310       320       330       340       350       360       370       380       390       400
HERVK13-int     GTTCCCACTTCAGCCATTTGTACCACAGGATCTAACGACTCAGTCCCAACACAAATCTCAGTACAATGCTTG-TCCCCTGCCACCACAGGCTCAGCCACT
Amplicon HeLa   ---------------.......................T.................................T.................T.........
                       410       420       430       440       450       460       470       480       490       500
HERVK13-int     GTAGATC-TCTGCTGCACTCGAGATATTTCTCTCTTACCCGGAGAGCCACCTATTGCTGTTTCCACAGGTGTGTTTGGCCGTTTGCCACCTGGCACTGTC
Amplicon HeLa   .......T............................................................................................
                       510       520       530       540       550       560       570       580       590       600
HERVK13-int     AGCTTGCTTCTTGGTCGCTCAAGCCTAAATTTAAAAGGTGTTCAAGTACATACTGGTGTAATCGACTCTGATTATTCAGGTGAAATTCACATTGTCATTA
Amplicon HeLa   ....................................................................................................
                       610       620       630       640       650       660       670       680       690       700
HERVK13-int     GCTCTGCAGTATCTTGGTATGCAGCAGCAGGAGACAGCATTGCTCAACTTCTTATACTCCGTTACATTCCACTAGGATCTAGTTCTTGTAAGAGAACCAG
Amplicon HeLa   ....................................................................................................
                       710       720       730       740       750       760       770       780       790       800
HERVK13-int     AGGTTTCAGAAGCACAGACTTTCAGGGCAAAACAGCCTATAGGGCCAGCAAGATTTCTGATACTCGTCCTGTGTGCTCCATGCATACATATTCAAGGAAG
Amplicon HeLa   ........................................T...........................................................
                       810       820       830       840       850       860       870       880       890       900
HERVK13-int     GAAGTTTGAGGGAATGATTGACTCGGGTGCTGATATCACCATTATAGCCTCATATCAATAGCCTAGACACTGGCCCAAAGAATATGCATCCACAGCCTTA
Amplicon HeLa   ......................A.............................................................................
                       910       920       930       940       950       960       970       980       990      1000
HERVK13-int     GTTGGTGTTGGTCAGGCTTCAGAGGTTTATGAAAATTCCACTATTTTACACTGTACGGGCCCAGAGGGACAGACTGGTACTGTTCACCCCCTCATTACGC
Amplicon HeLa   ....................................................................................................
                      1010      1020      1030      1040      1050      1060      1070      1080      1090      1100
HERVK13-int     CTATTCCAGTTAATCTTTGGGGAAGAGATCTTTTACAACAATGGAGGACACAGATTTCTTTCCCACAAGGTAATTACAGCTAACAAAGTAAAGACATTAT
Amplicon HeLa   ..........G.........................................................................................
                      1110      1120      1130      1140      1150      1160      1170      1180      1190      1200
HERVK13-int     GGCAAAAATGGGATTTGTTCAAGGTATGGTTCTGGAAAAATCAGCACAAGGTATTACTGATACTATTATAACTACTCATAAATCTGATTCAACAGGACTT
Amplicon HeLa   ..........................................................................--------------------------
                      1210      1220      1230      1240      1250      1260      1270      1280      1290      1300
HERVK13-int     GGTTGTTCTTTTTAGAAGCAGTCACTATCAAGCCTCCAGATCCCATCCCCTTAACTTGGAAAACTCAGAAACCGGTTTGGGTAGATCAGTGGCTGCTCCC
Amplicon HeLa   ----------------------------------------------------------------------------------------------------
                      1310      1320      1330      1340      1350      1360      1370      1380      1390      1400
HERVK13-int     TAAAAATAAGCTGGCGGCACTCCATATTTTGGTCCTTGAACAATTAAAATTGGGACACATAAAGCAATCTTTTTCTCCTTGGAATTCACCTGTTTTTGTT
Amplicon HeLa   ----------------------------------------------------------------------------------------------------
                      1410      1420      1430      1440      1450      1460      1470      1480      1490      1500
HERVK13-int     ATTCAAAAGAAATCTGGTAAGTGGAAAATGCTTACTGATCTTAGGGCAGTAAATGCTGTCCTTCAACGTATGGCAACCCAACTTGCCATCCCCCACTATG
Amplicon HeLa   ----------------------------------------------------------------------------------------------------
                      1510      1520      1530      1540      1550      1560      1570
HERVK13-int     ATTCCTGAGTATTGGCCATTTATCATCATTGATCTTAAAACTGTTTCTTTAATATTCCTCTGGCCCCCTCAGGACTTT
Amplicon HeLa   ------------------------------------------------------------------------------
```

Supplementary Figure 4
