## Supplemental Figure 5 for "Unraveling viral identity: Avoiding the trap of endogenous sequences for viral surveillance of small ruminant oncogenic retroviruses"

### A. ENTV-2 *gag/ pro* (primers set #3)

```
                                    10        20        30        40        50        60        70        80        90       100
ERV II.5 Oa                 TGTGCTCGTATAGCTGATGTTAACCGACAGCAAGGTATACAGACCTCCTATGAAATGTTGATTGGTGAAGGCCCTTACCAGGCTACTGATACTCAACTTA
Amplicon IDO5               -...................................................................................................
                                   110       120       130       140       150       160       170       180       190       200
ERV II.5 Oa                 ATTTCTTACCTGGTGCATATGCACAAATATCAAATGCGGCTCGGCAGGCATGGAAAAAACTTCCTAGCTCCAGTACTAAGACAGAGGATCTTTCAAAAGT
Amplicon IDO5               ....................................................................................................
                                   210       220       230       240       250       260       270       280       290       300
ERV II.5 Oa                 CCGGCAGGGACCTGATGAGCCTTACCAGGACTTCGTGGCACGACTTTTAGATACTATAGGTAAGATAATGTCAGATGAAAAGGCTGGGATGGTATTGGCA
Amplicon IDO5               .............................................C......................................................
                                   310       320       330       340       350       360       370       380       390       400
ERV II.5 Oa                 AAACAATTGGCTTTTGAAAACGCTAACTCTGCTTGTCAAGCTGCTTTAAGACCTTATCGAAAAAAGGGAGATCTGTCTGATTTTATTCGCATTTGTGCTG
Amplicon IDO                ....................................................................................................
                                   410       420       430       440       450       460       470       480       490       500
ERV II.5 Oa                 ACATTGGACCCTCCTACATGCAAGGCATTGCTATGGCAGCAGCATTACAAGGAAAAAGCATNAAAGAGGTACTTTTCCAGCAGCAAGCCCGGAACAAGAA
Amplicon IDO5               .............................................................A..............T.......................
                                   510       520       530       540       550       560       570       580       590       600
ERV II.5 Oa                 AGGACTTCAAAAGTCAGGTAATTCGGGTTGCTTTGTTTGTGGTCAGCCTGGCCATCGGGCTGCAGTGTGCCCTCAAAAACAACAAAGCCCTGTTAACACT
Amplicon IDO5               ....................................................................................................
                                   610       620       630       640       650       660       670       680       690       700
ERV II.5 Oa                 CCTAATTTGTGCCCACGCTGTAAAAAAGGAAAGCATTGGGCGCGGGATTGCCGTTCCAAAACGGATGTTCAAGGTAATCCTTTGCCCCCGGTTTCGGGAA
Amplicon IDO5               ....................................................................................................
                                   710       720       730       740       750       760       770       780       790       800
ERV II.5 Oa                 ACTGGGTGAGGGGCCAGCCCCTGGCCCCGAAACAATGTTATGGGGCAACACTGCAGGTTCCAAAAGGACCATTGCAGACCTCTGTCGAGCCACAAGAGGC
Amplicon IDO5               ....................................................................................................
                                   810       820       830       840       850       860       870       880       890       900
ERV II.5 Oa                 AGCGCGGGATTGGACCTCTGTGCCACCTCCTACACAGTATTAACTCCCGAGATGGGGGTCCAAACCCTTGCCACAGGAGTGTTTGGGCCTTTACCTCCAG
PCR03                       ....................................................................................................
                                   910       920       930       940       950       960       970       980       990      1000
ERV II.5 Oa                 GGACAGCTGGACTGCTTTTAGGGCGCAGCAGTGCGTCTTTAAAAGGAATACTTATTCATCCTGGTGTGATTGACTCTGATTATACAGGAGAGATAAAAAT
Amplicon IDO5               ....................................................................................................
                                  1010      1020      1030      1040      1050      1060      1070       1080      1090
ERV II.5 Oa                 ATTAGCCTCCGCTCCTAACAAAATTATTGTAATCAATGCAGGACAGCGTATAGCTCAACTTCTTTTAGTT-CCATTAGTCATACAGGGAAAAACAA
Amplicon IDO5               ..............................G.......................................C..............A..........
```

### B. ENTV-2 *gag/ pol* (primers set #4)

```
                                    10        20        30        40        50        60        70        80        90       100
ERV II.5 Oa                 GATGTTCAAGGTAATCCTTTGCCCCCGGTTTCGGGAAACTGGGTGAGGGGCCAGCCCCTGGCCCCGAAACAATGTTATGGGGCAACACTGCAGGTTCCAA
Amplicon IDO5               ....................................................................................................
                                   110       120       130       140       150       160       170       180       190       200
ERV II.5 Oa                 AAGGACCATTGCAGACCTCTGTCGAGCCACAAGAGGCAGCGCGGGATTGGACCTCTGTGCCACCTCCTACACAGTATTAACTCCCGAGATGGGGGTCCAA
Amplicon IDO5               ....................................................................................................
                                   210       220       230       240       250       260       270       280       290       300
ERV II.5 Oa                 ACCCTTGCCACAGGAGTGTTTGGGCCTTTACCTCCAGGGACAGCTGGACTGCTTTTAGGGCGCAGCAGTGCGTCTTTAAAAGGAATACTTATTCATCCTG
Amplicon IDO5               ....................................................................................................
                                   310       320       330       340       350       360       370       380       390       400
ERV II.5 Oa                 GTGTGATTGACTCTGATTATACAGGAGAGATAAAAATATTAGCCTCCGCTCCTAACAAAATTATTGTAATCAATGCAGGACAGCGTATAGCTCAACTTCT
Amplicon IDO5               ...................................................................G................................
                                   410       420       430       440       450       460       470       480       490       500
ERV II.5 Oa                 TTTAGTTCCATTAGTCATACAGGGAAAAACAATTAACCGAGACCGTCAAGATAAAGGTTTCGGGTCCTCTGACGCCTATTGGGTGCAAAATGTTACCGAG
Amplicon IDO5               .....................A..............................................................................
                                   510       520       530       540       550       560       570       580       590       600
ERV II.5 Oa                 GCACGACCAGAACTTGAGCTACGCATTAATGGTAAGCTTTTCCGCGGAGTGCTTGATACAGGGGCCGATATTAGTGTTATTTCTGATAAATATTGGCCTA
Amplicon IDO5               ....................................................................................................
                                   610       620       630       640       650       660
ERV II.5 Oa                 CTACATGGCCAAAACAGATGGCTATTTCCACTCTCCAGGGTATTGGCCAAACTACCAATCCAGAACAGA
Amplicon IDO5               ...................................................................A.
```

Supplementary Figure 5

### C. ENTV2 *pro/pol* (primers set #5)

```
                                      10        20        30        40        50        60        70        80        90       100
ERV II.5 Oa                   GGCAGCGCGGGATTGGACCTCTGTGCCACCTCCTACACAGTATTAACTCCCGAGATGGGGGTCCAAACCCTTGCCACAGGAGTGTTTGGGCCTTTACCTC
Amplicon IDO5                 ....................................................................................................
                                     110       120       130       140       150       160       170       180       190       200
ERV II.5 Oa                   CAGGGACAGCTGGACTGCTTTTAGGGCGCAGCAGTGCGTCTTTAAAAGGAATACTTATTCATCCTGGTGTGATTGACTCTGATTATACAGGAGAGATAAA
Amplicon IDO5                 ....................................................................................................
                                     210       220       230       240       250       260       270       280       290       300
ERV II.5 Oa                   AATATTAGCCTCCGCTCCTAACAAAATTATTGTAATCAATGCAGGACAGCGTATAGCTCAACTTCTTTTAGTTCCATTAGTCATACAGGGAAAAACAATT
Amplicon IDO5                 ....................................................................................................
                                     310       320       330       340       350       360       370       380       390       400
ERV II.5 Oa                   AACCGAGACCGTCAAGATAAAGGTTTCGGGTCCTCTGACGCCTATTGGGTGCAAAATGTTACCGAGGCACGACCAGAACTTGAGCTACGCATTAATGGTA
Amplicon IDO5                 ....................................................................................................
                                     410       420       430       440       450       460       470       480       490       500
ERV II.5 Oa                   AGCTTTTCCGCGGAGTGCTTGATACAGGGGCCGATATTAGTGTTATTTCTGATAAATATTGGCCTACTACATGGCCAAAACAGATGGCTATTTCCACTCT
Amplicon IDO5                 ....................................................................................................
                                     510       520       530       540       550       560       570       580       590       600
ERV II.5 Oa                   CCAGGGTATTGGCCAAACTACCAATCCAGAACAGAGTTCGTCCCTTCTTACTTGGAAGGATAAAGATGGACATACAGGCCAATTTAAACCTTATATTCTG
Amplicon IDO5                 .......................................A............................................................
                                     610       620       630       640       650       660       670       680       690       700
ERV II.5 Oa                   CCCCATCTTCCAGTTAATCTATGGGGGCGTGATATATTAAGCAAAATGGGTGTTTATTTATATAGTCCTTCACCCACTGTGACAGATTTGATGTTAGATC
Amplicon IDO5                 ...T................................................................................................
                                     710       720       730       740       750       760       770       780       790       800
ERV II.5 Oa                   AGGGCTTACTTCCAAATCAAGGTTTAGGTAAACAACATCAAGGCATCATTTTGCCCCTTGATTTAAAATCTAATGAAGATCGAAAAGGCTTGGGGTGTTT
Amplicon IDO5                 ..........................................................................C.........................
                                     810       820       830       840       850       860       870       880       890       900
ERV II.5 Oa                   TCCCTAGGGACCTCTGNTTCTCCTGTGACGCATGCCGATCCTATTG-TTGGAAATCTGAGGAACCGGTATGGGTCGATCAGTGGCCCCTAACACAGGAAA
Amplicon IDO5                 ................A............A................A.....................................................
                                     910       920       930       940       950       960       970       980       990      1000
ERV II.5 Oa                   AACTTTCTGCCGCACAACAGCTGGTGCAGGAACAGCTGAGACTTGGGCATATTGAACCCTCTACCTCTGCTTGGAATTCCCCAATTTTTGTTATTAAAAA
Amplicon IDO5                 ....................................................................................................
                                    1010      1020      1030      1040      1050      1060      1070      1080      1090      1100
ERV II.5 Oa                   GAAGTCTGGGAAATGGAGATTGCTACAAGATCTTCGTAAGGTAAATGAAACAATGATGCATATGGGAGCCCTACAACCTGGGTTGCCCACTCCTTCCGCT
Amplicon IDO5                 ...................................................................................................-
```

### D. ENTV2 *pol* (primers set #6)

```
                                      10        20        30        40        50        60        70        80        90       100
ERV II.5 Oa                   TTCTTACTAAACAAGTGTTTTTCCAATCAGCTATTGATGCAGCCCGAAAATCCCATGATTTACATCACCAAAATAGTCATTCTTTACGCTTGCAATTTAA
Amplicon IDO5                 ......................T...........C..........A......T...A.C.............................G..A........
                                     110       120       130       140       150       160       170       180       190       200
ERV II.5 Oa                   AATTTCCCGTGAAGCTGCACGGCAAATTGTTAAATCTTGCTCTACTTGTCCTCAATTCTTTGTTCTCCCTCAATATGGTGTCAACCCTCGAGGTTTACGC
Amplicon IDO5                 G....................A.....C..A..............C.................................A....................
                                     210       220       230       240       250       260       270       280       290       300
ERV II.5 Oa                   CCTAATCACCTCTGGCAAACAGACGTTACTCACATTCCTCAATTTGGGCGTCTTAAATATGTTCATGTCTCTATTGACACTTTTTCCAATTTTCTCATGG
Amplicon IDO5                 ........T..............T..C...........................................................TC............
                                     310       320       330       340       350       360       370       380       390       400
ERV II.5 Oa                   CCTCCCTTCACACTGGAGAATCAACACGTCACTGTATTCAACATTTGCTGTTTTGCTTTTCTACTTCAGGAATCCCACAAACCCTTAAAACAGATAATGG
Amplicon IDO5                 ..........T..............T.....................................T...............C....................
                                     410       420       430       440       450       460       470       480       490       500
ERV II.5 Oa                   ACCTGGTTATACTAGCCGTTCTTTTCAACGTTTTTGTCTTTCTTTCCAAATTCATCATAAAACAGGAATTCCTTATAATCCACAGGGACAAGGTATTGTG
Amplicon IDO5                 ........................................................................A.....C.....................
                                     510       520       530       540       550       560       570       580       590       600
ERV II.5 Oa                   GAACGAGCCCATCAACGCCTTAAACATCAATTATTAAAACAAAAAAAGGGGAATGAACTGTATAGCCCCTCACCGCATAACGCCTTAAACCATGCTCTTT
Amplicon IDO5                 ..............G..T..C.........C.........................CT......................T.....G..T..........
                                     610       620       630       640       650       660       670       680       690       700
ERV II.5 Oa                   ATGTTTTAAATTTTTTAACTTTAGACGCAGAAGGCAATTCAGCAGCCCAGCGTTTTTGGGGAGAACGATCCTCATGCAAAAAACCACTTGTACGATGGAA
Amplicon IDO5                 .....C...........................................A..................................................
                                     710       720       730       740       750
ERV II.5 Oa                   GGATCCACTTACCAATCTGTGGTATGGGCCAGACCCTGTACTAATATGGGG
Amplicon IDO5                 A.......................................T..........
```

Supplementary Figure 5

### E. ENTV2 *pol* (primers set #7)

```
10        20        30        40        50        60        70        80        90        100
ERV II.5 Oa                 TACCCAATTAGTAAAACTGCAGACTGACCATTTAAAAACTCTAAATGACTTTCAAAAACTTTTAGGAGACATTAATTGGATACGTCCTTATTTAAAATTA
Amplicon IDO5               ................T.A.............................T............C............G.........C...............
                                   110       120       130       140       150       160       170       180       190       200
ERV II.5 Oa                 CCCACTTATACCTTGCAGCCATTATTTGACATCCTTAAAGGTGACTCTGATCCTGCGTCACCCCGAACACTTTCTTTAGAAGGACGAACTGCTTTACAAT
Amplicon IDO5               .....C..C........A................................C....................................T.A..C.......
                                   210       220       230       240       250       260       270       280       290       300
ERV II.5 Oa                 CAATAGAAGAAGCTATTAGACAACAACAGATTACTTATTGTGATTACCAACGATCATGGGGTTTGTATATACTTCCTACCCCCCGAGCACCCACAGGGGT
Amplicon IDO5               ..................................................................................T.................
                                   310       320       330       340       350       360       370       380       390       400
ERV II.5 Oa                 TCTCTATCAAGATAAACCTTTGCGATGGATATATTTGTCTGCTACTCCAACTAAACATCTGCTCCCTTACTATGAACTTGTTGCAAAAATTGTAGCAAAG
Amplicon IDO5               A..T..............................C.A......................................G.......................A
                                   410       420       430       440       450       460       470       480       490       500
ERV II.5 Oa                 GGACGTCACGAGGCCATCCAATATTTTGGTATGGAACCCCC-TTCATTTGTGTTCCTTATGCTTTANGAACAACAAGATTGGCTTTTTCAATTTTCAGAT
Amplicon IDO5               .....C..T.............T..................T.........A..............-.................................
                                   510       520       530       540       550       560       570       580       590       600
ERV II.5 Oa                 AATTGGTCTATAGCTTTTGCAAATTACCCGGGACGGATTACTCATCATTACCCTTCTGATAAATTGTTACAATTTGCTAGCTCTCATGCCTTTATTTTTC
Amplicon IDO5               .................C................................T...............................T.................
                                   610       620       630       640       650       660       670       680       690       700
ERV II.5 Oa                 CAAAAATAGTTCGCCGACAACCTATTCCCGAAGCGACACTTATATTTACAGATGGATCTTCTAATGGAACTGCAGCTTTAATCATTAACCATCAAACCTA
Amplicon IDO5               ............A.........C............................................T..............T.................
                                   710       720       730       740       750       760       770       780       790       800
ERV II.5 Oa                 TTACGCACAAACCAGTTTTTCTTCTGCTCAAGTTGTGGAATTATTTGCAGTCCACCAAGCGTTGCTAACTGTACCTACTTCCTTCAATTTATTTACAGAC
Amplicon IDO5               ...T.....T........C...........GA...................T..T.....A.....G........C........................
                                   810       820       830       840       850       860       870       880       890       900
ERV II.5 Oa                 AGCTCCTATGTGGTCGGTGCCTTACAGATGATTGAAACTGTTCCAATTATCGGCACCACCTCTCCTGAAGTTCTTAACTTATTTACATTGATTCAACAGG
Amplicon IDO5               ..............................C..............G..........A................................A..........
                                   910       920       930       940       950       960       970       980       990      1000
ERV II.5 Oa                 TTCTCCATTGCCGCCAACACCCCTGTTTCTTTGGACATATTCGTGCACACTCCACCCTTCCTGGTGCCCTGGTACAAGGCAATCACACTGCGGACGTTCT
Amplicon IDO5               CC..T..CC.T......T....G...........G..............T................................C.................
                                  1010      1020      1030      1040      1050      1060      1070      1080      1090      1100
ERV II.5 Oa                 TACTAAACAAGTGTTTTTCCAATCAGCTATTGATGCAGCCCGAAAATCCCATGATTTACATCACCAAAATAGTCATTCTTTACGCTTGCAATTTAAAATT
Amplicon IDO5               .........G........T...........C.................T...A.C.............................G..A........G...
                                  1110      1120
ERV II.5 Oa                 TCCCGTGAAGCTGCACGGCAA
Amplicon IDO5               .................A...
```

### F. ENTV2 *env* **(**primers set #8)

```
                                    10        20        30        40        50        60        70        80        90       100
ERV II.5 Oa                 AAACTTTTGGCTGCTTTTGGTCATGGCAATAGTCTATATTTACAGCCCAATATTAGTGGGAGCAAATATGGTGATGTGGGAGTTACAGGATTTTTATATC
Amplicon IDO5               .........................................-....................T..................................T..
                                   110       120       130       140       150       160       170       180       190       200
ERV II.5 Oa                 CCCGAGCTTGTGTCCCTTACCCATTCATGTTGATACAAGGCCATATGGAAATAACGCTGTCATTGAATATTTATCATTTAAATTGTTCTAATTGCATACT
Amplicon IDO5               .............T.........................................A............................................
                                   210       220       230       240       250       260       270       280       290       300
ERV II.5 Oa                 TACTAATTGCATTAGAGGTGTAGCCAAAGGAGAACAAGTTATAATAGTAAAACAACCTGCTTTTGTAATGTTACCTGTTGAAATAACTGAAGAATGGTAT
Amplicon IDO5               ....................................................................................................
                                   310       320       330       340       350       360       370       380       390       400
ERV II.5 Oa                 GATGAAACTGCTTTAGAATTGTTACAACGCATTAATACGGCTCTTAGCCGTCCTAAAAGAGGTCTGAGCCTGATTATTCTGGGTATAGTGTCTTTAATCA
Amplicon IDO5               .....G..............................................................................................
                                   410       420       430       440       450       460       470       480       490       500
ERV II.5 Oa                 CCCTTATAGCAACTGCTGTTACTGCTTCTGTATCTTTAGCACAATCCATTCAAGCTGCTCATACTGTAGATTCCTTGTCATATAATGTTACTAAAGTAAT
Amplicon IDO5               ....................................................................................................
                                   510       520       530       540       550       560       570       580       590       600
ERV II.5 Oa                 GGGAACTCAAGAAGATATAGATAAAAAAATAGAAGATAGATTATCAGCTTTATATGATGTAGTTAGAGTTCTAGGAGAACAAGTTCAGAGCATTAATTTT
Amplicon IDO5               ......................................................................T.............................
                                   610       620       630       640       650       660       670       680       690       700
ERV II.5 Oa                 CGCATGAAAATTCAATGCCATGCTAATTATAAATGGATTTGTGTTACAAAAAAGCCTTACAATACTTCTGACTTTCCGTGGGATAAGGTGAAAAAACATC
Amplicon IDO5               ...................................................................T................................
                                   710       720       730       740       750       760       770       780       790       800
ERV II.5 Oa                 TGCAAGGAATTTGGTTTAATACTAATGTTTCTTTAGATCTTTTACAATTGCACAATGAAATTCTTGACATCGAAAATTCTCCAAAAGCTACTTTGAATAT
Amplicon IDO5               ....................................................................................................
                                   810       820       830       840       850       860       870       880       890       900
ERV II.5 Oa                 AGCTGATACCGTCGATAATTTTTTACAAAATTTATTTTCTAACTTTCCTAGCCTTCATTCACTGTGGCGAAGTATAATTGCTATGGGCGCGGTTCTGACT
Amplicon IDO5               ....................................................................................................
                                   910       920       930       940       950       960       970       980       990      1000
ERV II.5 Oa                 GTTGTGCTTATCATAATTTGTCTAGCTCCTTGTCTTATTCGTAGCATTGTTAAAGAATTTCTACATATGAGAGTTTTAATACATAAAAACATGTTGCAAC
Amplicon IDO5               .....................T..........C...........................T.......................................
                                  1010      1020      1030      1040        1050      1060      1070      1080      1090
ERV II.5 Oa                 ACCGACATCTTATGGAGCTTTTAAAAAATAAAGAGAGGGGA--TGCGGGGGACGACCCGTGAAGGGTTAAGTCTTGGGAGCTCCCTGGCAGGTATG
Amplicon IDO5               ...A.....................................GC.....................................................
```

Supplementary Figure 5

G. JSRV *env* (primers set #14)

```
                                      10        20        30        40        50        60        70        80        90       100
ERV II.5 Oa                   CGCATGACACTGAGCGAGCCCACGAGTGAGCTGCCTACCCAGAGGCAAATTGAGGCGCTGATGCGTTATGCTTGGAATGAGGCTCATGTACAACCTCCAG
Amplicon IDO5                 .................................................................A..................................
                                     110       120       130       140       150       160       170       180       190       200
ERV II.5 Oa                   TGACACCTACTAATATACTGATCATGTTATTATTATTGTTACAGCGGATACAAAACGGGGCGGCTGCGGCTTTTTGGGCATACATTCCTGATCCGCCTAT
Amplicon IDO5                 ....................................................................................................
                                     210       220       230       240       250       260       270       280       290       300
ERV II.5 Oa                   GATTCAATCCTTAGGATGGGATAAAGAAACAGTACCTGTATATGTTAATGATACAAGTCTTTTAGGAGGAAAATCAGATATTCACATTTCTCCTCAGCAA
Amplicon IDO5                 ....................................................................................................
                                     310       320       330       340       350       360       370       380       390       400
ERV II.5 Oa                   GCCAATATCTCCTTTTATGGTCTTACTACTCAATACCCTATGTGCTTTTCTTATCAATCACAGCATCCTCATTGTATACAGGTGTCAGCTGATATATCCT
Amplicon IDO5                 ....................................................................................................
                                     410       420       430       440       450       460       470       480       490       500
ERV II.5 Oa                   ATCCTCGAGTGACTATTTCAGGCATTGATGAAAAAACCGGAAAGAGATCGTACCGTGACGGAACCGGACCCCTCGACATTCCGTTTTGTGACAAACATTT
Amplicon IDO5                 ....................................................................................................
                                     510       520       530       540       550       560       570       580       590       600
ERV II.5 Oa                   AAGCATCGGCATAGGAATAGACACTCCTTGGACTTTATGTCGAGCACGAATTGCATCGGTGTATAACATCAACAATGCCAATACCACCCTTTTATGGGAC
Amplicon IDO5                 ....................................................................................................
                                     610       620       630       640       650       660       670       680       690       700
ERV II.5 Oa                   TGGGCACCTGGAGGAACACCTGATTTCCCCGAATATCGAGGACAGCATCCACCCATTCTCTCTGTAAACACTGCTCCTATATTTCAAACTGAACTGTGGA
Amplicon IDO5                 ..................................................................................A...G..A..........
                                     710       720       730       740       750       760       770       780       790       800
ERV II.5 Oa                   AACTTTTGGCTGCTTTTGGTCATGGCAATAGTCTATATTTACAGCCCAATATTAGTGGGAGCAAATATGGTGATGTGGGAGTTACAGGATTTTTATATCC
Amplicon IDO5                 .............................................................T......................................
                                     810       820       830       840       850       860       870       880       890       900
ERV II.5 Oa                   CCGAGCTTGTGTCCCTTACCCATTCATGTTGATACAAGGCCATATGGAAATAACGCTGTCATTGAATATTTATCATTTAAATTGTTCTAATTGCATACTT
Amplicon IDO5                 ............T.........................................A.............................................
                                     910       920       930       940       950       960       970       980       990      1000
ERV II.5 Oa                   ACTAATTGCATTAGAGGTGTAGCCAAAGGAGAACAAGTTATAATAGTAAAACAACCTGCTTTTGTAATGTTACCTGTTGAAATAACTGAAGAATGGTATG
Amplicon IDO5                 ....................................................................................................
                                    1010      1020      1030      1040      1050       1060      1070
ERV II.5 Oa                   ATGAAACTGCTTTAGAATTGTTACAACGCATTAATACGGCTCTTAGCCGTCCTAAAAG-AGGTCTGAGCCTGATT
Amplicon IDO5                 ....G.....................................................A................
```

Supplementary Figure 5
