## Supplemental Figure 6 for "Unraveling viral identity: Avoiding the trap of endogenous sequences for viral surveillance of small ruminant oncogenic retroviruses"

```
                    7310      7320      7330      7340      7350      7360      7370      7380      7390      7400
ERV II.5 Oa   CTTTCCTAGCCTTCATTCACTGTGGCGAAGTATAATTGCTATGGGCGCGGTTCTGACTGTTGTGCTTATCATAATTTGTCTAGCTCCTTGTCTTATTCGT
KT266728      ----........................AC..........G...........................AT.G.......T..........C.........
JSRV FR2054   T..C.....T......G..T.....AA..CCC.G..C.G.C.A..AATAT..G...TAA..A.TGC....G....C.T.G..TT...C...G.CG.....
JSRV FR989     T..C.....T..C.....G......AA..CCC.G....G.G.A..AATAC..G..TT.A..A.AA..G..G....CCT.A..TT......C...G.....

                                                                  H  K  X  M
                    7410      7420      7430      7440      7450      7460      7470      7480      7490      7500
ERV II.5 Oa   AGCATTGTTAAAGAATTTCTACATATGAGAGTTTTAAT------------ACATAAAAACATGTTGCAACACCGACATCTTATGGAGCTTTT---AAAAA
KT266728      ..................T..............C.G..------------..........................................AAA.....
                                                                  Y  X  X  M
JSRV FR2054   G.TC.C...CGT..C...T..A.G.........GA...GCTACATATGAA.T...G..CT.....A..............A...........---...G.
JSRV FR989     G....G...CGC..T......A.G.........GA...GCTGCATATGAA.T...G...T.....A..G....A......A...........---.....

                    7510      7520      7530      7540      7550      7560      7570      7580      7590      7600
ERV II.5 Oa   ATAAAGAGAGGGGA--TGCGGGGGACGACCCGTGAAGGGTTAAGTCTTGGGAGCTCCCTGGCAGGTATGCCGGGCCCTAGGACACGTGCCTAAGCTCCCT
KT266728      ..............GC..........TG...........................G.TC...GT--TATG.A.AG...TA.G..T..T............
JSRV FR2054   ..............GC..............................C.........TT.....A--.A...AAAG.T.......AA.A............
JSRV FR989     ..............GA..............................C.........TT.....R--.A...AAAG.........A..A............

                    7610      7620      7630      7640      7650      7660      7670      7680      7690      7700
ERV II.5 Oa   GTCCCGCCACCCTCAAGAGTTTTTATAA-CCCTTAAGGCTCCAAGATGTTTGGTTTCGGCAACATTTCATAGAA--GATAGATTATCTTATTGTGTATAT
KT266728      ........C....G....A...A.TA---..............G.C.........CTT................--.................-------
JSRV FR2054   ..............T..GA..C..GA..G.T.......T..--G........C.GCT...TT.C.........GAT..CG.GAA....G...A...----
JSRV FR989     ..................A.....A.GG.T...........--G........C..CT...CTGC..........ATACC..GAA....G...A.A.----

                    7710      7720      7730      7740      7750      7760      7770      7780      7790      7800
ERV II.5 Oa   TTCATAGAAGATAGATATTCTGATTGTGTTCTGTATACAATGGTAAGGGTCTGGTGATTGTATCCTGAGATTAAAAAACAACCTTGTGAGTGCCTTAAGT
KT266728      ---.............TA...T......A......C................T..........TTG........G..CA-.T.......A..T.AG....
JSRV FR2054   ----------.AGA..CCGG......----------------....AAA..C....GG....GTT......G..T....-----------------.G..
JSRV FR989     ----------.AGA..CCGG......----------------.....AA..C.....G....GTGAATA..G..T....-----------------....

                    7810      7820      7830      7840      7850      7860      7870      7880      7890      7900
ERV II.5 Oa   CACGTACTTTACCCTATATATACCGCAGCACAATAAAGCAAGGTATCAGCCATTTTGGTCTGATCCTCTCAACCCCATCTTTTGTCTATCTCTTATTTTC
KT266728      ..........T.........T............G...C.................................................C...T-AT.....
JSRV FR2054   T.T...A.G...TG....A...TA...A.GT...................GC.C.................................T....--.G....
JSRV FR989     T.T..TG.A...TT....A...TA...TTGT.........GA.............................................C............

                    7910      7920      7930      7940      7950      7960
ERV II.5 Oa   TTAGCGGGGACGCTCCGTTCTCTCCCTGTGCAGGTGCGACT-CTTGCTTGTGCTGGCCGCGGCA
KT266728      .....................................T...A....A...........------
JSRV FR2054   .........................................-....T.A.......T.......
JSRV FR989     .....................................T...-........C.............
```

Supplementary Figure 6
