## Supplemental Figure 7 for "Unraveling viral identity: Avoiding the trap of endogenous sequences for viral surveillance of small ruminant oncogenic retroviruses"

## A. DR1

```
                600       610       620       630       640       650       660       670       680       690       700
ERV II.5 Ch    GTTCCTGATCCGGAACCCAGGTATGCTGTTCCCAAGGGAGTTGAAGGCGACCCTCCGTTTTCTAAATTATCGCGTCCTTCAGATAATGATGATTCACTTTC
ERV II.5 Oa    .........T.........A...........G....G...A..A...................C....T.........G.............T......   ERV
TickP118       .........T..........................................................................................

ENTV1 FR4403    .A......CT.....A.T..T....A..C...T---------CCCCCTCCT.A.....CC.TAC.T.CG........C................T......
ENTV2 FR3824    A.......C..A..G..TG.T....A..C...T------------CCTCCT.....TCCG..CCTGAA..T...A........................C.   XRV
JSRV FR2054     ACC......T...T...T..T....A.CC...T------------CCTCCT......CCA...CTGAA.AT..AC.....T........C..........
JSRV FR989      AC...C...T...G...T..T....A.CC...T------------CCTCCT.....CCCA...CTGAA.AT..AC...................T......
```

## B. DR2

```
                    860       870       880       890       900       910       920       930       940       950       960       970       980
ERV II.5 Ch    AAGCTTCCCATTAGCAACTCTAAATCTCCTTCTGCTCCGCCTCTGCCCCCTGCCTATGCTCCTTCCATTATAGCAGGTCTCGATCCCCCTCCGGGGCCTCCTCCTCCTCCTCCTCCATCTGAGATT
ERV II.5 Oa    ........T.G..A...TG.C......................T........T...........TG..G.G..T.................A...---------..C.....A..G.......AC   ERV
TickP118       ...............................................................................................................................

ENTV1 FR4403    ..A.....T.C..AT..TGA....CT.T................T..T.....T.....C.....TG..G.C.......................---------..C.....A..G...A.A.A.
ENTV2 FR3824    ...T..T.T.A..ATC..GTC...C....C...........AT.C..A.....T..C........TG..G.G..C.....T..........A...---------...T..TTA..GA.......C   XRV
JSRV FR2054     C.......T.G..AT...AA..CC.T...C.............T.T..T.....T...A....CA.TG....G..T..C..T..........C...---------TT......G..G...A.ACAC
JSRV FR989      TC......TCA..AT..---..CC.T...C.............T.T..T..C..T...A.......TG....G..T..C..T........G..C...---------TT......A..G...A.ACA.
```

## C. DR3

```
                  7020      7030      7040      7050      7060      7070      7080      7090      7100
ERV II.5 Ch    CGTAGCATAATTGCTGTGGGCGCGGTTCTGACTGTTGTGCTTATATTGATTTGTTTAGCTCCTTGCCTTATTCGTAGCATTGTT
ERV II.5 Oa    ..A..T.........A..........................CA.A......C..........T..................   ERV
TickP118       ....................................................................................

ENTV1 FR4403    .AA..T..T....T.A.A.CTATAA..G..TT.A.....A....TG.A...CT.....T.........C.C...AG.TT.AA..
ENTV2 FR3824    .AG..T...C........ATTATAA..G...TC...A.AA....TG.A..........T.........CC.....G.TC.CT..   XRV
JSRV FR2054     AAA.C.C.G..C.G.C.A..AATAT..G...TAA..A.TGC...CG.A..C.T.G..TT...C..TG.CG.....G.TC.C...
JSRV FR989      AAA.C.C.G....G...A..AATAC..G..TT.A..A.AA..G.CG.A..CCT.A..TT..........G.....G....G...
```

```
                        7110      7120      7130      7140      7150      7160      7170      7180
                                                        H  K  N  M
ERV II.5 Ch    AAAGAATTTTTACATATGAGAGT------------TCTGATACATAAAAACATGTTGCAACACCGACATCTTATGGAGCTTTTAAAA
ERV II.5 Oa    .........C.............------------.T.A.............................................-...   ERV
TickP118       .........C....G........------------.T.A.............................................-...
                                                        Y  X  X  M
ENTV1 FR4403    .....T..CC.G..A......A.TGAATTAATACA.A...A.T...G.T.............T...A.A.A.........T...G.---G
ENTV2 FR3824    CG...T..CC............CTGAATTGCTACA.T...A.T...G.C.T...A.....T.......G..A...........-.GG   XRV
JSRV FR2054     CGT..C......A.G........TGAAATGCTACA.A...A.T...G..CT.....A..............A...........-...
JSRV FR989      CGC..T...C..A.G........TGAAATGCTGCA.A...A.T...G...T.....A..G....A......A...........-...
```

## D. DR4

```
                    7230      7240      7250      7260      7270      7280      7290      7300      7310      7320      7330      7340
ERV II.5 Ch    TGAAGGGTTAAGTCTTGGGAGCTGCTCGGC-GTTATGCAGAGCCCTAGGACACGTTCCTAAGCTCCCTGTCCCGCCCCCCTGAAGAA--TTTATTA-CCCTTAAGGCTCCGGGACGTCTGGTTCTT
ERV II.5 Oa    .......................C.CT...A.G.....C.G..............G.....................A....C....GTT.....A.-.............AA..T..T.....TCG   ERV
TickP118       .............................-............T.......T................................TT.......-............A......T........

ENTV1 FR4403    C.G...........C........C..T...-AAA..A.CAG.G......G..A..A.....................A....C.G...GTC...AG.G.TT..GG.....A.ATGTT.T.T.G.T..
ENTV2 FR3824    ..............C.....C..C..T...--AA....CAG.A...T..GA.A..A.....A...............A....C.G...GTC...AA.G.T....C.....--AA.T.ATC...GT.G   XRV
JSRV FR2054     ..............C........CT.T...--AA.A..CA.AG.T.......AA.A.....................A....CT..G.TTC..GAA.G.T.......T..--...T..T..C.G..G
JSRV FR989      ..............C........CT.T...--AR.A..CA.AG.........A..A.....................A....C.....TT....AAGG.T..........--...T..T..C....G

                 7350      7360      7370      7380      7390      7400      7410      7420      7430      7440      7450      7460      7470
ERV II.5 Ch    GCAACATTTCATAGAA---GATAGATTAGCTTATTG----------ATAGAAGATAGATTATCTATTTGTGATCTGTACACAACGGTAAGGGTCTTGTGATTGTATT-TGGAGATTAA-GAACAAT
ERV II.5 Oa    ................---.........T.......TGTATATTTC.............AT...GA.....T......T....T...........G..........C-CT........AA.....C   ERV
TickP118       ................---.................---------------------------------.............T.....................-..........-.......

ENTV1 FR4403    .......GG.T..A..----GC..GAA.T..G...ATGT----------------------------------------------A.GAAAA.CA..A.....---------G.A.GTA..TG..
ENTV2 FR3824    ...TA.C.........AAGAGC..GAA.T..G...ACAT-----------------------------------------------....A..CG..........C.AA.A..A.GCAG.T..G   XRV
JSRV FR2054     ...TT.C.........GA-T..CG.GAA.T..G...ATGT--------------.AGA..CCGG.GA....---------------....AAA..CG...GG....G.-.T.....G..TA...---
JSRV FR989      ...CTGC.........A-TACC..GAA.T..G...ATAT--------------.AGA..CCGG.GA....---------------.....AA..CG....G....G.-GAATA..G..TA...---

                   7480      7490      7500      7510      7520
ERV II.5 Ch    CTTGTGAATGTCAGAAGTCACGTACTTTATCCTATATATACTGCAGCAC
ERV II.5 Oa    .......G..C.TT...............C...........C.......   ERV
TickP118       .................................................

ENTV1 FR4403    ...-----------....T..C.GA...GC-.............TA...
ENTV2 FR3824    A.C-----------..........TAA..CT...........ATG....   XRV
JSRV FR2054     --------------.G..T.T...A.G..CTG....A...TA...A.GT
JSRV FR989      --------------....T.T..TG.A..CTT....A...TA...TTGT
```

Supplementary Figure 7

E. DR3-DR4

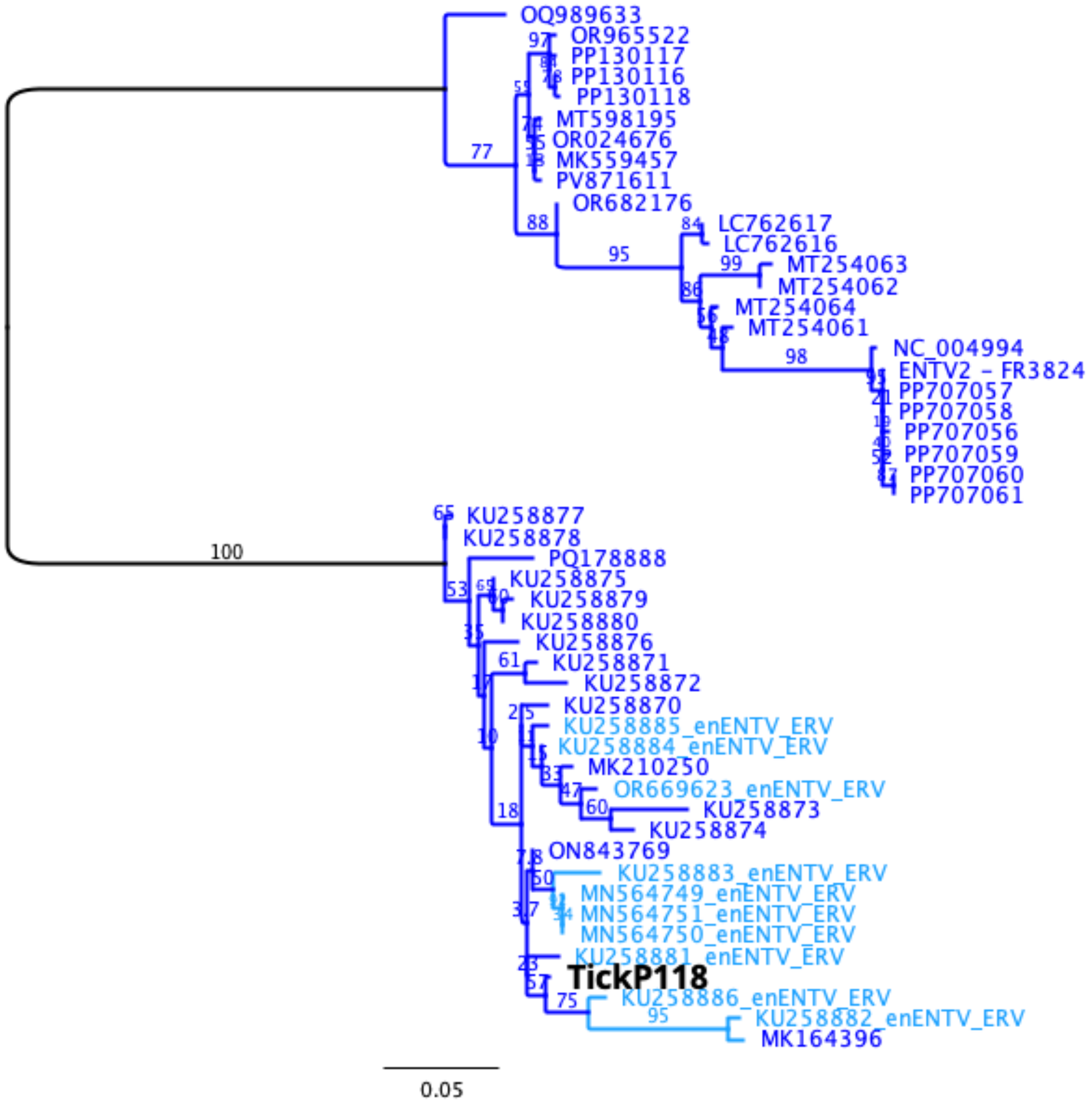


Supplementary Figure 7
